# *IN VIVO* ISOLATION OF A QUIESCENT MELANOMA POPULATION WITH INVASIVE PROPERTIES UNVEILS A TRANSCRIPTIONAL REPROGRAMMING DRIVEN BY THE TUMOR NICHE

**DOI:** 10.1101/2023.07.31.551214

**Authors:** Lotti Fiorenza, Meliksetyan Marine, Malferrari Marco, Quaresima Nicolò, Rapino Stefania, Mollo Velia, Ferrarotto Ilaria, Vlachou Thalia, Bossi Daniela, Pelicci Pier Giuseppe, Luzi Lucilla, Lanfrancone Luisa

## Abstract

Melanoma is a heterogeneous tumor composed of many interacting cellular populations and highly plastic melanoma cells that pass through distinct cell states to adapt to the surrounding microenvironment. Slow cycling is a transient state that defines a minor population of cells with cancer-initiating features. These cells are enriched upon drug therapy and can trigger cancer relapse and metastasis dissemination when they acquire proliferative potential. This population is still not entirely characterized.

Here we provide evidence of the existence of a slow cycling melanoma population isolated *in vivo* from melanoma PDXs using the H2B-GFP system. These cells display a highly invasive phenotype and are able to dynamically respond to cancer microenvironmental stimuli. Single cell transcriptomic analysis unveils a significant transcriptional heterogeneity of GFP-retaining slow cycling cells, defining a quiescent subpopulation of cells. These cells show a different phenotype in primary tumors and matched metastases, suggesting that tumor niche pressure drives a transcriptional reprogramming of quiescent cells during melanoma progression.

## INTRODUCTION

Metastatic spreading represents the most frequent cause of death for melanoma patients and the major challenge for oncologists (1). Metastatic dissemination is favored by melanoma heterogeneity and plasticity. Indeed, melanoma displays a phenotypical and functional heterogeneity that allows melanoma cells to sense the tumor microenvironment (TME) and progressively adapt to it developing new mechanisms of survival and resistance (2–8).

Melanoma plasticity is mainly based on the capability of the cells to switch between proliferative and invasive phenotypes during tumor development and metastasis formation and it has been described in preclinical models and clinical samples (3,9,10). Recently, different functional cell states with unique transcriptional profiles and tumor spatial organization were dissected in melanoma (5,11–14). The modulation of these cell states within the tumor is mainly due to altered interaction and secretory activities among cells and TME, and it contributes to the emergence of different drug-tolerant states that impact on therapeutic sensitivity, preventing control of the local disease and causing clinical failure (12). Even if extensively investigated, the biological understanding of all the different populations composing the melanoma has yet to be dissected.

Among the different phenotypes, slow cycling (quiescent) cells, which account for a small subpopulation within the tumor, are characterized by a non-proliferative state with a reversible cell cycle arrest and have a pivotal role in driving tumor relapse and drug resistance (15–20). Slow cycling cells were reported in patients (21) and they have been described with some biological features in common with cancer stem cells, such as mediating invasion/metastasis and resistance to treatments, but also for their ability to undergo a dynamically reprogramming in response to microenvironmental cues (10,22–27). A variety of biomarkers, such as CD133 (28), aldehyde dehydrogenase 1 (ALDH1) (29), CD271 (30,31), ABCB5 (8), CD44 (32,33), lysine demethylase 5B (JARID1B) (15) and SOX10 (34) have been proposed for their isolation but none of them have been shown to univocally identify this population in the tumor (35,36). The technical challenges somehow limited their complete characterization.

In the current study, we have labelled and isolated slow cycling cells from melanoma patient derived xenografts (PDXs) *in vivo* using the H2B-GFP expression system. We have characterized their functional properties *in vitro* and *in vivo*, demonstrating their ability to invade, to respond and adapt to TME stimuli, such as changes in oxygen levels, and to resist to drug treatment. Single cell transcriptomic analysis has shown a significant transcriptional heterogeneity of GFP-retaining slow cycling cells, allowing the identification of a subpopulation of quiescent GFP-positive and KI67-negative cells. Notably, these quiescent cells display a different transcriptional profile in primary tumors versus matched metastatic lesions, suggesting that the metastatic niche drives a transcriptional reprogramming of quiescent melanoma cells.

Further studies are required to understand the role of quiescent cells in the metastatic process and to fully elucidate the quiescence regulators driven by the cancer niche in order to design more effective therapeutic strategies.

## RESULTS

### *In vivo* isolation of slow cycling cells from metastatic melanoma PDXs

To directly visualize and isolate melanoma slow cycling cells *in vivo*, we took advantage of the doxycycline-repressible histone H2B-GFP green fluorescent protein reporter system stably incorporated into the DNA (Supp. Fig. 1A). Once doxycycline is added, it suppresses the expression of H2B-GFP, allowing the serial dilution of the GFP signal in proliferating cells and its retention into slow cycling cells (18,37). Melanoma cells of two PDXs carrying NRAS (MM13) or BRAF (MM27) mutations, were transduced with the H2B-GFP-containing lentiviral vector and then serially transplanted into NSG mice, sorting the GFP expressing population at each passage (Supp. Fig. 1B). The purified GFP-sorted melanoma cells were injected in NSG mice and tumors were grown for about two weeks until palpable (Fig. 1A upper panel; Supp. Fig. 1C upper panel) before doxycycline was added for additional three weeks. GFP-negative (GFP-), proliferating cells that have been dividing during the chasing period can be neatly separated over time from the GFP-positive (GFP+), label retaining cells that are slow cycling and represent only a small fraction of the tumor population (around 0.2-1%, Fig. 1A upper panel; Supp. Fig. 1C upper panel; Supp. Fig. 1D), as expected (18). The GFP+ cell population is gradually reduced upon doxycycline treatment as a consequence of GFP decrease at each cell division, as shown by the histograms (Fig. 1A lower panel; Supp. Fig. 1C lower panel). We confirmed the slow cycling features of the GFP+ population by an *in vivo* 10 days long pulse with 5-bromo-2’-deoxyuridine (BrdU), enough for being incorporated in all the cells, and a subsequent simultaneous chase of BrdU, and GFP suppression by doxycycline treatment. GFP+ cells were indeed 100% BrdU-positive (BrdU+) while GFP- cells completely lost BrdU expression (Fig. 1B). Conversely, a short BrdU pulse (24h) of freshly sorted GFP+ and GFP- cells, showed no BrdU incorporation in GFP-retaining cells compared to proliferative cells, confirming their slow cycling state (Fig. 1C; Supp. Fig. 1E).

**Figure 1.**
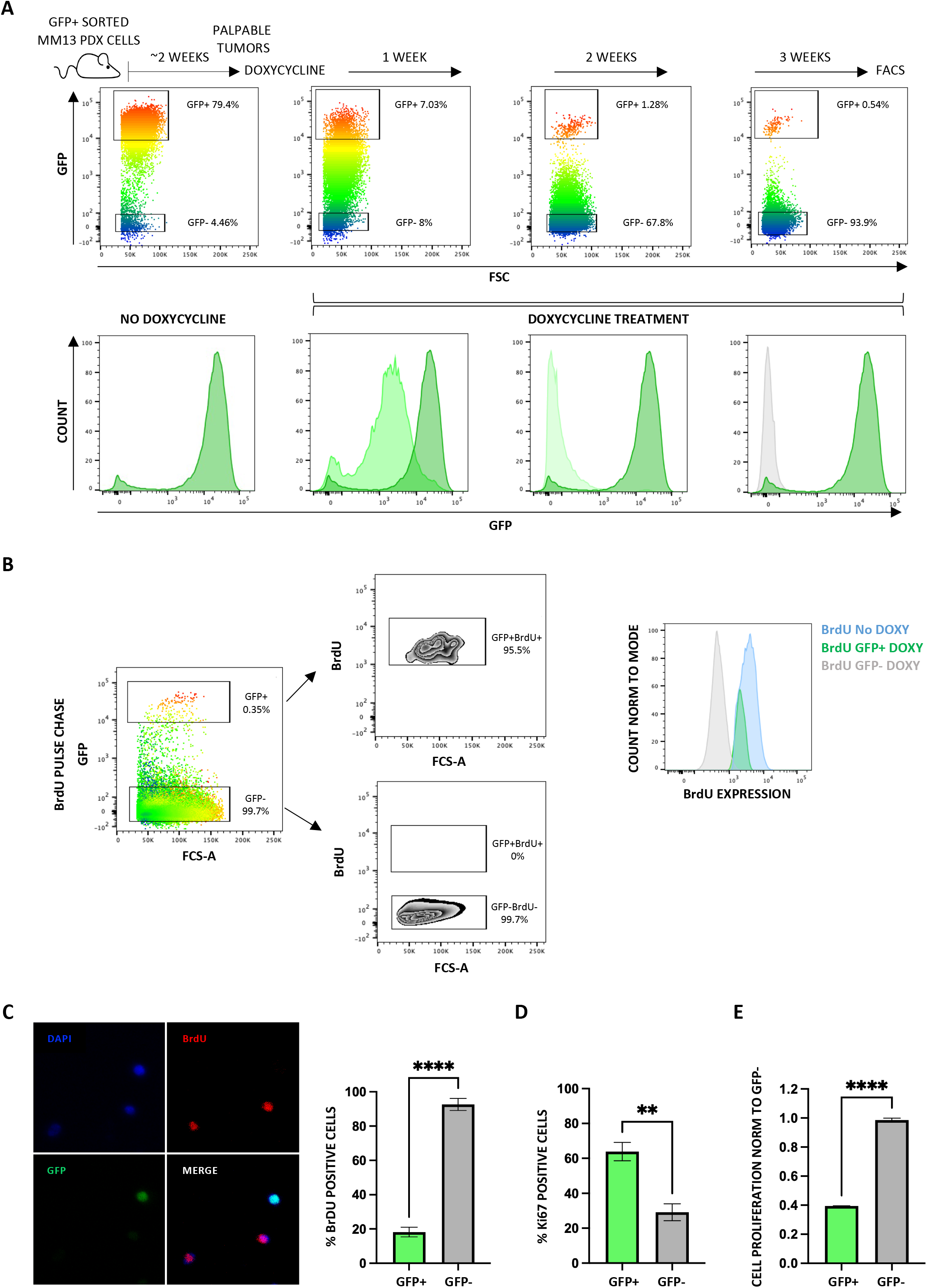
Isolation of slow cycling MM13 PDX cells *in vivo* by label-retaining H2B-GFP tet-off vector expression. **A)** H2B-GFP MM13 cells were chased *in vivo* with doxycycline. Representative dot-plot flow cytometry analysis (upper panel) and distribution histograms of GFP expressing cells (lower panel) showed the cell proliferation (gradual GFP loss) over time with around 0.5% of the cells retaining the maximum label after 3 weeks of chasing. Not treated cells represent the positive control (no doxycycline). **B)** H2B-GFP MM13 cells were pulsed with BrdU *in vivo* (n=4) for 10 days and then chased for BrdU and GFP suppression. Representative (1 mouse) flow cytometry analysis shows BrdU expression within GFP+ and GFP- populations. Histograms report BrdU expression after labelling in not treated positive control cells (blue peak) and diluted expression after chasing in BrdU/GFP+ (green peak) and BrdU/GFP- (gray peak) cells. **C)** GFP+ and GFP- MM13 cells were pulsed with BrdU for 24h *in vitro*. Cells were fixed and analyzed by confocal microscopy (representative images) and counted for GFP and BrdU expression by ImageJ. Data are presented as mean ± SD (n=5). Student t test was applied to assess the significance (****pvalue<0.00001). The experiment was performed twice. **D)** GFP+ and GFP- MM13 cells were stained with KI67 and analyzed by flow cytometry. Data are presented as mean ± SD (n=3). Student t test was used (**pvalue<0.01). The experiment was performed three times. **E)** GFP+ MM13 cell proliferation was assessed by CyQuant and normalized to GFP- cells, as mean ± SD (n = 3). p-values are based on unpaired Student’s t test (****p < 0.00001). The experiment was performed three times.

To better characterize our slow cycling GFP+ population, we assessed the expression of the proliferation marker MKI67 (KI67), whose absence is indicative of entry into a quiescent cell state in many cancers (18,26,38). Interestingly, around 30-35% of GFP+ cells were KI67 positive (KI67+, Fig. 1D; Supp. Fig. 1F), indicating that our population is composed of cells with different levels of quiescence/proliferation. Indeed, it has been previously reported that KI67 levels decrease as a function of time during quiescence entry, and that quiescent cells re-entering the cell cycle display heterogeneous levels of KI67 depending on how long the cells have been quiescent, thus suggesting that GFP+KI67+ cells can be either entering or exiting the quiescent state (39–41). However, the GFP- population clearly shows an increased proliferation rate compared to the GFP+ population, despite the presence of a proliferative component (KI67+) within the population (Fig. 1E; Supp. Fig. 1G).

### PDX slow cycling melanoma cells display high invasive potential, reduced drug sensitivity and increased oxidative phosphorylation *in vitro*

Some features of slow cycling cells, such as enhanced capability to migrate, invade and tolerate treatments, are shared among different solid tumors (25,42) and melanoma as well (15–17). We therefore characterized the functional properties of GFP+ and GFP– cells, isolated by cell sorting from MM13 and MM27 PDXs. An *in vitro* transwell migration assay showed that GFP+ cells have an increased migration capacity than GFP- cells (Fig. 2A; Supp. Fig. 2A). We then investigated the invasive properties of MM13 and MM27 GFP+ and GFP– populations as 3D spheroids embedded in a collagen matrix. As shown in Fig. 2B and Supp. Fig. 2B, both populations were capable of invading the matrix in 48 hours, being the GFP+ cells significantly more invasive than the GFP- cells. We next tested the sensitivity of MM13 and MM27 GFP+ and GFP- cells to the MEK inhibitor trametinib. Indeed, GFP+ cells showed a significant more resistant phenotype in both PDXs (Fig. 2C; Supp. Fig. 2C).

**Figure 2.**
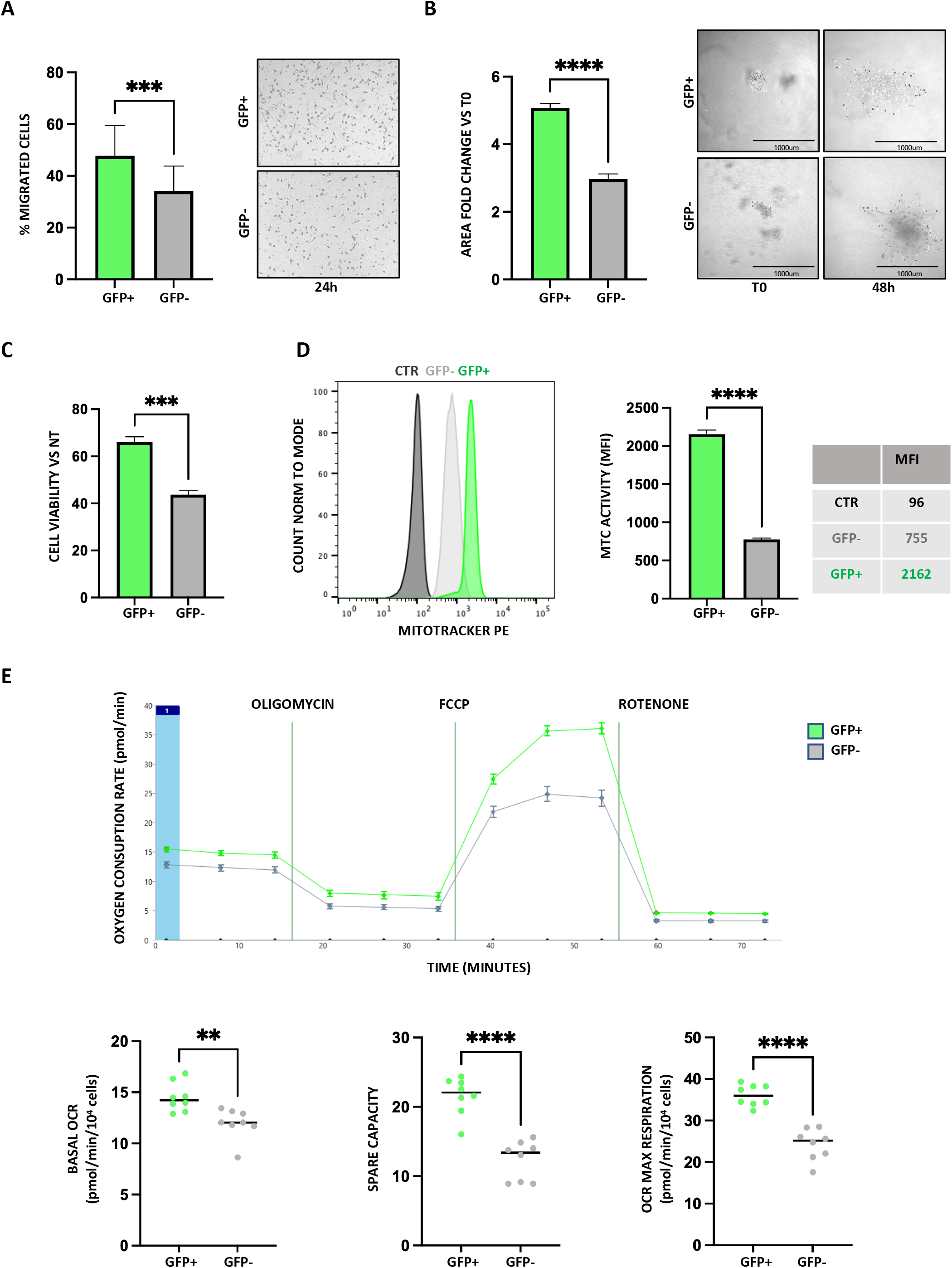
*In vitro* functional characterization of MM13 slow cycling cells. **A)** GFP+ and GFP- MM13 cell migration was assessed by transwell migration assay at 24h. Representative EVOS microscopy images of migrated cells (transwell outer surface) are reported. Data are mean+-SD (n=3). p-values are based on unpaired Student’s t test (***p < 0.0001). The experiment was performed three times. **B)** Spheroid collagen invasion area in GFP+ and GFP- MM13 cells measured as fold change vs T0 (imaged by EVOS microscopy every 24h for 48h). Data are shown as the mean ± SD of 10 different spheroids per group. Student t-test (****pvalue<0.00001). Representative images are shown. **C)** Treatment of MM13 GFP+ and GFP- cells with 10nM trametinib for 72 hours *in vitro*. Cell viability was assessed by CyQuant and normalized to DMSO control. Mean ± SD (n=3). p values are based on unpaired Student’s t test (***p < 0.001). The experiment was performed three times. **D)** GFP+ and GFP- MM13 cell mitochondrial activity was assessed by flow cytometry analysis of mitotracker orange (25nM) stained cells. Data are shown as mean fluorescence intensity ± SD (n=3). Student t-test (****pvalue<0.00001). The experiment was done twice. **E)** Basal oxygen consumption rate (OCR), OCR spare capacity and OCR max respiration of GFP+ and GFP- MM13 cells were assessed by Seahorse XF Cell Mito Stress. Data are mean ± SD (n=7). Student t-test (**pvalue<0.01; ****pvalue<0.00001). The experiment was performed twice.

Together our data show that the slow cycling GFP+ cells display increased migration, invasion and drug resistance features, suggesting that this population more likely escapes the primary tumor and promotes tumor recurrence.

Since it has been shown that different tumor subpopulations may rely on distinct metabolic pathways for their growth and survival, and that slow cycling cells preferentially use mitochondrial oxidative phosphorylation (20,22,43,44), we explored the metabolic landscape of our melanoma slow cycling populations. We compared the mitochondrial respiratory activities of freshly isolated GFP+ and GFP– MM13 and MM27 cells. We showed an increased mitochondrial functionality by detection of mitochondrial respiration (assessed by MitoTracker Orange), coupled with no substantial mitochondrial mass changes (assessed by MitoTracker Red) in GFP+ cells (Fig. 2D; Supp. Fig. 2D). In addition, oxygen consumption rate (OCR), measured by Seahorse, showed that the basal and maximal oxygen consumption rates, as well as ATP production were significantly higher in GFP+ cells. Taken together, these data demonstrated that GFP+ cells displayed higher OXPHOS and mitochondrial activities compared to GFP- cells, suggesting that the increased metabolic capacity of slow cycling cells allows a better adaptation to microenvironmental changes and an increased metastatic potential.

### PDX slow cycling cells showed an increased ability to give rise to melanoma and to metastasize *in vivo*

To check if the functional properties characterizing GFP+ cells *in vitro* were maintained in an *in vivo* setting, we intradermally injected 10,000 GFP+ and GFP- MM13 and MM27 populations in NSG mice and monitored primary tumor growth and, upon resection, metastatic potential. Even though the tumor initiation capability was comparable in both PDXs (Fig. 3A; Supp. Fig. 3A), we found that the GFP- population grew more rapidly, showing a shorter latency to palpability and to tumor resection (performed at a fixed volume of ∼0.3cm^3^) (Fig. 3B-C; Supp. Fig. 3B-C-H). These features can be explained by their proliferative phenotype, which inversely correlate with the invasive capacities, as shown by the increased latency to lymph-node invasion (Fig. 3D; Supp. Fig. 3D-H), in line with our *in vitro* observations in collagen invasion assays. Moreover, after mice sacrifice at day 100 post injection, even though both populations were able to metastasize to lymph-nodes and distant organs (Fig. 3E; Supp. Fig. 3E), GFP+ cells disseminated more efficiently showing increased lymph-node volumes (Fig. 3F; Supp. Fig. 3F) and metastases size, measured by percentage of human CD298+ melanoma cells within the organs (Fig. 3F-G; Supp. Fig. 3F-G).

**Figure 3.**
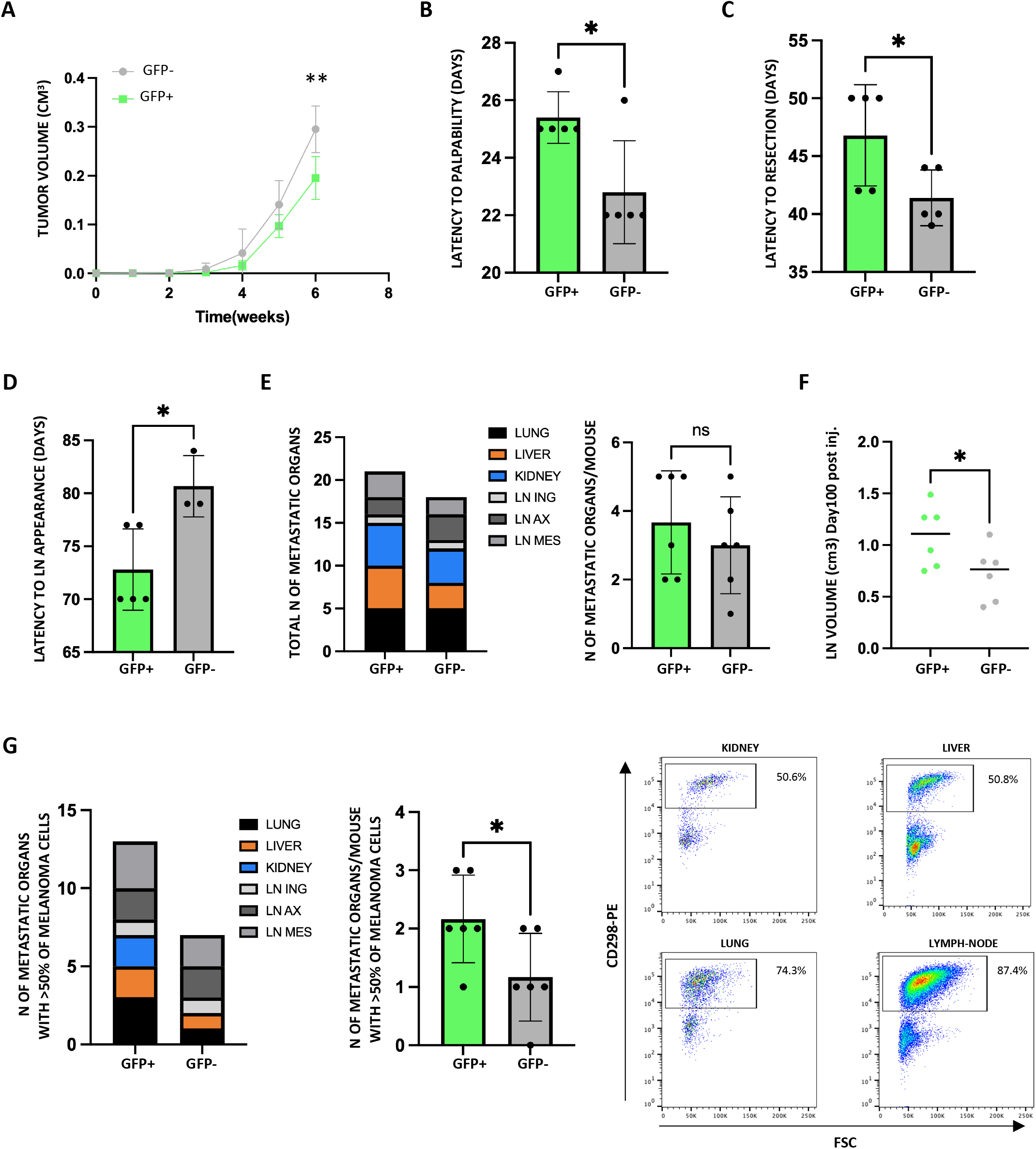
*In vivo* assessment of tumorigenic and invasive properties of GFP+ and GFP- MM13 populations in immunocompromised mice. **A)** tumor growth (at 6 weeks); **B)** latency to tumor palpability; **C)** latency to tumor resection (volume ∼0.3cm^3^); **D)** latency to lymph-node (LN) appearance; **E)** total number of organs with detectable metastases and average number of organs with metastases *per* mouse; **F)** LN volume at day100 after injection; **G)** total number of organs with at least 50% of melanoma cells and average number of organs with at least 50% of melanoma cells *per* mouse. A representative flow cytometry analysis is reported to show the assessment of melanoma metastatic cells within each organ based on the expression of human CD298. Data are mean ± SD (n=5). Student t test was applied to assess the significance (ns, not significant; *pvalue< 0.05; **pvalue<0.001).

### Oxygen levels affect the migratory and invasive properties of slow cycling cells

Metabolic features of tumor microenvironment are crucial in modulating tumor plasticity. Among relevant factors, low oxygen conditions have been shown to promote quiescence (15,22,45–48). We therefore decided to study how hypoxia impacts on our slow cycling melanoma phenotype. To do so, we used freshly sorted GFP+ and GFP– MM13 and MM27 cells and we measured their migration/invasive properties and resistance to trametinib treatment in low oxygen condition (3%O2). Our results clearly showed that hypoxia affects the quiescent cancer state properties, driving migration (Fig. 4A; Suppl. Fig. 4A), invasiveness (Fig. 4B; Suppl. Fig. 4B) and drug resistance (Fig. 4C; Suppl. Fig. 4C) more effectively than in normal oxygen conditions.

**Figure 4.**
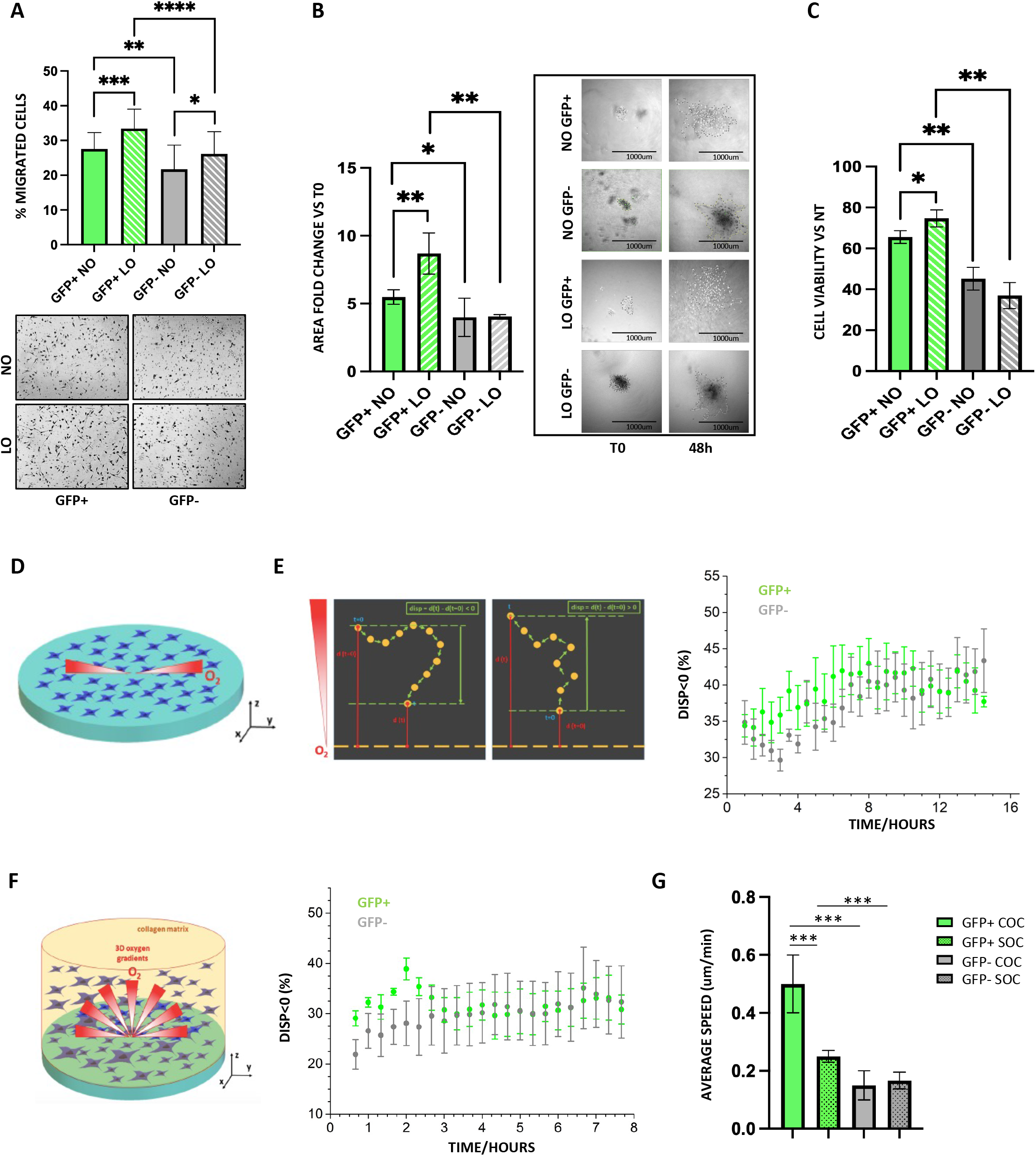
Impact of the hypoxic tumor microenvironment on MM13 slow cycling cells. **A)** GFP+ and GFP- cell migration was assessed by transwell migration assay after 24h in normal (20% O_2_) and low oxygen (3% O_2_) incubators. Representative EVOS microscopy images of migrated cells (transwell outer surface) are reported. Data are mean+-SD (n=3). p-values are based on unpaired Student’s t test (***p < 0.0001). The experiment was performed three times. **B)** Spheroid collagen invasion in GFP+ and GFP- cells in normal (20% O_2_) and low oxygen (3% O_2_) incubators was measured as area of invasion fold change vs T0, imaged every 24h by EVOS microscopy for T48h. Data shown are the mean ± SD of 10 different spheroids per group. Student t-test (****pvalue<0.00001). Representative images are shown. **C)** *In vitro* trametinib (10nM) treatment (72h) of GFP+ and GFP- MM13 cells in normal (20% O_2_) and low oxygen (3% O_2_) incubators. Cell viability was assessed by CyQuant and normalized to DMSO control. Mean ± SD (n=3). p values are based on unpaired Student’s t test (***p < 0.001). The experiment was performed three times. **D)** Schematic representation of the experiment conducted under spatially controlled oxygen conditions. **E)** Definition of displacement, *disp*, as function of distance of cell position at time *t*, *d(t)*, from zero oxygen level and % of disp<0 (% of cells migrating toward hypoxia) in GFP+ (green) and GFP- (gray) MM13 assessed during time-lapse of migration assay in chemical controlled culture system. Data are mean ± SD (n=3). Significance was assessed by one-way ANOVA (p value 0.02298 and confidence level >95%). Number of trajectories analyzed (GFP+ n=668; GFP- n=1158) **F)** Schematic representation of 3D invasion assays in controlled oxygen conditions and percentage of *disp*<0 in GFP+ (green) and GFP- (gray) MM13 assessed during time-lapse of invasion assay. Data are mean ± SD (n=3). Significance for the was assessed by one-way ANOVA (p value 0.02587 and confidence level >95%). Number of trajectories analyzed (GFP+ n=1601; GFP- n=976). **G)** Average speed of GFP+ and GFP- in 3D collagen invasion assay in controlled oxygen conditions (COC) and in standard oxygen conditions (SOC) were analyzed. Data are mean ± SD (n=3). Student t test was applied to assess the significance (ns, not significant; **p value<0.01; ***pvalue<0.001). Number of trajectories analyzed (GFP+ n=4693; GFP- n=1551).

To strengthen these data for a more relevant clinical application, we employed an *in vitro* model able to better reproduce the complexity of tumor niche oxygen conditions (49). The culture dishes are equipped with a technology that can modulate the concentration of oxygen from 0% to 20% in the culture plate (Fig. 4D). In this way, low oxygen concentration, i.e. from 9% to 0% (50), typically present in tissues or in the tumor microenvironment, can be replicated. Taking advantage of this system, we studied the ability of the melanoma MM13 and MM27 GFP+ and GFP– populations to migrate in 2D (Fig. 4E; Supp. Fig. 4D) and invade in 3D collagen matrix (Fig. 4F; Supp. Fig. 4E) under these controlled oxygen conditions. Following cells through time-lapse confocal analysis, we clearly showed different propensities of cells to move along oxygen gradients toward hypoxic conditions; in particular GFP+ populations adapt and better survive in hypoxic conditions, which in turn promote their migratory and invasive features in the first 8-16 hours of the experiments. Analyzing other cell motility parameters, we considered cell speed during the invasion assay and we observed that low oxygen conditions promoted an increased the average speed of GFP+ cell (Fig. 4G; Supp. Fig. 4F). All these data, collected in a microenvironment which better replicate the patient tumor niche, support the early and evident invasive phenotype of the slow cycling population we observed in *in vivo* experiments.

### scRNA-seq analysis of GFP+ and GFP- populations

To dissect the transcriptomic profiles of proliferating and slow cycling/quiescent melanoma cells during melanoma dissemination, we performed scRNA-seq analysis of MM13 GFP+ and GFP- primary tumors and matched lymph-node metastases. PCA analysis followed by a KNN-graph (k-nearest neighbor graph), Louvain clustering analysis and UMAP (uniform manifold approximation and projection) dimensionality reduction identified 14 transcriptional melanoma cell states (clusters) (Fig. 5A). The analysis of the cell cycle phases performed by Seurat and the assessment of KI67 expression levels showed that cells in the G1 phase of the cell cycle have the same transcriptional profile of low KI67 level cells and were assigned to specific clusters, mainly 0, 6, 7, 8, 12 and 13 clusters (Fig. 5A and Suppl. Fig. 5A). Similarly, proliferating high expressing KI67 cells were consistently assigned to cell cycle phases (S-G2M) in clusters 2, 4, 5, 9, 10, 11, 13 and partially in 1, 2, 3 (Fig. 5A and Suppl. Fig. 5A). We then grouped cells according to median KI67 expression (see Mat&Met), showing a strong consistency with the cell cycle phase annotation performed with the Seurat algorithm. In particular, KI67- and KI67+ populations overlapped with G1 and S-G2/M cells respectively, as clearly showed by UMAP projections and stack plots (Fig. 5A-B; Suppl. Fig. 5A-B). Gene set enrichment analysis (GSEA) was used to functionally annotate the 14 clusters (Fig. 5C). G1 phase clusters were enriched in specific pathways known to be involved in the invasive and metastatic process such as iron ion homeostasis (51–53), cell response to oxygen levels (15,22,47) and mitochondrial electron transport (7,16,54), while clusters of cells in S-G2M phases showed upregulation of cell cycle control and transition pathways (Fig. 5C). We then grouped melanoma cells according to their tissue of origin, namely primary tumor or metastases. Cell populations showed a clear transcriptional separation of the clusters on UMAPs (Fig. 5D, upper and middle panels) and on stack plots (Fig. 5D, lower panel), suggesting the key role of the tumor site and the niche in shaping the melanoma transcriptional landscape. GFP+ and GFP- populations showed only a partial segregation on UMAPs (Fig. 5E, left panel), with clusters 0, 1 and 13 mainly represented by GFP+ cells and clusters 7, 8 by GFP- cells, as shown in the stack plot (Fig. 5E right panel).

**Figure 5.**
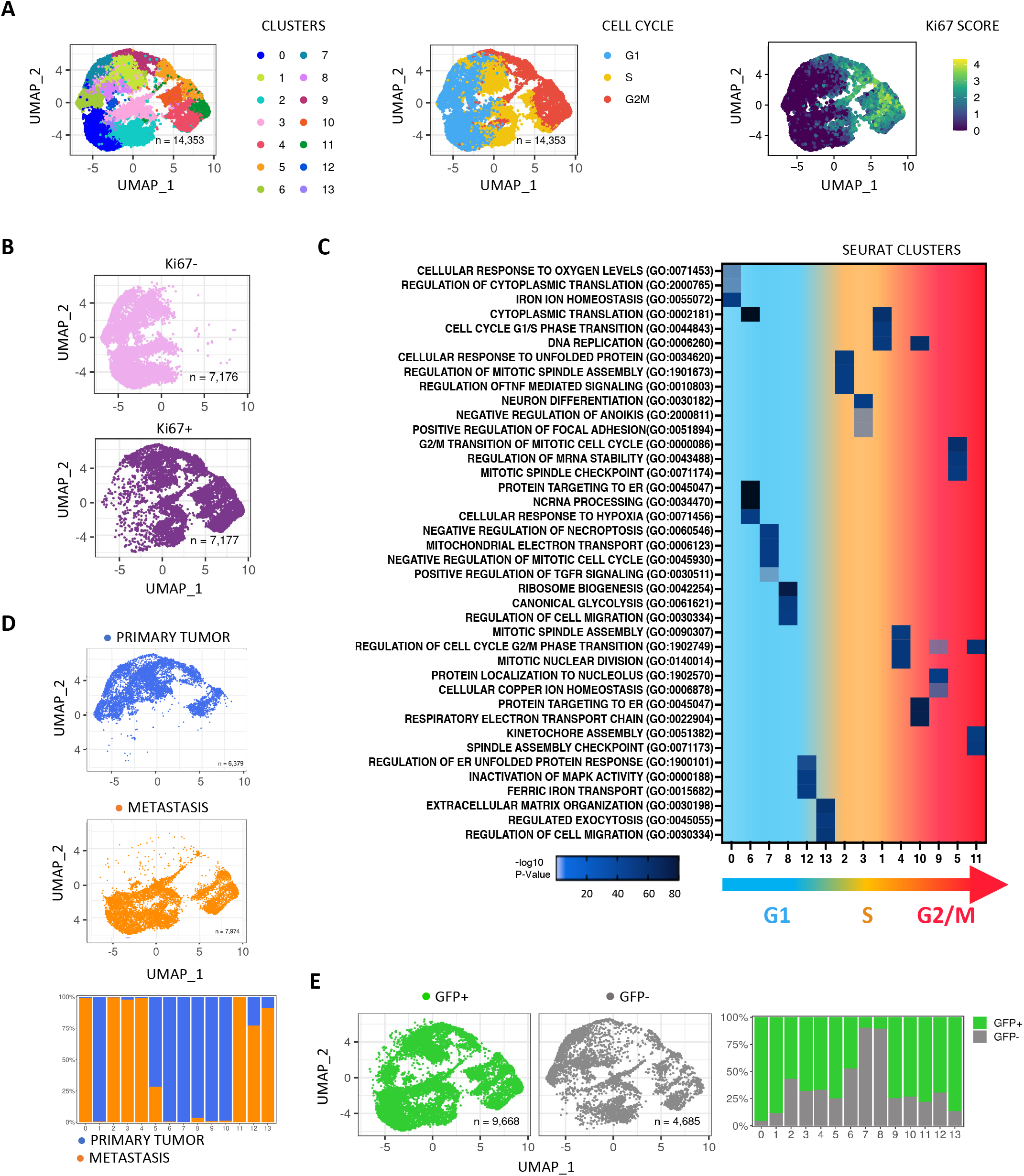
Single-cell transcriptional landscape of primary and metastatic MM13 melanoma. **A)** UMAP visualization of 14,353 MM13 melanoma cells analysed by scRNA-seq and integrated across 3 different melanoma primary tumors with 4 matched lymp-node metastatic lesions: cells colored by Seurat clusters (left), Seurat cell cycle annotation (middle), KI67 expression score (right). **B)** UMAP projection of KI67- and + MM13 melanoma cells, grouped on median of KI67 expression. **C)** Enrichment pathway analysis (GO Biological Processes) of the Seurat clusters grouped by cell cycle phase annotation. **D**) UMAP projection (upper panel) of primary and metastatic MM13 melanoma cells and stack plot (lower panel) of relative proportions of primary and metastatic cells across Seurat clusters. **E**) UMAP projection GFP+ and GFP- compartments of MM13 melanoma populations (left) and stack plot (right) of relative proportions of GFP+ and GFP- cells across Seurat clusters.

To better characterize our scRNA data, we annotated our populations using previously published melanoma signatures (12). The expression was assessed by UMAP projection (Supp. Fig. 5B, upper panel) and showed that GFP+/KI67-/G1 cell populations were significantly enriched in “MITF targets”, “invasion” and “MSC” melanoma signatures, while cells belonging to GFP-KI67+/G2/M clusters were enriched in the melanoma “mitotic” and “pigmentation” signatures (Supp. Fig. 5B, lower panel). To further explore the transcriptional heterogeneity of GFP-retaining cells, we arranged our dataset in 4 groups based on the relative expression of both GFP and KI67 genes: GFP+KI67- (QQ: fully quiescent cells), GFP+KI67+ (QP: cells retaining GFP and expressing KI67 at detectable levels), GFP-KI67- (PQ: proliferative cells that are losing KI67 expression) and GFP-KI67+ (PP: fully proliferative cells) phenotypes (Fig. 6A). Looking to the UMAP projections of the four subgroups, it was clear that the expression level of KI67 prevails on GFP expression in shaping the relationships between populations, revealing that PQ cells were more similar to QQ and QP to PP cells (Fig. 6A). We then defined a transcriptional signature for each of the four populations, identifying specific marker genes for each of them (Supp. Table1). QQ and PP populations were clearly separated on UMAP projections, while PQ and QP populations were partially overlapping with QQ and PP, respectively (Fig. 6A). Interestingly, the top ranked genes of the QQ signature showed intermediate expression levels in the PQ population, while they were downregulated in the QP and PP populations (Supp. Fig. 6A). Gene set enrichment analysis corroborated the distinct nature of the four populations, showing close similarities between QQ and PQ, and PP and QP, with the first two populations being mainly enriched in pathways involved in metabolism, angiogenesis, cell death control, oxygen stress and extracellular matrix organization, and the second ones in pathways related to cell cycle transition and mRNA processing (Fig. 6B).

**Figure 6.**
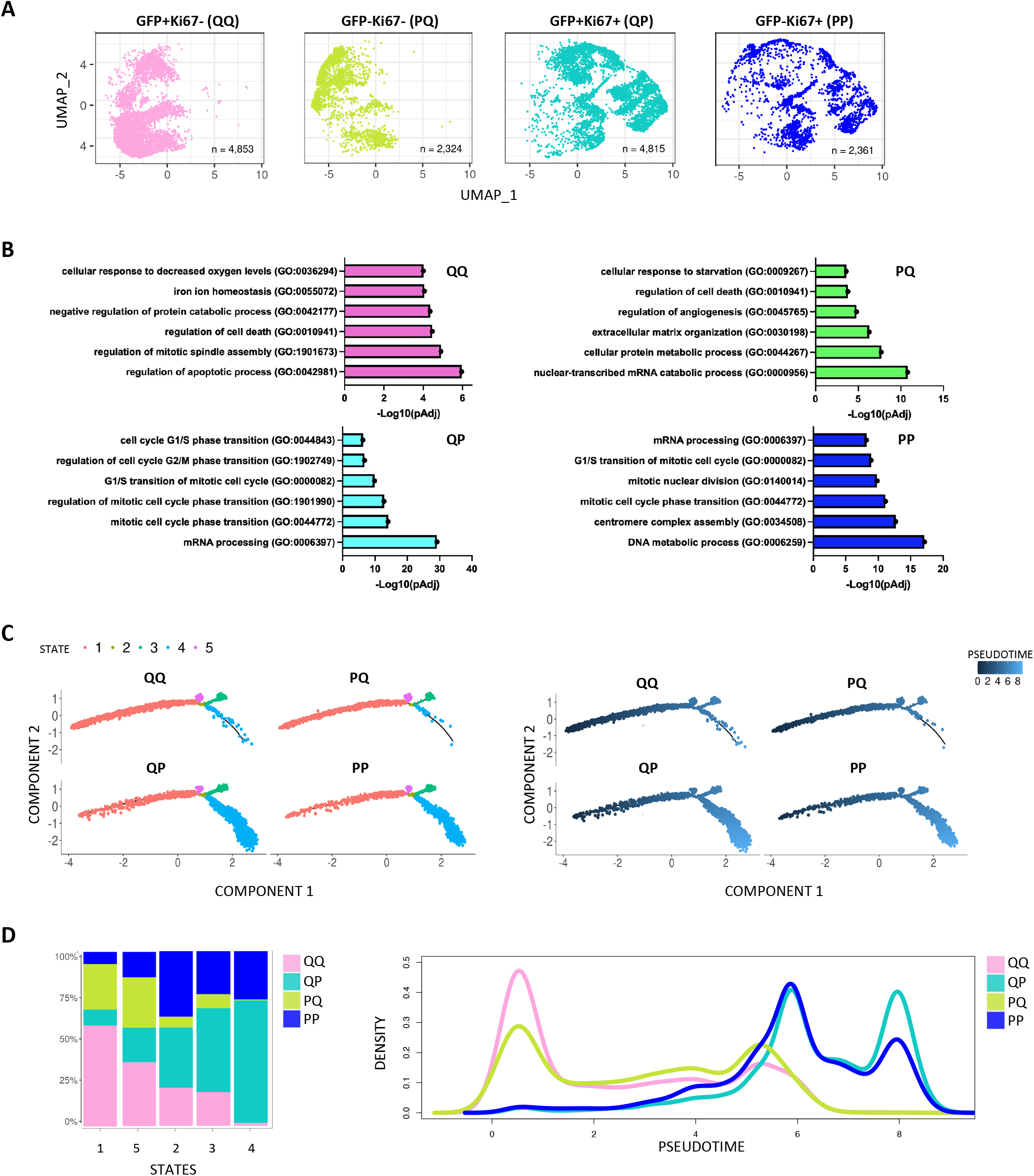
Single cell transcriptomic characterization of 4 cell populations grouped based on GFP fluorescence and KI67 expression. **A)** UMAP visualization of the four populations grouped on quiescent/proliferative state: GFP+KI67- (QQ), GFP-KI67- (PQ), GFP-KI6+ (QP), GFP-KI67+ (PP). **B**) Enrichment pathway analysis (GO) of the populations defined in A. **C**) Trajectory analysis of QQ, QP, PQ and PP populations based on the expression of the top 20 variable genes. The trajectories are colored by state (left panel) and pseudotime (right panel). State 2 was selected as root state. **D**) Stack plots (left) showing relative proportions of the 4 populations across 5 pseudotime states; density plots (right) showing distribution of the 4 populations along pseudotime.

To verify the quiescent phenotype of QQ cells, we assessed the expression of a set of “growth inhibitory genes”, obtained from the Molecular Signature Database (MSigDB) (see Mat&Met) (MODULE_488 GSEA), confirming that their expression was enriched in QQ/PQ compared to PP/QP populations (Suppl. Fig. 6B, left panel). Even more strikingly, we showed that the expression of these genes was mainly associated with the metastatic compartment, suggesting a role of the tumor niche in promoting cell quiescence (Suppl. Fig. 6B, right panel).

To get insights into the transcriptional dynamics that occur during the conversion from quiescent to proliferative states (or vice-versa), we performed a trajectory analysis with the monocle 2 R package. The analysis identified a tree structure profile with 5 different transcriptional states, being QQ cells mainly enriched at the left extremity of the trajectory in Fig. 6C, left panel, corresponding to the state 1. The proportion of QQ cells then progressively decreases moving toward the opposite edge of the trajectory (state 4) where, conversely, PP cells pile up. As expected, QP and PQ cells occupied intermediate positions in the trajectory path, clearly mimicking the enrichment trend of QQ and PP populations, being the PQ most strongly enriched in early states and QP in the late ones along the pseudotime (Fig. 6C, right panel), confirming previous observations which suggested this relative functional hierarchy between groups. Moreover, QQ/PQ and PP/QP cells were distributed with specific enrichments at different definite points along the pseudotime in the density plot, suggesting that melanoma cells switch between the two main functional endpoint states (QQ and PP) with rare stable intermediates (Fig. 6D).

We then performed a bulk RNA-seq analysis of KI67-fixed sorted MM13 cells (KI67+ and KI67- cells) and we identified 424 upregulated genes and 1,079 downregulated genes in KI67- compared to KI67+ cells (|log2FC| > 1, pvalue < 0.01) (Supp. Fig. 6C). Comparing the KI67-fixed sorted RNA-seq dataset with the QQ population we found seventy-seven upregulated genes (18%) in common (Supp. Fig. 6D), providing further confirmation for the quiescent cells signature.

To validate our QQ signature in a clinical setting, we took advantage of an RNA-seq dataset of melanoma patients pre- and post-RAF/MEKi treatment (GEO: GSE77940) and two scRNA-seq datasets of patients before and after immune checkpoint inhibitors treatment (GEO: GSE115978). Notably, GSEA analysis showed an enrichment of our QQ signature in relapsed patients treated with MAPK and immune checkpoint inhibitors, confirming the association of our QQ signature with a more aggressive and therapy-resistant phenotype (Supp. Fig. 6E). These data suggest that QQ signature genes might contain valuable targets for the development of treatments against therapy-resistant melanoma cell populations.

### Tumor niche pressure controls transcriptional reprogramming of quiescent cells

To further investigate the transcriptional heterogeneity of the QQ population, we re-clustered the cells using h-KNN and Louvain algorithms. The analysis identified 16 clusters, in which primary and metastatic QQ cells were clearly segregated on UMAP and were assigned to different clusters (Fig. 7A and Supp. Fig. 7B). This finding indicates that QQ cells undergo a transcriptional reprogramming after homing within the metastatic niche. The trajectory analysis confirmed the different transcriptional profiles of QQ cells in primary tumors and corresponding metastases. Indeed, using the top 20 variable genes, the analysis identified a trajectory profile with 3 different states, with state 1 as a root branch. Pseudotemporal gene expression analysis demonstrated that primary QQ cells were enriched at early stages of the trajectory and lost over pseudotime, while metastatic QQ were maintained or even enriched at later points of pseudotime (Fig. 7C-D). Moreover, we confirmed the enrichment of QQ cells in lymph node metastatic samples compared to each matched primary tumor (Fig. 7E). This observation is supported by FACS analysis of the GFP+ primary and metastatic population obtained after doxycycline chasing that showed an enrichment of these cells in lymph node metastases compared to primary tumor site (Fig. 7F). These data strongly suggest that KI67 negative GFP-retaining slow cycling cells are selective maintained and enriched under metastatic niche pressure.

**Figure 7.**
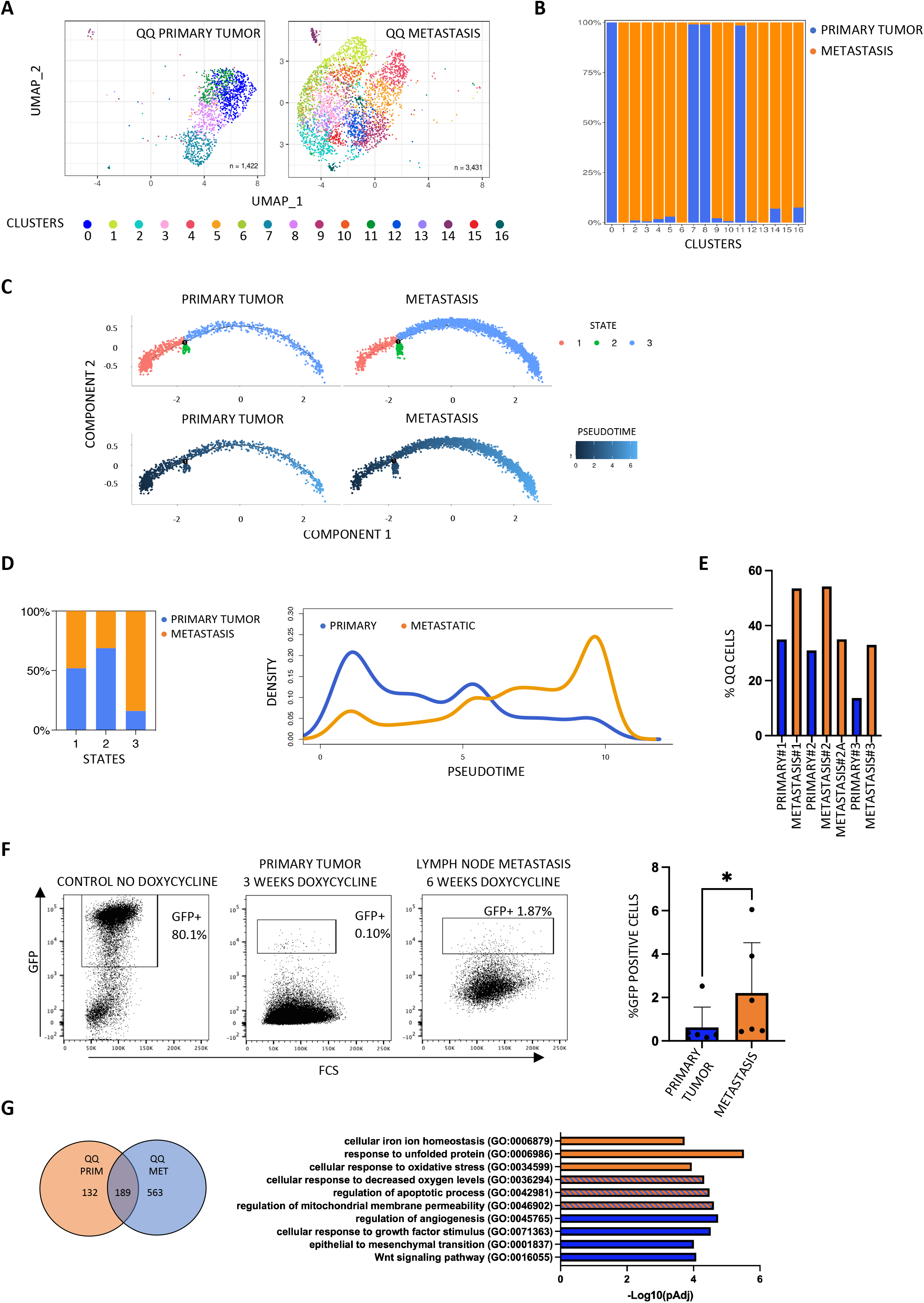
Analysis of transcriptional heterogeneity of MM13 melanoma QQ cells. **A)** UMAP projection of re-clustering analysis of QQ cells derived from primary tumor and metastatic samples. **B**) Stack plot showing relative proportions of primary and metastatic QQ cells across Seurat clusters. **C**) Trajectory analysis of primary and metastatic QQ cell populations based on the expression of the top 20 variable genes. The trajectories are colored by state (upper panel) and pseudotime (lower panel). State 1 was selected as root state. **D**) Stack plots (left) showing relative proportions of primary and metastatic QQ cells across pseudotime states; density plots (right) showing distribution of primary and metastatic QQ cells across along pseudotime. **E**) Bar plot showing percentage of QQ cells in each primary and matched metastatic lymph nodes. **F**) FACS analysis (representative dot plot) showing the percentage of GFP+ cells in primary tumors and matched metastatic lymph nodes. Student t test was applied to assess the significance (*pvalue< 0.05). **G**) Venn diagram (left) showing overlap of marker genes of QQ primary and metastatic populations; GSEA functional annotation (right) of 189 common marker genes.

To gain insight into the transcriptional features of primary and metastatic QQ cells, we identified 321 and 752 marker genes for the first and second group of QQ cells respectively, with 189 markers in common at FDR <0.05 (Supp. Fig. 6G). GSEA analysis showed that quiescence in primary tumor is linked to cellular iron homeostasis, stress response and apoptosis, while quiescence in metastases is associated with epithelial to mesenchymal transition, regulation of angiogenesis, Wnt pathway and response to growth factors (Fig. 7G). Both QQ primary and metastatic cells rely on the upregulation of pathways involved in cellular response to oxidative stress, regulation of apoptotic process and mitochondrial membrane permeability (Fig. 7G).

Finally, we compared the newly identified QQ signature with a recently published mesenchymal-like melanoma cell state signature which characterizes melanoma metastatic initiating cells (11). As shown in Supp. Fig. 6F some of the top ranked genes of the metastatic QQ signature (PRRX1, LUM, CTSK, SERPINF1 and SPARC) are members of the mesenchymal melanoma signature. Furthermore, the expression of PRRX1 regulon (PRRX1 target genes) was significantly higher in metastatic QQ cells compared to other populations (Supp. Fig. 6F). These data further suggest that metastatic QQ cells might be a fraction of the metastatic melanoma initiating cells.

## DISCUSSION

Cellular quiescence is a state of reversible growth arrest in which cells have exited the cell cycle, but they can re-enter it upon appropriate stimulation being still transcriptionally and metabolically active (55). Quiescent cells play a key role in the physiology of tissue homeostasis but they represent a major hurdle in cancer treatment. Indeed, quiescent cells are resistant to anti-cancer therapies and they can stay in a dormant state for long periods of time, reawakening to give rise to cancer recurrence and metastasis (56). In melanoma, quiescent cells can represent the population to target to prevent relapse (57). For this reason, an in-depth investigation into the cellular populations composing the melanoma at various progression stages, both in the primary site and in the metastatic lesions, can help resuming a clearer picture of such a highly heterogeneous and plastic tumor. To this end, we made use of the H2B-GFP reporter system, whose fluorescence halves at each cell division upon doxycycline treatment, to isolate and characterize *in vivo* slow cycling/quiescent cells in our human melanoma PDXs.

We herein describe a subpopulation of NRAS and BRAF mutant melanoma cells with slow cycling features that we have isolated from PDX primary tumors and matched metastases. The PDXs, well characterized in terms of mutation profile and clinical records, faithfully recapitulate the patient genetic and phenotypic landscape and represent an excellent model of spontaneous metastasis when injected intradermally. We clearly demonstrate that GFP+ slow cycling cells display an increased capability of migrating, invading matrix *in vitro* and tissues *in vivo*, and resisting to therapy. A similar approach was previously used by Puig et al to define a slow cycling, chemoresistant melanoma population with a distinctive gene expression profile, enriched in pathways related to drug detoxification, stemness, hypoxia, and crosstalk with the immune system (18). The authors showed that this population is common to melanoma, glioblastoma and colon carcinoma identifying the TET2 epigenetic enzyme as a crucial factor controlling the number and survival of slow cycling cells, and tumor recurrence in the end, without characterizing specific transcriptional profiles in primary tumors and metastases. To complete our GFP+ slow cycling cell characterization considering the metabolic features, we show that they preferentially use mitochondrial oxidative phosphorylation (OXPHOS). Interestingly, a quiescent melanoma population recently isolated by dual labelling of endogenous p27 and KI67 of melanoma cell lines (20), shows high c-Myc expression, which drives cell survival and OXPHOS metabolism. This is in contrast with our data where the enhanced oxidative phosphorylation (Fig. 2E; Suppl Fig 2E) does not correlate with an increased expression of c-Myc (data not shown), suggesting that different pathways might be activated in cell lines and PDXs.

Taking together, our data strongly support a critical role of slow cycling cells in the invasive process that underlies metastasis dissemination. As tumor microenvironment features, such as hypoxia, acidity, nutrient deprivation, have been described as relevant factors in promoting quiescence, tumor progression and resistance (58), to better dissect how the TME impacts on melanoma cell subpopulations, we used a novel *in vitro* model that was manageable, that reproduced the characteristics of the *in vivo* tissue and in which oxygen levels could be modified at will. Previous works made use of the "organs on a chip" technology based on microfluidics, which is claimed to reproduce the tumor tissue, but still lacks full representation of the microenvironment (59,60). We therefore used a technology that was developed to modulate oxygen levels in normal culture conditions (49). This culture system allowed us to study the specific functional properties of slow cycling cells, such as migration, invasion and cell velocity in a setting where oxygen levels can be fine-tuned. In the future, it will provide us with a strong tool to investigate the distinctive features of isolated quiescent populations by fine tuning the parameters of the tumor niche. Slow cycling cells can then be retrieved and further analysed for their survival and therapy resistance.

In further characterizing our slow cycling population, we observed the co-existence of KI67- and KI67+ cells within the GFP population at the protein level. Indeed GFP+ cells showed different levels of the proliferation marker that can account for the presence of a long- and short-term quiescent populations. To refine the picture, single-cell RNA-sequencing (scRNA-seq) analyses, applying a digital filter for KI67 expression, unveiled four different populations, the quiescent (QQ: GFP+KI67-) and the partially quiescent (PQ: GFP-KI67-), the partially proliferative (QP: GFP+KI67+) and the fully proliferative (PP: GFP-KI67+) populations. The PQ state shares a transcriptional profile closer to the QQ state, even if they have peculiar functional states as the pseudotime analysis highlights. GSEA analysis confirms that they both have a specific role in the invasive process since they upregulate pathways, such as cellular response to starvation/cell metabolic process and regulation of cell death, involved in the acquired capacity of surviving in hostile conditions, and extracellular matrix organization/regulation of angiogenesis, involved in the invasion and in the creation of a pro-tumorigenic metastatic niche. The QP state instead is closer to the PP state, with a signature fully related to cell cycle transition control. These similarities suggest that KI67 expression defines the phenotype of the populations, indicating its use in drawing specific temporal trajectories of what might be the biological paths that the populations follow during tumor progression. Based on the transcriptional profile of melanoma populations and KI67 expression, we can speculate that melanoma is composed of 2 groups of cells, represented by the two opposite edges of the trajectory, QQ/PQ and PP/QP. Within these two groups, cells display interchangeable profiles, where KI67 expression subdivides the cell states. Indeed, if we analyze the cluster composition of quiescent cells (QQ and PQ), beside a major overlap, we can also observe some differences, such as the clusters 0 and 7 that are present only in QQ and in PQ respectively. This suggests that quiescent cells have a dynamic phenotype, based on KI67 expression but also on a more complex transcriptional profile. Our data support the idea that quiescent QQ cells, that never exit the G0 state, show a specific adjustment to iron ion homeostasis and to oxygen level, whose important impact and driving force was confirmed in our low oxygen experiments. While hypoxia conditions are known to play a role in pushing cells towards quiescence (15,22,45–47) the iron ion homeostasis has been linked more to cancer stemness maintenance (61–63), and its specific involvement in quiescence will deserve further studies.

Strikingly, if we re-cluster the QQ cells according to tumor site, either primary or metastatic, we can find another level of heterogeneity. The QQ transcriptomic profile of the two different stages are clearly unique with iron ion homeostasis as master regulator of the primary tumor and response to oxygen level/angiogenesis of the metastasis. The metastatic site drives a well-defined quiescent phenotype with a stronger correlation with the GFP+KI67- segregation. Moreover, the clinical patterns resemble the transcriptomic and phenotype dynamics since the QQ signature is significantly enriched in relapsed or resistant treated melanoma patients. This finding strongly suggests that contributions to the development of specific cancer phenotypes by the expression of distinct genes are dependent on the specific features of the niche associated with tumor progression, adding another layer of complexity to the molecular mechanisms underlying the metastatic process.

To conclude, we need to better understand the biological events that drive and maintain the quiescent phenotype at the metastatic site. It is also crucial to target the quiescent cells both in early stages of the primary tumor growth, to avoid their dissemination before surgical intervention to prevent metastases formation, and in late stages to block the metastatic process and halt relapse.

## MATERIALS AND METHODS

### Patient-Derived Xenografts (PDXs) generation, characterization and *in vitro* culture

Tissue biopsies of metastatic melanomas were collected from patients whose informed consent was obtained in writing according to the policies of the Ethics Committee of the European Institute of Oncology and regulations of the Italian Ministry of Health. The studies were conducted in full compliance with the Declaration of Helsinki.

Metastatic melanoma PDXs were generated in NOD.Cg-Prkdcscid Il2rgtm1Wjl/SzJ (NSG) mice as previously reported (64). For the herein described experiments, MM13 and MM27 PDXs were used for *ex vivo* and *in vivo* studies. For the *in vitro* functional experiments PDXs were maintained in culture in Iscove’s modified Dulbecco’s medium (IMDM, Sigma Aldrich-Merck, Cat#I3390) supplemented with 200 mmol/L L-glutamine (Euroclone, Cat#ECB3000D) and 10% FBS (HyClone products Cytiva, Cat#SH30071.03IH).

### *In vivo* generation of H2B-GFP model

*In vivo* studies were performed after approval from our fully authorized animal facility and notification to the Ministry of Health (as required by the Italian Law; No 564/2019-PR), and in accordance with EU directive 2010/63. Mice of both genders were used at the age of 7–8 weeks.

MM13 and MM27 cells were transduced with the H2B-GFP reporter system (37), which allows the tracking of cell division based on the inducible (Tet-off) expression system. In the absence of doxycycline, H2B-GFP is stably expressed; once doxycycline is added, the GFP signal is reduced at each cell cycle division, allowing the distinction of slow cycling (GFP positive) and proliferating (GFP negative) cell populations. 2×10^5^ PDX melanoma cells, stably infected with the H2B-GFP vector, were injected intradermally in the back of immunocompromised mice. At palpability (≅2 weeks), mice were treated with doxycycline via modified food pellets at a dose of 625 mg/kg for ≅3 weeks before sacrifice or, for the metastatic model, before tumor was resected at the volume of ∼0.3 cm^3^. Tumor volumes were monitored and annotated at least twice. Tumor volume was calculated using the modified ellipsoid formula: 1/2 (length × width^2^). To monitor metastasis onset, mice were kept under doxycycline treatment for additional 3 weeks before sacrifice. Autopsies were performed to check for the presence of metastases.

### Fluorescence-activated cell sorting

Sorting was performed using fluorescence-associated cell sorters (Aria and Melody, BD Bioscience) to isolate GFP+ and GFP-populations from primary tumors after DAPI staining for live cells and from mice metastatic organs after labeling with human CD298-PE (BD, Cat#749741, clone P-3E10) and DAPI for human melanoma and live cells. Analysis was performed using the FlowJo software (BD; version 10.5.2 or higher).

### *In vivo* bromodeoxyuridine (BrdU) incorporation

BrdU incorporates newly synthesized DNA, enabling the identification of actively proliferating cells. 2×10^5^ MM13 cells were injected intradermally in NSG mice (n=8). For the *in vivo* long pulse chase BrdU assay, mice were immediately injected intraperitoneally with 1.5 mg of BrdU (Sigma-Aldrich, Cat#B5002) in sterile PBS or PBS as control (4 mice per group), then they received 1mg/ml of BrdU in drinking water added with 5% glucose for 10 days or normal water in control mice. Control mice were then treated with doxycycline, while the remaining 2 BrdU-treated and 2 non-treated animals used as controls. After ≅3 weeks of chasing, mice were sacrificed and primary tumors collected, dissociated at single cell suspension, fixed in 70% of ethanol, denatured with 2 M HCl for 25′ and the reaction stopped adding Sodium Borate pH 8.5 for 2’. Cells were then labeled with primary anti-BrdU antibody (BD Bioscience, Cat#347580) and secondary antibody. BrdU incorporation was assessed by FACS (Celesta, BD Bioscience) flow cytometry. Data were analyzed using the FlowJo software (BD; version 10.5.2 or higher).

### *In vitro* bromodeoxyuridine (BrdU) incorporation

For *in vitro* short pulse chase, 3.000 freshly sorted MM13 and MM27 GFP+ and GFP- cells were seeded in 96 well-plates in triplicates and 10uM of BrdU (Sigma-Aldrich, Cat#B5002) was added to complete culture medium. After 24h cells were fixed in 70% of ethanol, denatured with 2 M HCl for 25′ and the reaction stopped adding Sodium Borate pH 8.5 for 2’. Cells were then stained with primary anti-BrdU antibody (BD Bioscience, Cat#347580) and secondary antibody and analyzed by confocal microscopy (10X objective). All wells were acquired and GFP+ and BrdU+ cells were counted using ImageJ Software (National Institutes of Health, Bethesda, MD).

### Immunofluorescence staining for FACS

For KI67 studies, freshly sorted GFP+ and GFP– PDX melanoma cells were fixed and permeabilized with Cytofix/Cytoperm and Cytoperm Permeabilization Plus buffers (BD). Cells were then stained with human KI67-APC antibody (Biolegend, Cat#151206, clone 11F6).

For metastases studies, organs were dissociated mechanically and enzymatically with collagenase type III (Worthington Biochem, Cat#LS004182) at 37°C to single cell suspension and GFP+ and GFP- cells were collected from each organ (lung, liver, kidney and lymph-nodes) and stained for human CD298 (PE conjugated) to mark human metastatic cells within mouse organs. Stained cells were analyzed by FACS (Celesta, BD Bioscience) and data were analyzed using the FlowJo software (BD; version 10.5.2 or higher).

### Transwell migration assay

The migration assay was performed using 8.0 μm pore size (Corning, cat# 353093), fibronectin-coated (5 μg/cm^2^ coating the outer part of the membrane, Roche, cat#11080938001) inserts in 24-well plates (Corning-Falcon cat# 353097). Triplicates of 1,5 x 10^4^ freshly sorted MM13 and MM27 GFP+ and GFP- cells were plated in the upper chamber in IMDM serum-free medium. Complete medium was added to the lower chamber. After 24 hours in a 37°C incubator (20%O_2_ for normal oxygen condition and 3%O_2_ for low oxygen condition), non-migrated cells were recovered from the upper surface by PBS washing and membrane tip scraping, while cells migrated to the lower surface of the inserts were stained with 0.5% Crystal Violet solution (50% Crystal Violet 1% Sigma V-5265, 35% ethanol in water). Four images of each insert were acquired at EVOS microscope and analyzed with the ImageJ Software to estimate the occupied area and calculate migration rate.

### Spheroid collagen invasion assay

1000 freshly sorted GFP+ and GFP- MM13 and MM27 cells were grown as hanging drops in IMDM complete medium (Sigma Aldrich-Merck, Cat#I3390) for 72 h at 37°C in normal (20%O_2_) and low (3%O_2_) oxygen conditions. Single spheroids were harvested, resuspended in 1.5 mg/mL Collagen Type I Rat Tail (Corning, Cat#354249) and plated on a 96 well-plate. After 1 h at 37°C at either 20%O_2_ or 3%O_2_, the solidified collagen gel was covered with IMDM supplemented with 2%FBS. The ability of cells to invade the area was monitored and images of 10 spheroids *per* group were collected at 24h and 48h using a 4× objective lens. The invasion area was analyzed with the ImageJ Software and normalized against the area at time 0.

### *In vitro* drug treatment viability assay

*In vitro* drug sensitivity was assessed by CyQuant (Invitrogen, Cat#C35012), plating 3000 freshly sorted GFP+ and GFP- MM13 and MM27 cells in 96-well plates in triplicates. Cells were treated by a single exposure to either vehicle (DMSO), 10nM trametinib (GSK1120212, catalog number A-1258) for 72 hours at 37°C at 20%O_2_ or 3%O_2_. After adding CyQuant reagent, fluorescence signal was acquired with PHERAstar FSX Microplate Reader (EuroClone, BMG Labtech) and the relative viability (%) was calculated upon normalization to DMSO controls.

### Assessment of mitochondrial function

Seahorse experiments were performed using XF Cell Mito Stress Kit (Seahorse Bioscience). Freshly sorted MM13 and MM27 GFP+ and GFP- cells were seeded at equal density of 40,000 cells into XF96 tissue microculture plates pre-coated with 50 μg/ml poly-D-lysine. Cells were incubated for 24 h in standard growth medium IMDM in a humidified incubator at 37°C with 5% CO2. After 24 hours, the standard medium was replaced with XF DMEM Medium pH 7.4 supplemented with 25 mM glucose, 2 mM L-glutamine, and 1 mM sodium pyruvate. The cells were then incubated for 1 hour at 37°C without CO2 before running the assay. The drug injection ports of the XF96 Assay Cartridge were loaded with assay reagents to a final concentration of 1 μM oligomycin (ATP synthase inhibitor), 2 μM FCCP (uncoupler), 0.5 μM rotenone, and 0.5 μM antimycin A (complex I/II inhibitors). OCR was measured under basal conditions and prior to and following additions of the metabolic modulators. Data were normalized to urea lysate protein levels and analyzed using Wave Desktop Software (Seahorse Bioscience), following the manufacturer’s instructions. For mitochondrial studies, GFP+ and GFP- cells were incubated with 50 nM MitoTracker Deep Red FM (mitochondrial mass) and 25 nM MitoTracker Orange CMTMRos (mitochondrial activity) for 30 min at 37°C before cytometry analysis. Reagents were obtained from Life Technologies. All FACS data were acquired using a FACS (Celesta, BD Bioscience) and analyzed with FlowJo software.

### *In vivo* functional experiments

For *in vivo* functional studies, 10^4^ MM27 and MM13 GFP+ and GFP– cells were intradermally injected into the flank of NSG mice (5 mice *per* group) with Matrigel Matrix HC (Corning, Cat#354248) and L15 medium (Sigma-Aldrich, Cat#L5520). Tumor growth was monitored every 3 days using a digital caliper and resected when the tumor reached ∼0.3 cm^3^ in volume. The statistical difference in tumor volume among the two groups was assessed by Student’s t-test. Survival analysis was calculated with GraphPad Prism 9.0, and differences among groups were estimated by using Log rank test. All mice were sacrificed when lymph-nodes were detected, autopsies were performed and all the organs were collected and checked for metastases.

### Culture plates for controlled oxygen conditions

The culture plates for the experiments in controlled oxygen conditions have been detailed elsewhere (49) (Becconi M. et al., submitted). For invasion assay, cells suspended in the collagen mixture were deposited on the surface of the well for the oxygen control. The collagen matrix and cell dispersion were prepared as detailed previously in the *spheroid collagen invasion assay* paragraph with cell quantities reported in the next paragraph.

### Timelapses of PDX melanoma cells under controlled oxygen condition

Migration and invasion assays for sorted GFP+ and GFP- cells in the presence of controlled oxygen conditions were obtained with Nikon Eclipse Ti2 spinning disc confocal89 and Leica Thunder microscopes. Migration and invasion experiments were performed for 16 and 8 hours under controlled atmosphere and temperature (5% CO_2_ and 37°C). For migrations, after sorting, GFP+ and GFP- fractions were counted and 3000 cells seeded in 300 µl of growing medium in the modified well culture plates. Samples for invasion assays were prepared by seeding 6500 cells in 37 µl of 2.5 mg/ml collagen in growing medium. 300 µl of growing medium are then added in each well. Green (excitation 480/30 nm; emission 535/45 nm) fluorescence and differential interference contrast (DIC) images were collected every 20 minutes for 8 hours for invasion assays and every 15 minutes for 16 hours for migration assays; data from 3 independent samples were measured under controlled oxygen conditions for both GFP+ and GFP- cells.

### Quantitative analysis of trajectories of PDX melanoma cells under controlled oxygen conditions

Quantitative analysis of cell trajectories extracted from migration and invasion assays in the presence of controlled oxygen conditions were performed with scripts and computer routines written for Fiji (65) and MATLAB version 2020a software. Segmentation was performed for both migration and invasion assays respectively on DIC and green fluorescent images by using the pixel classification method in Ilastik (66); training of the machine learning algorithm was performed on a single green fluorescent image for each timelapse, after applying on the original images a gaussian smooth filters available on Ilastik. Watershed algorithm implemented in Fiji was applied to refine identification of neighboring cells. Refinement of binarized labeled images were obtained employing an in-house developed MATLAB routine. Bidimensional and tridimensional tracking for migration and invasion assays were performed with TrackMate plug-in available in Fiji. Trackmate identification of single cells was performed by setting estimated blob diameter and threshold for the LoG detector equal to 10 and 0 µm, respectively; trajectories reconstruction was obtained with linking max distance and gap-closing max distance equal to 10 and 0 µm, respectively.

Cell position with respect to pixel coordinate in the bidimensional images of timelapse data were converted to displacement, *disp*, values as described in Figure 4 and its caption. All the statistical analyses which followed cell trajectory reconstruction were performed with in-house developed MATLAB routines.

### Sample preparation for scRNA-seq

3 primary MM13 PDXs and 4 matched lymph nodes (1 for primary tumor#1 and #2 and 2 for primary tumor#3) freshly sorted for GFP expression were counted, resuspended in an appropriate volume of 0.04 % BSA/PBS (10x Genomics guidelines) and submitted for sequencing. On average, 2,000-3,000 cells per sample were used for sequencing. Cells were processed and libraries were prepared with Chromium Single Cell 3’ Reagent kit v2 (10xGenomics), Chromium Single Cell 3’ Reagent kit v3 (10x Genomics), or Chromium Next 48 GEM Single Cell 3’ Reagent kit v3.1 (10x Genomics). Paired-end sequencing at high coverage (50,000 reads per cell) was performed on Illumina NovaSeq 6000 platform.

### Single-cell RNA-seq data pre-processing and analysis

Sequencing results were converted to FASTQ format using Illumina bcl2fastq software. The Cell Ranger Single-Cell Software Suite v6.0.1 (https://support.10xgenomics.com/single-cellgeneexpression/software/pipelines/latest/what-is-cell-ranger) was used to perform barcode processing and single-cell 3′ gene counting. The cDNA insert was aligned to the GRCh38 human reference genome. Only confidently mapped, non-PCR duplicates with valid barcodes and UMIs were used to generate the gene-barcode matrix containing 15 627 cells. Further analysis, including quality filtering, identification of highly variable genes, dimensionality reduction, standard unsupervised clustering algorithms, and discovery of DEGs, was performed using the Seurat R package 6.1 (67). Additional conservative cut-offs were further applied based on the number of genes detected per cell (> 100), and the percentage of mitochondrial unique molecular identifier (UMI) counts (< 30). To limit background noise and increase robustness of differential analysis only cells with at least 4000 features were selected for further analysis (14,853). PCA analysis was performed to reduce data dimension and the cells were clustered in a two-step process. First, the cells were embedded in a k-nearest neighbor (KNN) graph and then, the Resolution-Optimized Louvain algorithm was applied. The dimensionality reduction and clustering visualization was performed using UMAP (Uniform Manifold Approximation and Projection) approach. The marker genes that define the clusters were then found via Seurat’s FindAllMarkers function using the ‘wilcox’ method (Wilcoxon rank sum test).

The marker genes that define the QQ in primary tumor and the QQ in metastases were found via Seurat’s FindAllMarkers function versus all the other populations. Venn Diagram was used to highlight common genes. Cell cycle score assignment was performed using CellCycleScoring function from Seurat package. Gene set variation analysis (GSVA R package with RNA-seq mode) was used to identify the enrichment of each gene set across clusters and cell populations (68). False discovery rate adjusted p-values (qFDR) were computed using the method of Benjamini & Hochberg. “GO biological processes”, “KEGG Canonical pathways” and the “Hallmark gene sets” were obtained from the MsigDB website (http://software.broadinstitute.org/gsea/msigdb) as well the ‘growth inhibitory genes’ gene set (https://www.gsea-msigdb.org/gsea/msigdb/cards/module_488) and the cell cycle-related gene set. Gene sets with FDR adjusted p-value qFDR < 0.05 were considered significant. Cerebro App package (69) was used for interactive visualization of single cell RNA-seq data. The dataset containing FACS sorted GFP positive and negative populations was grouped in 4 classes (GFP+/KI67- (QQ), GFP+/KI67+(QP), GFP-/KI67-(PQ) and GFP-/KI67+ (PP) based on median KI67 expression score. Re-clustering of QQ population was performed by PCA analysis and subsequent KNN-graph and Louvain alogorithm based clustering based on expression 2000 most variable features. Trajectory and pseudotemporal gene analysis were performed using Monocle 2.20.0 version. Genes, whose expression was changed along pseudotime were determined using *differentialGeneTest* function and 20 most variable genes for each dataset were used to construct the cell trajectories.

### RNA-seq bulk on KI67 positive and negative fixed melanoma PDX

MM13 PDXs cells were incubated with LIVE/DEAD Fixable Viability Dye, then fixed using PFA 1% supplemented with RNAse Inhibitor, permeabilized with saponin 0.1% supplemented with RNAse Inhibitor and stained for KI67-647 antibody. Samples were sorted at 4C for KI67 expression and sub-populations collected. RNA extraction was immediately performed using the RNeasy FFPE RNA extraction kit (Qiagen) according to manufacturer’s instructions.

RNA-seq libraries were prepared from 1μg of total RNA using the <u>TruSeq</u> <u>RNA Library</u> Preparation Kit v2 according to manufacturer’s instructions. The libraries were additionally purified using AMPure beads (Beckman Coulter), quality checked at Agilent 2100 Bioanalyzer and 50 bp paired-end sequenced on an HiSeq 2000 Sequencing System (Illumina). RNA-seq reads were aligned to genome (hg19, GRCh38) using TopHat2 2.0.9 (70) starting from 3 × 10^7^ mapped paired-end reads per sample. Read counts of each gene were quantified using HTseq (71) and differential analysis was performed using DESeq bioconductor packages (71). Genes were identified as differentially expressed (DEGs) when the following criteria were met: log_2_fold-change (FC) ≥ |1|, pvalue <0.05. DEGs were analyzed with Gene Ontology term enrichment using DAVID tool (version 6.8 Beta) (72), Ingenuity Pathway Analysis (IPA) and with GSEA (Gene Set Enrichment Analysis) software v2.2.0 (73). Gene sets enriched at <u>FDR</u> <=0.05 or 0.1 were considered statistically significant.

### Quantification and statistical analysis

Data are represented as mean ± SD of biological triplicates (if not, differently indicated in the text). Comparisons between two or more groups were assessed by using unpaired two-tailed Student’s *t-test*. For the statistical difference in tumor volume among different populations, unpaired *t test* was used. For all the statistical tests: ns, not significant; ^∗^, p< 0.05; ^∗∗^, p< 0.01; ^∗∗∗^, p< 0.001; ^∗∗∗∗^, p< 0.0001

### Data and code availability

Single-cell RNA-seq data will be deposited at European Nucleotide Archive (ENA) and publicly available as of the date of publication. Any additional information required to re-analyze the data reported in this paper is available from the lead contact upon request.

## CONFLICT OF INTEREST

The authors state no conflict of interest.

## ACKNOWLEDGMENTS

This work was supported by the Ricerca Finalizzata 2018 Grant *GR-2018-12367747* to FL, and the Italian Ministry of Health with Ricerca Corrente and 5×1000 funds.

The authors wish to thank Alberto Gobbi and Manuela Capillo (Cogentech) for the excellent support in animal work, the IEO Genomic Unit, the FACS and sorting, and Imaging facilities for the considerable technical support.

## AUTHORS CONTRIBUTION

Conceptualization: FL, PGP, LL; Data Curation: FL, MM, LuL, MMa, NQ; Investigation: FL, MM, LuL, MMa; Methodology: FL, VM, IF, TV, DB, MMa, NQ, and MM; Project Administration: FL, LL; Software: MM, LuL, MMa, NQ; Supervision: LL; Validation: FL, SR, LuL, LL; Funding Acquisition: FL, LL; Writing – original draft preparation: FL, LL; Writing – review and editing: FL, MM, LuL, MMa, SR, LL.

## CONTACT FOR REAGENT AND RESOURCE SHARING

Further information and requests for resources and reagents should be directed to and will be fulfilled by the Lead Contact, Luisa Lanfrancone

## Notes

### Competing Interest Statement

The authors have declared no competing interest.

