## Supplementary Figures, legends and Table for "*IN VIVO* ISOLATION OF A QUIESCENT MELANOMA POPULATION WITH INVASIVE PROPERTIES UNVEILS A TRANSCRIPTIONAL REPROGRAMMING DRIVEN BY THE TUMOR NICHE": Supplementary.pdf

**Supplemental Figure 1.** Isolation of slow cycling MM27 PDX cells *in vivo* by label-retaining H2B-GFP tet-off vector expression.

**A)** Scheme of the doxycycline-repressible histone H2B-GFP reporter system, stably incorporated into the DNA, constructed by combining Tet-Off regulatory elements with the Histone 2B-GFP (H2B-GFP) fusion construct, in a lentiviral HIV-1 based vector backbone. P<sub>EF-1α</sub>, elongation factor 1α promoter; tTA2 and Ptight, Tet-trans-activator and promoter; LTR/SIN, self-inactivating retroviral long terminal repeat sequences; RRE, HIV Rev response element; cPPT, central polypurine tract; WPRE, *Woodchuck* hepatitis virus post-transcriptional regulation element. **B)** GFP+ MM13 (upper panel) and MM27 (lower panel) were serially passaged (3 passages) *in vivo* in NSG mice to purify the GFP+ population. Representative flow cytometry analysis of GFP+ and GFP- separation over passages are shown. **C)** H2B-GFP transduced MM27 cells were chased *in vivo* with doxycycline. Representative dot-plot flow cytometry analysis (upper panel) and distribution histograms of GFP expressing cells (lower panel) show cell proliferation (gradual GFP lost) over time with around 0.5% of the cells retained the maximum label after 3 weeks of chasing. **D)** Percentage of GFP+ MM13 and MM27 cells isolated by cytofluorimetry after doxycycline chasing in 6 mice *per* PDX. **E)** GFP+ and GFP- MM27 cells were pulsed for 24h with BrdU *in vitro* and chased for 24h. Cells were analyzed by confocal microscopy (representative images) and counted for GFP and BrdU expression by ImageJ. Data are presented as mean ± SD (n=5). Student t test was applied to assess the significance (\*\*\*\*pvalue<0.00001). The experiment was performed twice. **F)** GFP+ and GFP- MM27 cells were stained with KI67 and analyzed by flow cytometry. Data are presented as mean ± SD (n=3). Student t test was used (\*\*pvalue<0.01). The experiment was performed 3 times. **G)** GFP+ MM27 cell proliferation was assessed by CyQuant over GFP- as mean ± SD (n = 3). p-values are based on unpaired Student's t test (\*\*\*\*p < 0.00001). The experiment was performed three times.

**Supplemental Figure 2.** *In vitro* functional characterization of MM27 slow cycling cells.

**A)** GFP+ and GFP- MM27 cell migration was assessed by transwell migration assay at 24h. Representative EVOS microscopy images of migrated cells (transwell outer surface) are reported. Data are mean±-SD (n=3). p-values are based on unpaired Student's t test (\*\*\*p < 0.0001). The experiment was performed three times. **B)** Spheroid collagen invasion in GFP+ and GFP- MM27 cells was measured as fold change vs T0 (imaged by EVOS microscopy every 24h for 48h). Data are shown as the mean ± SD of 10 different spheroids per group. Student t-test (\*\*\*\*pvalue<0.00001). Representative images are shown. **C)** Treatment of GFP+ and GFP- MM27 cells with 10nM trametinib for 72 hours *in vitro*. Cell viability was assessed by CyQuant and normalized to DMSO control. Mean ± SD (n=3). p values are based on unpaired Student's t test (\*\*\*p < 0.001). The experiment was performed three times. **D)** GFP+ and GFP- MM27 cell mitochondrial activity was assessed by flow cytometry analysis of mitotracker orange (25nM) stained cells. Data are shown as mean fluorescence intensity ± SD (n=3). Student t-test (\*\*\*\*pvalue<0.00001). The experiment was done twice. **E)** Basal oxygen consumption rate (OCR), OCR spare capacity and OCR max respiration of GFP+ and GFP- MM27 cells were assessed by Seahorse XF Cell Mito Stress. Data are mean ± SD (n=7). Student t-test (\*\*pvalue<0.01; \*\*\*\*pvalue<0.00001). The experiment was performed twice.

**Supplemental Figure 3.** *In vivo* assessment of tumorigenic and invasive properties of GFP+ and GFP- MM27 populations in immunocompromised mice.

**A)** tumor growth (at 6 weeks); **B)** latency to tumor palpability; **C)** latency to tumor resection (volume ∼0.3cm<sup>3</sup>); **D)** latency to lymph-node (LN) appearance; **E)** total number of organs with detectable metastases and average number of organs with metastases per mouse; **F)** LN volume at day100 after injection; **G)** total number of organs with at least 50% of melanoma cells and average number of organs with at least 50% of melanoma cells *per* mouse. **H)** Table summarizing days of latency to palpability, resection and LN appearance for each mouse injected with GFP+ and GFP- MM13 and MM27 cells. Data are mean ± SD (n=5). Student t test was applied to assess the significance (ns, not significant; \*pvalue< 0.05; \*\*pvalue<0.001; \*\*\*pvalue<0.0001).

**Supplemental Figure 4.** Impact of the hypoxic tumor microenvironment on MM27 slow cycling cells.

**A)** GFP+ and GFP- cell migration was assessed by transwell migration assay after 24h in normal (20%O<sub>2</sub>) and low oxygen (3%O<sub>2</sub>) incubators. Representative EVOS microscopy images of migrated cells (transwell outer surface) are reported. Data are mean±-SD (n=3). p-values are based on unpaired Student's t test (\*\*\*p < 0.0001). The experiment was performed three times. **B)** Spheroid collagen invasion in GFP+ and GFP- cells in normal (20%O<sub>2</sub>) and low oxygen (3% O<sub>2</sub>) incubators was measured as area of invasion fold change vs T0, imaged every 24h by EVOS microscopy for 48h. Data shown are the mean ± SD of 10 different spheroids per group. Student t-test (\*\*\*\*pvalue<0.00001). Representative images are shown.

**C)** *In vitro* trametinib (10nM) treatment (72h) of GFP+ and GFP- MM13 cells in normal (20%O<sub>2</sub>) and low oxygen (3%O<sub>2</sub>) incubators. Cell viability was assessed by CyQuant and normalized to DMSO control. Mean ± SD (n=3). p values are based on unpaired Student's t test (\*\*\*p < 0.001). The experiment was performed three times. **D)** Percentage of cell moving toward hypoxia, *disp*<0, for GFP+ (green) and GFP- (gray) MM27 assessed during time-lapse of migration assay in chemical controlled culture system. Data are mean ± SD (n=3). Significance for the first 16h was assessed by one-way ANOVA (p value 0.02298 and confidence level >95%). Number of trajectories analyzed (GFP+ n=1992; GFP- n=1568). **E)** Percentage of cells moving toward hypoxia, *disp*<0, for GFP+ (green) and GFP- (gray) MM13 assessed during time-lapse of invasion assay. Data are mean ± SD (n=3). Significance for the first 8h was assessed by one-way ANOVA (p value 0.03082 and confidence level >95%). Number of trajectories analyzed (GFP+ n=1136; GFP- n=481). **F)** Average speed of GFP+ and GFP- in 3D collagen invasion assay in controlled oxygen conditions (COC) and in standard oxygen conditions (SOC) were analyzed. Data are mean ± SD (n=3). Student t test was applied to assess the significance (ns, not significant; \*\*p value<0.01; \*\*\*p value<0.001). Number of trajectories analyzed (GFP+ n=2179; GFP- n=1321).

##### **Supp. Figure 5.**

**A)** Stack plots showing relative proportions of MM13 melanoma cells annotated according to G1, S, G2/M cell cycle phases (left) and KI67- and KI67+ cells (right) across Seurat clusters. **B)** UMAP visualization (upper panel) and violin plot (lower panel) illustrating expression of reference melanoma signatures (12) in GFP+ and GFP- MM13 populations. p-values were determined by Mann-Whitney U-test.

##### **Supplemental Figure 6**

**A)** Heatmap of averaged expression of QQ signature genes in PP, PQ, QP and QQ populations. **B)** Heatmap of averaged expression of growth inhibitory gene set (GSEA cancer Module\_488) in QQ, PQ, QP and PP cell populations (left panel) and in QQ primary tumor and metastatic cells (right panel). **C)** Volcano plot (left) and heatmap (right) of differentially expressed genes identified by bulk RNA-seq analysis of FACS sorted KI67- vs KI67+ populations. Two biological MM13 replicates were used for the experiment. **D)** Venn diagram and heatmap of common genes between DEGs identified in C and QQ signature. **E)** GSEA enrichment analysis of QQ signature in RNA-seq dataset of 6 RAF/MEKi treated patients before and after treatment (GEO: GSE77940) and 11+12 patients from scRNA-seq (GEO: GSE115978) before and after treatment ICI treatment. **F)** UMAP visualization (left) of MM13 melanoma dataset colored by mesenchymal metastatic signature from Karras et. Al., 2022; list of common genes among the 2 signatures (table); violin plot (right) showing expression of PRRX1 regulon in primary and metastatic QQ populations. p-values were calculated using two-tailed Mann-Whitney U-test.

**A**

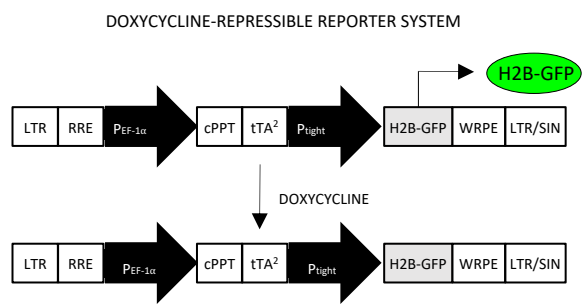

**B**

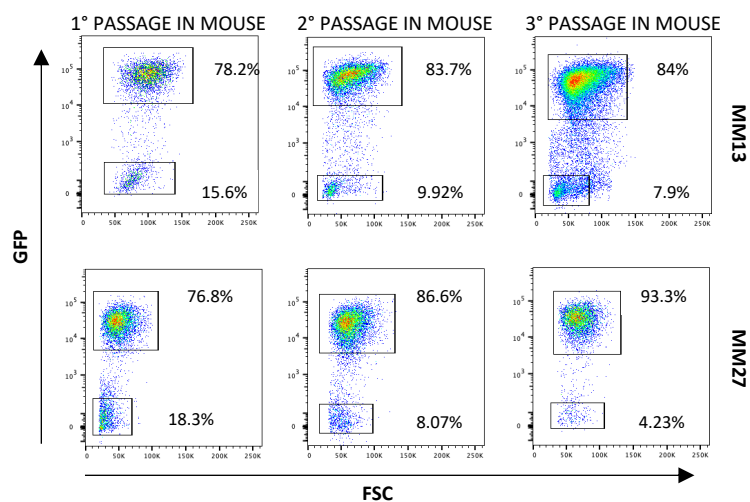

**C**

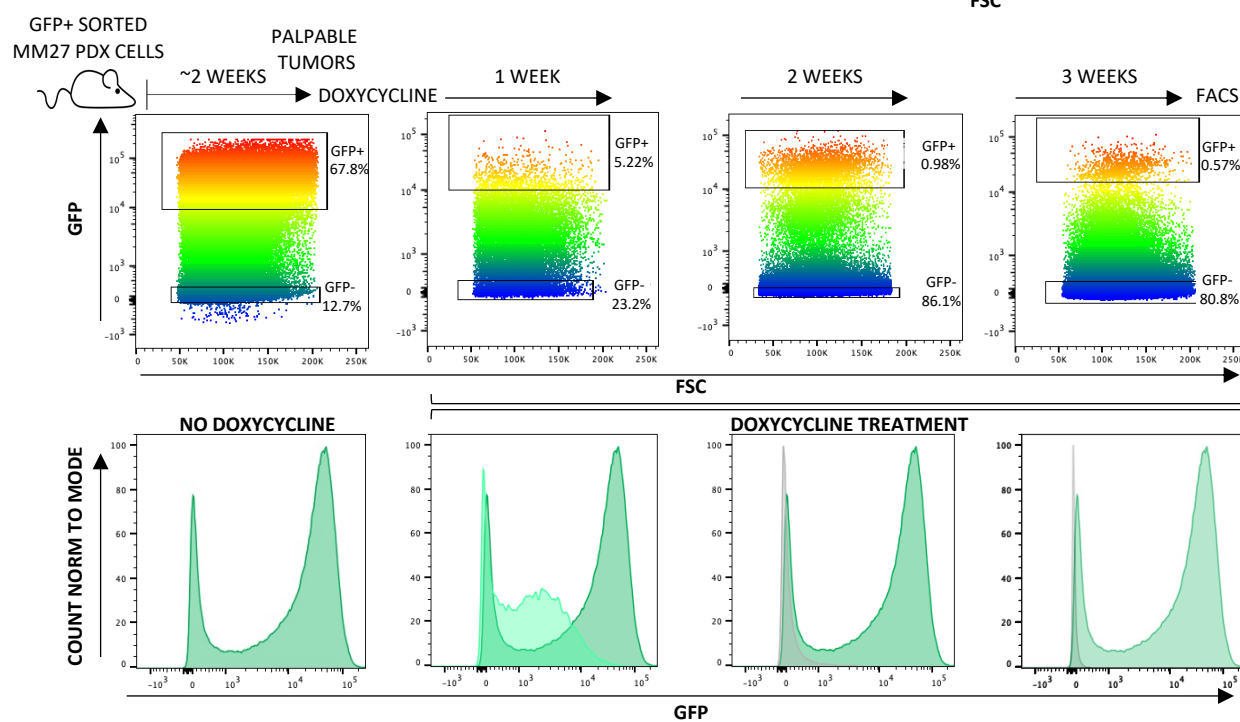

**D**

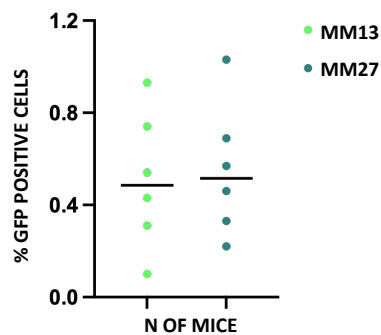

**E**

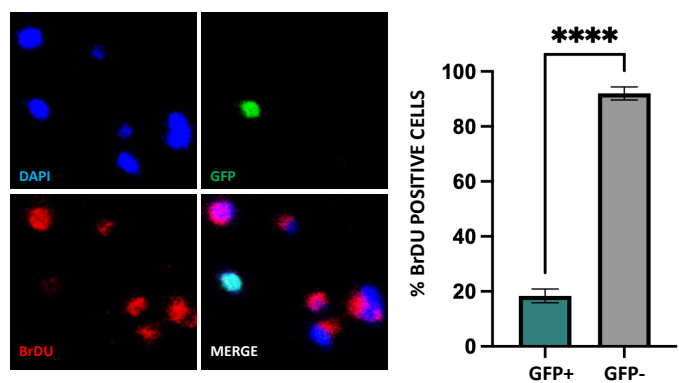

**F**

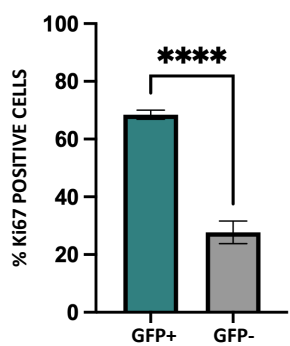

**G**

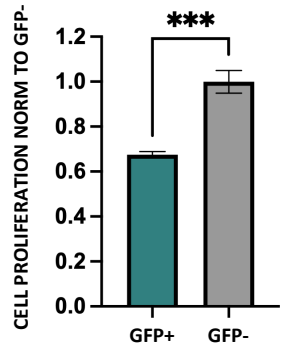

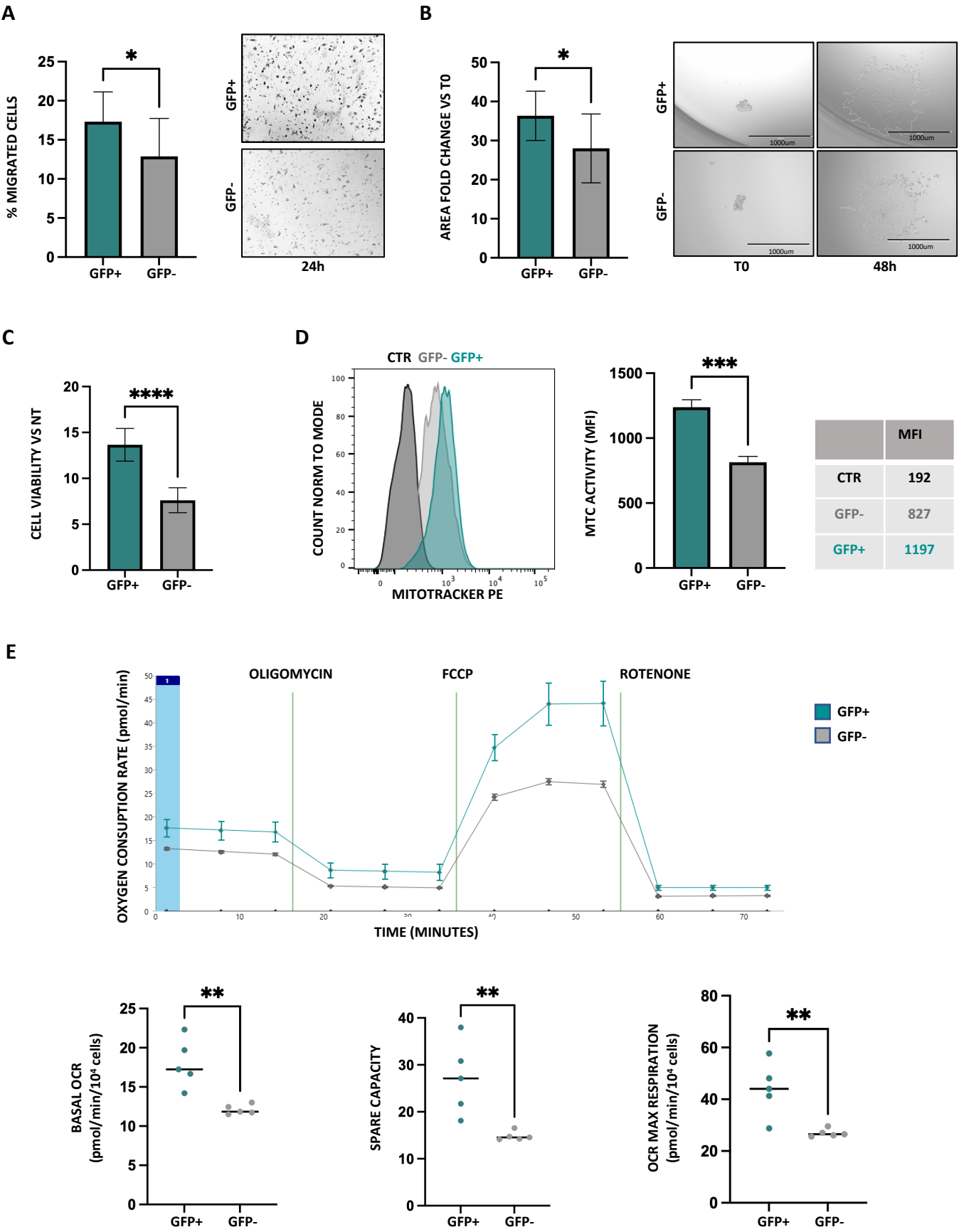

A

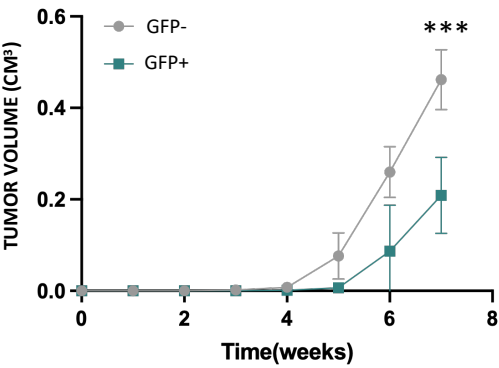

B

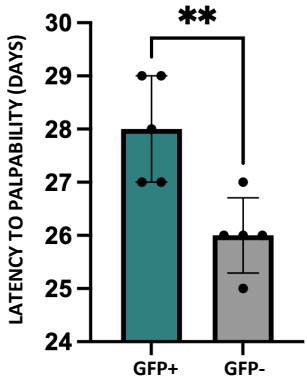

C

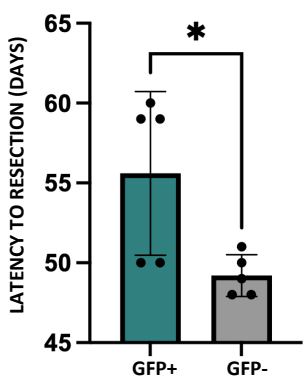

D

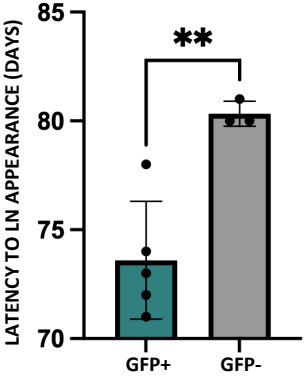

E

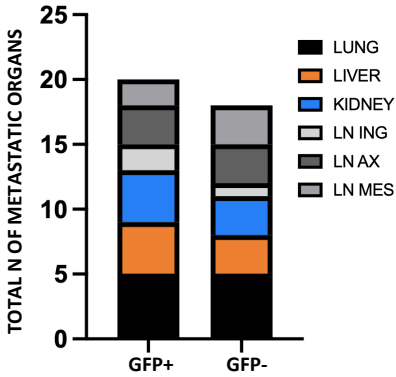

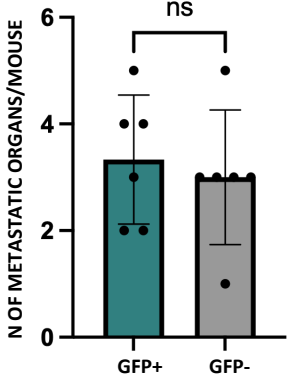

F

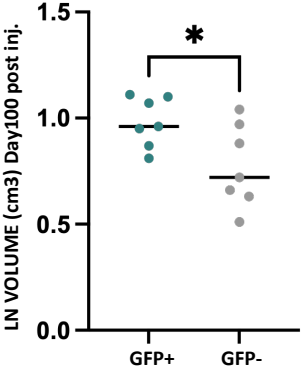

G

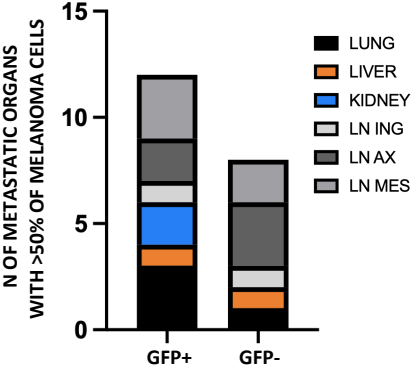

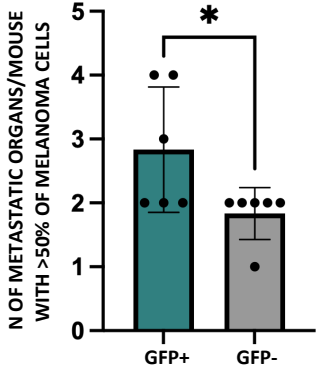

H

|  | LATENCY TO PALPABILITY | LATENCY TO RESECTION | LATENCY TO LN APPEARANCE |  | LATENCY TO PALPABILITY | LATENCY TO RESECTION | LATENCY TO LN APPEARANCE |
| --- | --- | --- | --- | --- | --- | --- | --- |
| MM13 GFP+ | DAYS |  |  | MM13 GFP- | DAYS |  |  |
| M1 | 25 | 42 | 70 | M1 | 22 | 40 | 84 |
| M2 | 25 | 50 | 70 | M2 | 22 | 44 | / |
| M3 | 25 | 42 | 77 | M3 | 22 | 39 | 79 |
| M4 | 27 | 50 | 70 | M4 | 26 | 44 | / |
| M5 | 25 | 50 | 77 | M5 | 22 | 40 | 79 |
| MM27 GFP+ | DAYS |  |  | MM27 GFP- | DAYS |  |  |
| M1 | 27 | 50 | 73 | M1 | 27 | 51 | 80 |
| M2 | 27 | 59 | 74 | M2 | 26 | 50 | 81 |
| M3 | 28 | 59 | 72 | M3 | 25 | 49 | 80 |
| M4 | 29 | 60 | 73 | M4 | 26 | 48 | / |
| M5 | 29 | 50 | 78 | M5 | 26 | 48 | / |

A

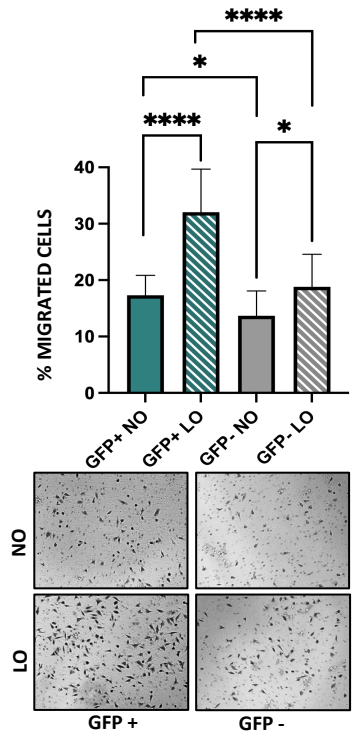

B

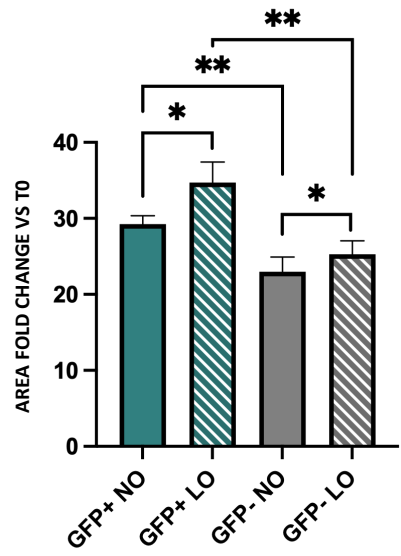

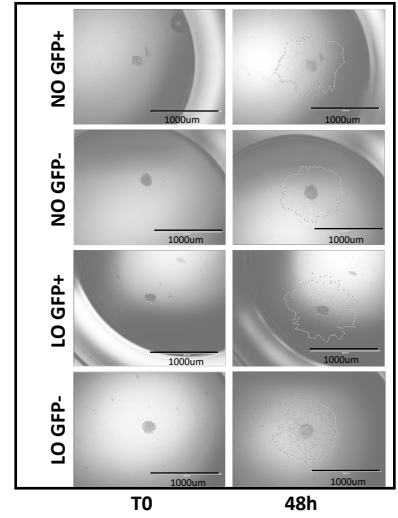

C

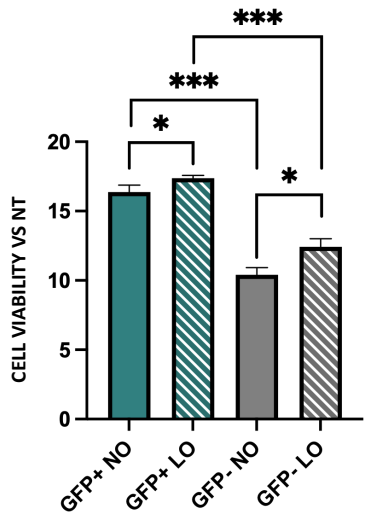

D

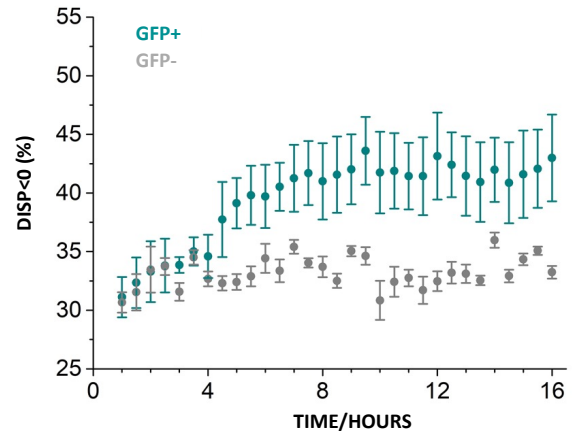

E

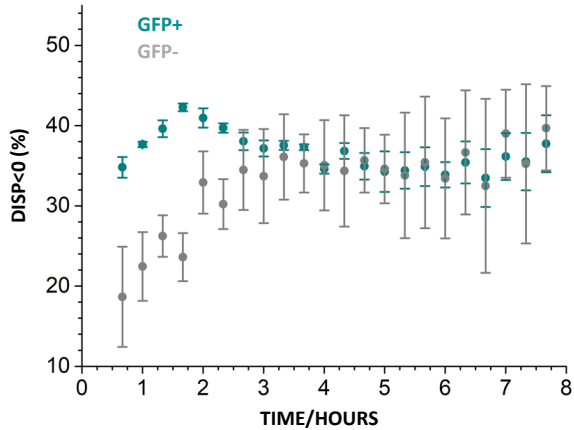

F

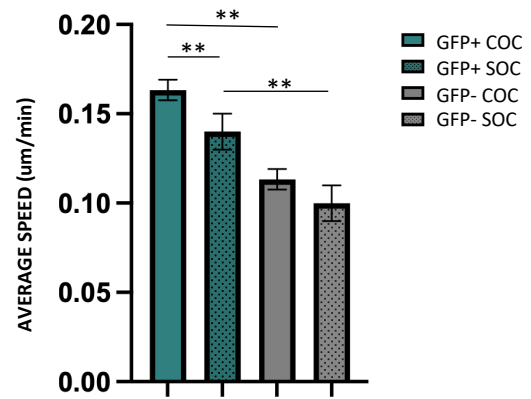

A

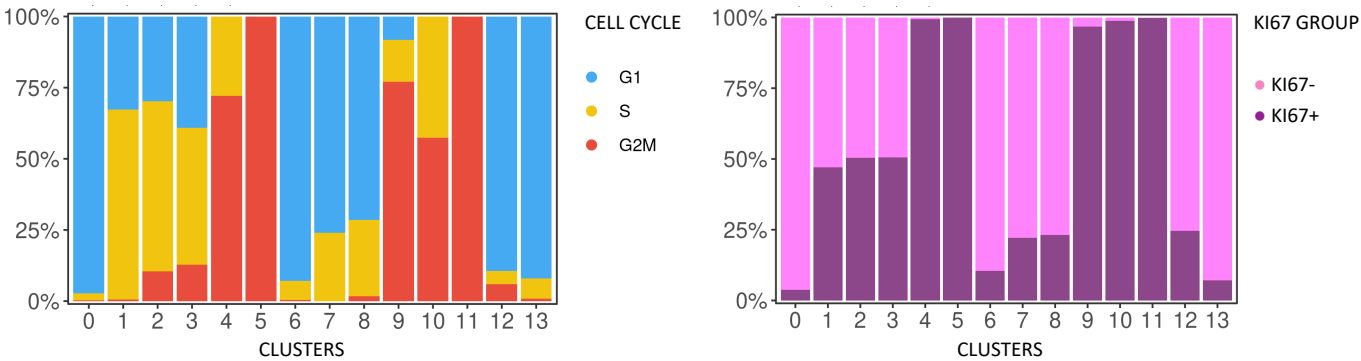

B

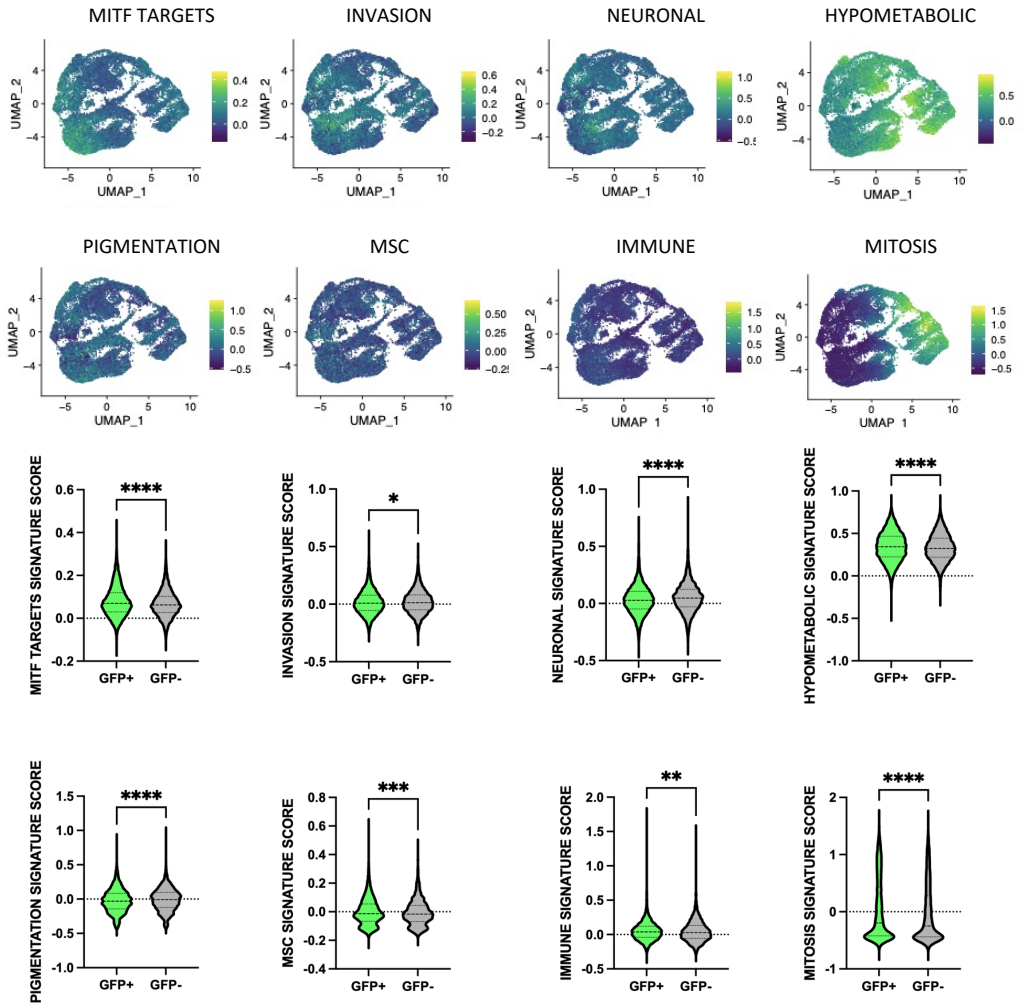

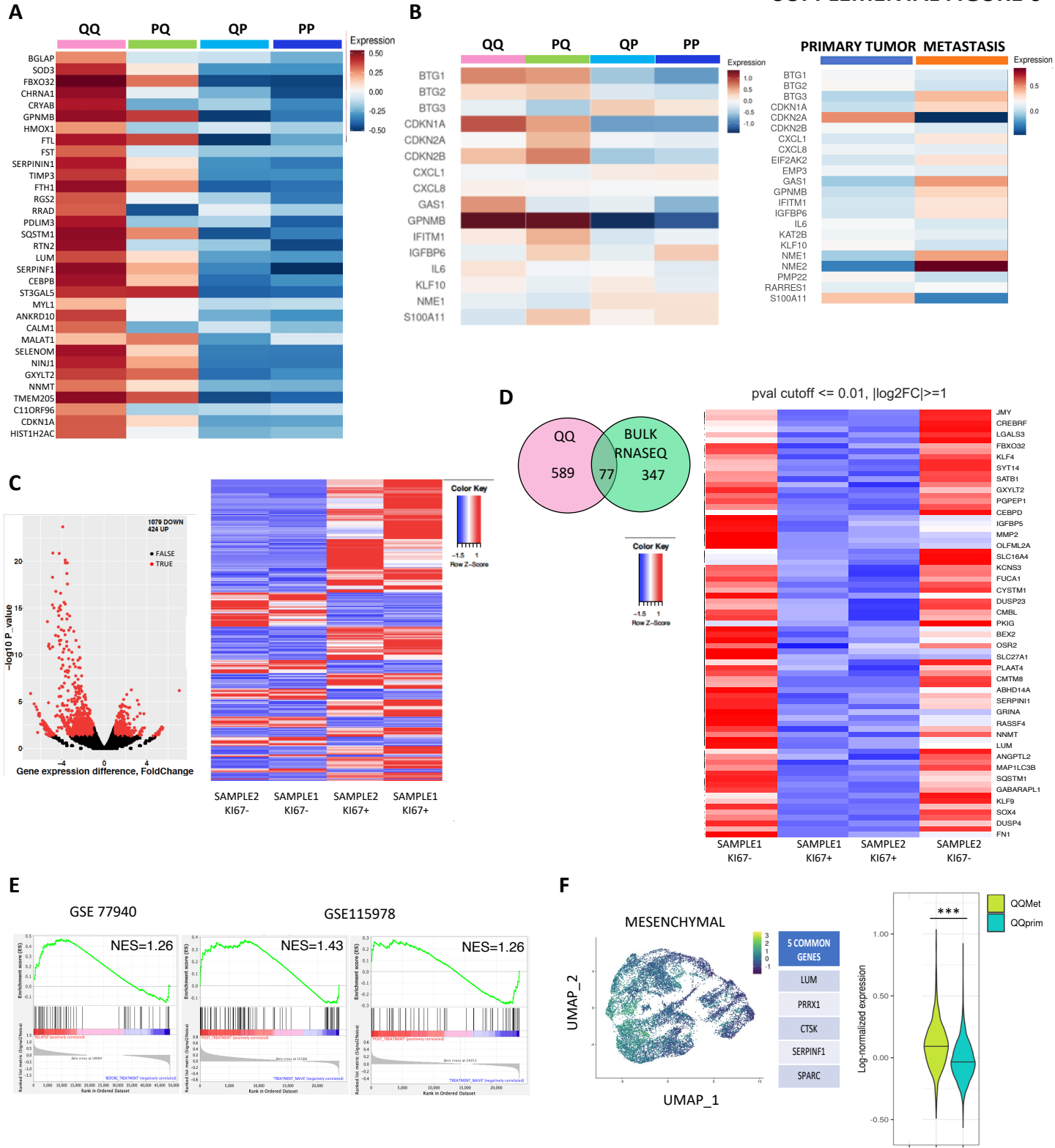

### MARKERS GENES OF QQ-PQ-QP-PP POPULATIONS

TABLE SUPP.1

| gene | avg_log2FC | p_val | p_val_adj | subtype |
| --- | --- | --- | --- | --- |
| BGLAP | 1.35764374 | 1.18E-57 | 1.67E-53 | QQ |
| SOD3 | 1.25914674 | 1.59E-186 | 2.25E-182 | QQ |
| FBXO32 | 1.22533773 | 0 | 0 | QQ |
| CHRNA1 | 1.21960208 | 0 | 0 | QQ |
| CRYAB | 1.2071042 | 4.90E-252 | 6.94E-248 | QQ |
| GPNMB | 1.17287907 | 1.35E-304 | 1.91E-300 | QQ |
| HMOX1 | 1.08259655 | 1.86E-61 | 2.63E-57 | QQ |
| FTL | 1.04505244 | 5.81E-287 | 8.22E-283 | QQ |
| FST | 1.02401482 | 8.67E-135 | 1.23E-130 | QQ |
| SERPINI1 | 0.98037376 | 1.84E-273 | 2.61E-269 | QQ |
| TIMP3 | 0.94742248 | 1.12E-241 | 1.58E-237 | QQ |
| FTH1 | 0.94237038 | 1.39E-300 | 1.97E-296 | QQ |
| RGS2 | 0.93959087 | 1.59E-160 | 2.25E-156 | QQ |
| RRAD | 0.92055988 | 1.40E-169 | 1.98E-165 | QQ |
| PDLIM3 | 0.92033744 | 0 | 0 | QQ |
| SQSTM1 | 0.89885302 | 0 | 0 | QQ |
| RTN2 | 0.8923271 | 0 | 0 | QQ |
| LUM | 0.88699656 | 9.10E-212 | 1.29E-207 | QQ |
| SERPINF1 | 0.88367051 | 0 | 0 | QQ |
| CEBPB | 0.88241196 | 0 | 0 | QQ |
| ST3GAL5 | 0.86821865 | 1.26E-238 | 1.78E-234 | QQ |
| MYL1 | 0.84903227 | 2.20E-48 | 3.11E-44 | QQ |
| ANKRD10 | 0.83193334 | 7.30E-204 | 1.03E-199 | QQ |
| CALM1 | 0.82686217 | 1.53E-104 | 2.17E-100 | QQ |
| MALAT1 | 0.82573542 | 6.28E-69 | 8.89E-65 | QQ |
| SELENOM | 0.81179178 | 1.31E-264 | 1.86E-260 | QQ |
| NINJ1 | 0.80147153 | 5.13E-265 | 7.26E-261 | QQ |
| GXYLT2 | 0.797351 | 4.88E-180 | 6.91E-176 | QQ |
| NNMT | 0.78768304 | 7.33E-109 | 1.04E-104 | QQ |
| TMEM205 | 0.78328105 | 0 | 0 | QQ |
| C11orf96 | 0.78041103 | 5.57E-90 | 7.88E-86 | QQ |
| CDKN1A | 0.7704679 | 4.03E-177 | 5.70E-173 | QQ |
| HIST1H2AC | 0.77032783 | 4.32E-121 | 6.12E-117 | QQ |
| KLHL41 | 0.74436578 | 8.35E-244 | 1.18E-239 | QQ |
| CHMP1B | 0.74411091 | 2.72E-266 | 3.85E-262 | QQ |
| MGP | 0.73862294 | 9.84E-130 | 1.39E-125 | QQ |
| ATP6V1G1 | 0.73817203 | 0 | 0 | QQ |
| YPEL3 | 0.72670973 | 5.14E-304 | 7.27E-300 | QQ |
| IGFBP5 | 0.72182391 | 3.04E-159 | 4.31E-155 | QQ |
| TGFB1 | 0.70138493 | 1.53E-105 | 2.17E-101 | QQ |
| PDXK | 0.70059828 | 2.19E-183 | 3.10E-179 | QQ |
| SOX4 | 0.69911971 | 3.18E-110 | 4.49E-106 | QQ |

|  |  |  |  |  |
| --- | --- | --- | --- | --- |
| GAS5 | 0.6927737 | 0 | 0 | QQ |
| GOLIM4 | 0.69028446 | 6.57E-68 | 9.29E-64 | QQ |
| PNRC1 | 0.67835825 | 5.11E-295 | 7.23E-291 | QQ |
| PBXIP1 | 0.67730301 | 5.62E-240 | 7.95E-236 | QQ |
| ISCU | 0.67631646 | 0 | 0 | QQ |
| EIF4A2 | 0.66715085 | 0 | 0 | QQ |
| YBX3 | 0.6669659 | 0 | 0 | QQ |
| PLCG2 | 0.66641202 | 9.53E-201 | 1.35E-196 | QQ |
| FN1 | 0.662304 | 4.39E-71 | 6.22E-67 | QQ |
| RANBP2 | 0.66160118 | 1.80E-113 | 2.54E-109 | QQ |
| ASAH1 | 0.65492438 | 1.55E-205 | 2.20E-201 | QQ |
| CEBPD | 0.64658699 | 7.52E-139 | 1.06E-134 | QQ |
| KLF9 | 0.64630681 | 1.20E-183 | 1.70E-179 | QQ |
| ZNF331 | 0.64615742 | 7.45E-48 | 1.05E-43 | QQ |
| QPRT | 0.64244951 | 9.36E-191 | 1.32E-186 | QQ |
| ID3 | 0.62605841 | 1.02E-82 | 1.44E-78 | QQ |
| NR4A2 | 0.62280308 | 1.12E-80 | 1.58E-76 | QQ |
| ARL6IP5 | 0.62160157 | 0 | 0 | QQ |
| EPB41L4A-A5 | 0.61245305 | 0 | 0 | QQ |
| JUND | 0.61105223 | 1.69E-303 | 2.40E-299 | QQ |
| PSAP | 0.60917951 | 5.45E-247 | 7.71E-243 | QQ |
| P4HA2 | 0.60769164 | 3.31E-110 | 4.69E-106 | QQ |
| GABARAP | 0.60754199 | 0 | 0 | QQ |
| CYSTM1 | 0.60665239 | 2.11E-178 | 2.99E-174 | QQ |
| MAP1LC3A | 0.59650067 | 1.95E-170 | 2.76E-166 | QQ |
| COMT | 0.59613661 | 0 | 0 | QQ |
| CTSD | 0.59600675 | 2.00E-184 | 2.83E-180 | QQ |
| MAP1LC3B | 0.59394749 | 1.61E-193 | 2.28E-189 | QQ |
| ITGA7 | 0.59343313 | 1.04E-147 | 1.46E-143 | QQ |
| NME3 | 0.59301129 | 5.75E-277 | 8.13E-273 | QQ |
| ZFAND5 | 0.58760138 | 5.19E-195 | 7.34E-191 | QQ |
| TRAPPC6A | 0.58702102 | 3.10E-174 | 4.38E-170 | QQ |
| GSN | 0.58444028 | 8.43E-304 | 1.19E-299 | QQ |
| CTSH | 0.5835127 | 7.75E-118 | 1.10E-113 | QQ |
| GLRX | 0.58270307 | 0 | 0 | QQ |
| S100A1 | 0.5803543 | 8.94E-72 | 1.27E-67 | QQ |
| MMP2 | 0.57999497 | 7.65E-151 | 1.08E-146 | QQ |
| SMIM3 | 0.57542061 | 3.32E-178 | 4.70E-174 | QQ |
| MMP11 | 0.57432346 | 1.29E-49 | 1.82E-45 | QQ |
| TIMP1 | 0.57235843 | 1.79E-81 | 2.54E-77 | QQ |
| YPEL5 | 0.56557833 | 1.27E-191 | 1.80E-187 | QQ |
| CYB5R1 | 0.56235208 | 4.76E-163 | 6.73E-159 | QQ |
| ISG15 | 0.55875955 | 2.91E-60 | 4.11E-56 | QQ |
| GMPR | 0.55788501 | 7.79E-139 | 1.10E-134 | QQ |
| GAS1 | 0.55784349 | 1.08E-149 | 1.53E-145 | QQ |
| CDKN1C | 0.55715022 | 4.15E-122 | 5.87E-118 | QQ |

|  |  |  |  |
| --- | --- | --- | --- |
| ANTXR1 | 0.55511946 | 9.46E-85 | 1.34E-80 QQ |
| PRDX2 | 0.55072023 | 1.20E-53 | 1.69E-49 QQ |
| PHLDA3 | 0.55034682 | 3.57E-216 | 5.04E-212 QQ |
| MARCKSL1 | 0.54626207 | 2.79E-140 | 3.95E-136 QQ |
| SPARC | 0.54494659 | 1.88E-184 | 2.66E-180 QQ |
| NUDT14 | 0.5427059 | 6.04E-136 | 8.55E-132 QQ |
| RAB27B | 0.53234336 | 1.71E-149 | 2.41E-145 QQ |
| PMAIP1 | 0.53035431 | 8.04E-49 | 1.14E-44 QQ |
| CTHRC1 | 0.52955669 | 1.01E-304 | 1.42E-300 QQ |
| PGM2L1 | 0.5279359 | 5.25E-92 | 7.43E-88 QQ |
| DUSP4 | 0.52760538 | 9.36E-120 | 1.32E-115 QQ |
| TSC22D3 | 0.52741287 | 2.40E-169 | 3.40E-165 QQ |
| ZMAT3 | 0.5259735 | 1.36E-172 | 1.92E-168 QQ |
| FCGRT | 0.52269579 | 2.59E-105 | 3.66E-101 QQ |
| SLC26A2 | 0.51815485 | 4.75E-120 | 6.72E-116 QQ |
| GYPC | 0.51813751 | 1.18E-183 | 1.68E-179 QQ |
| PCMTD1 | 0.51410702 | 1.00E-132 | 1.42E-128 QQ |
| ITGA10 | 0.51172005 | 9.24E-126 | 1.31E-121 QQ |
| SNHG32 | 0.50825846 | 2.13E-164 | 3.02E-160 QQ |
| KLHL24 | 0.5068998 | 5.60E-153 | 7.93E-149 QQ |
| NAMPT | 0.50573856 | 1.52E-108 | 2.16E-104 QQ |
| SNRPN | 0.50318435 | 3.47E-169 | 4.92E-165 QQ |
| SAT1 | 0.50161753 | 3.74E-114 | 5.29E-110 QQ |
| KIF13B | 0.499588 | 4.84E-120 | 6.85E-116 QQ |
| ATP6V0E1 | 0.49956032 | 0 | 0 QQ |
| ARID5B | 0.49884593 | 8.30E-138 | 1.17E-133 QQ |
| TWIST1 | 0.49883016 | 9.72E-52 | 1.38E-47 QQ |
| CHCHD10 | 0.49650899 | 1.22E-202 | 1.72E-198 QQ |
| PURB | 0.49507501 | 1.66E-97 | 2.36E-93 QQ |
| SOX9 | 0.49447268 | 9.22E-53 | 1.30E-48 QQ |
| CPEB4 | 0.49347018 | 1.30E-75 | 1.84E-71 QQ |
| CPEB2 | 0.49316281 | 4.65E-113 | 6.58E-109 QQ |
| CCPG1 | 0.49160396 | 1.35E-111 | 1.90E-107 QQ |
| HOPX | 0.48902484 | 1.89E-89 | 2.67E-85 QQ |
| STAC3 | 0.48844378 | 3.14E-22 | 4.44E-18 QQ |
| GABARAPL1 | 0.48823469 | 1.40E-144 | 1.98E-140 QQ |
| RAMP1 | 0.48517814 | 1.53E-45 | 2.17E-41 QQ |
| SLC27A1 | 0.48487168 | 2.28E-151 | 3.22E-147 QQ |
| COL11A1 | 0.48462603 | 4.69E-63 | 6.64E-59 QQ |
| SAT2 | 0.48224531 | 3.58E-204 | 5.07E-200 QQ |
| PLEKHO1 | 0.48220548 | 1.56E-170 | 2.21E-166 QQ |
| CDKN2B | 0.48201169 | 4.04E-84 | 5.72E-80 QQ |
| GNPDA1 | 0.48189672 | 1.54E-125 | 2.17E-121 QQ |
| JMY | 0.47956969 | 9.43E-91 | 1.33E-86 QQ |
| NPC2 | 0.47847042 | 1.89E-116 | 2.67E-112 QQ |
| DPP7 | 0.47778542 | 6.17E-174 | 8.74E-170 QQ |

|  |  |  |  |  |
| --- | --- | --- | --- | --- |
| CREBRF | 0.47760249 | 3.93E-106 | 5.56E-102 | QQ |
| POPDC3 | 0.4774132 | 1.13E-135 | 1.60E-131 | QQ |
| TIPARP | 0.4747854 | 3.76E-89 | 5.31E-85 | QQ |
| TMEM219 | 0.47279351 | 1.97E-242 | 2.79E-238 | QQ |
| IGBP1 | 0.47245857 | 5.66E-226 | 8.01E-222 | QQ |
| NUPR1 | 0.471605 | 4.07E-90 | 5.76E-86 | QQ |
| FABP3 | 0.4699187 | 7.55E-127 | 1.07E-122 | QQ |
| CLU | 0.46735999 | 4.96E-85 | 7.02E-81 | QQ |
| SLC1A3 | 0.46639863 | 2.40E-116 | 3.39E-112 | QQ |
| NEU1 | 0.466244 | 1.00E-157 | 1.42E-153 | QQ |
| DKK3 | 0.46497652 | 9.46E-94 | 1.34E-89 | QQ |
| FAM162A | 0.46471172 | 3.73E-273 | 5.28E-269 | QQ |
| CAP2 | 0.46318581 | 1.78E-84 | 2.52E-80 | QQ |
| TGIF1 | 0.46219619 | 1.10E-122 | 1.55E-118 | QQ |
| TP53I3 | 0.46169019 | 1.84E-116 | 2.60E-112 | QQ |
| DUSP23 | 0.46070121 | 1.30E-138 | 1.84E-134 | QQ |
| MEF2C | 0.46041898 | 7.27E-56 | 1.03E-51 | QQ |
| ETFRF1 | 0.45982633 | 7.00E-148 | 9.90E-144 | QQ |
| BRI3 | 0.45822753 | 6.23E-230 | 8.81E-226 | QQ |
| LRRN4CL | 0.45655723 | 8.00E-79 | 1.13E-74 | QQ |
| VGLL4 | 0.45651518 | 2.23E-122 | 3.16E-118 | QQ |
| JUNB | 0.45314855 | 1.43E-64 | 2.02E-60 | QQ |
| ITM2B | 0.45217921 | 1.80E-238 | 2.55E-234 | QQ |
| OLFML2A | 0.44751026 | 8.88E-85 | 1.26E-80 | QQ |
| LAPTM4A | 0.44486898 | 1.29E-140 | 1.82E-136 | QQ |
| CST3 | 0.44484015 | 2.83E-223 | 4.01E-219 | QQ |
| GRN | 0.44421419 | 2.19E-115 | 3.10E-111 | QQ |
| PLAAT3 | 0.4431157 | 2.02E-134 | 2.86E-130 | QQ |
| MXD4 | 0.44139854 | 3.22E-147 | 4.56E-143 | QQ |
| CTSF | 0.43977402 | 4.57E-86 | 6.47E-82 | QQ |
| DUSP10 | 0.43943531 | 1.19E-83 | 1.68E-79 | QQ |
| IFI27 | 0.43884807 | 2.60E-23 | 3.67E-19 | QQ |
| SPINK2 | 0.43857036 | 9.10E-92 | 1.29E-87 | QQ |
| SLC25A4 | 0.43700496 | 1.61E-106 | 2.28E-102 | QQ |
| KDM5B | 0.43573501 | 1.69E-94 | 2.39E-90 | QQ |
| MRAS | 0.43438545 | 1.07E-86 | 1.52E-82 | QQ |
| ARMCX3 | 0.43205181 | 1.78E-150 | 2.52E-146 | QQ |
| PGF | 0.43168857 | 7.67E-61 | 1.09E-56 | QQ |
| TSC22D1 | 0.43145623 | 3.39E-105 | 4.80E-101 | QQ |
| CXXC5 | 0.4304707 | 7.83E-87 | 1.11E-82 | QQ |
| SPON2 | 0.42964361 | 5.88E-117 | 8.33E-113 | QQ |
| BTG1 | 0.42862036 | 4.93E-142 | 6.97E-138 | QQ |
| PHLDB2 | 0.42705352 | 2.31E-24 | 3.26E-20 | QQ |
| SCN1B | 0.42475135 | 6.77E-105 | 9.58E-101 | QQ |
| MFSD12 | 0.42379107 | 8.49E-193 | 1.20E-188 | QQ |
| MAGED2 | 0.42227565 | 7.50E-102 | 1.06E-97 | QQ |

|  |  |  |  |
| --- | --- | --- | --- |
| PLAAT4 | 0.42081139 | 8.98E-95 | 1.27E-90 QQ |
| MPG | 0.42013678 | 1.03E-232 | 1.46E-228 QQ |
| RRM2B | 0.4192628 | 8.57E-74 | 1.21E-69 QQ |
| IER5L | 0.41867157 | 1.29E-53 | 1.83E-49 QQ |
| TERF2IP | 0.41853648 | 8.71E-158 | 1.23E-153 QQ |
| LRP1 | 0.41768373 | 3.71E-92 | 5.25E-88 QQ |
| GPX4 | 0.41712053 | 8.08E-141 | 1.14E-136 QQ |
| KLF4 | 0.41674792 | 3.46E-82 | 4.89E-78 QQ |
| IFI6 | 0.41646365 | 3.99E-17 | 5.64E-13 QQ |
| CMTM8 | 0.41488987 | 7.07E-103 | 1.00E-98 QQ |
| EIF4EBP3 | 0.41470921 | 9.61E-93 | 1.36E-88 QQ |
| ALKBH7 | 0.41372985 | 1.86E-185 | 2.63E-181 QQ |
| SBDS | 0.41329411 | 6.62E-179 | 9.36E-175 QQ |
| ARMCX1 | 0.41303732 | 1.19E-122 | 1.68E-118 QQ |
| SCG2 | 0.41288975 | 9.05E-28 | 1.28E-23 QQ |
| HSD17B14 | 0.41253205 | 4.78E-113 | 6.76E-109 QQ |
| TMEM59 | 0.41224201 | 2.72E-194 | 3.85E-190 QQ |
| SKIL | 0.41218098 | 1.67E-37 | 2.36E-33 QQ |
| TIMP2 | 0.41180903 | 1.29E-161 | 1.83E-157 QQ |
| NEAT1 | 0.40914671 | 3.24E-38 | 4.59E-34 QQ |
| CREM | 0.40694937 | 1.22E-64 | 1.72E-60 QQ |
| HTRA1 | 0.40636933 | 1.16E-126 | 1.64E-122 QQ |
| PLD3 | 0.40512627 | 8.24E-131 | 1.17E-126 QQ |
| SIX1 | 0.40404662 | 1.26E-87 | 1.78E-83 QQ |
| GEM | 0.40118479 | 2.55E-55 | 3.61E-51 QQ |
| CTSK | 0.40092818 | 1.31E-24 | 1.85E-20 QQ |
| BEX2 | 0.4006644 | 1.23E-105 | 1.75E-101 QQ |
| RASSF4 | 0.40028122 | 1.63E-53 | 2.31E-49 QQ |
| RETREG1 | 0.39852653 | 5.00E-75 | 7.07E-71 QQ |
| CHD2 | 0.39791614 | 1.52E-72 | 2.15E-68 QQ |
| DMPK | 0.39784025 | 4.37E-68 | 6.19E-64 QQ |
| TFPT | 0.39766376 | 8.57E-149 | 1.21E-144 QQ |
| BHLHE40 | 0.39695523 | 1.60E-43 | 2.27E-39 QQ |
| ARRDC3 | 0.39672962 | 7.12E-54 | 1.01E-49 QQ |
| C18orf32 | 0.39652643 | 4.19E-117 | 5.93E-113 QQ |
| RASSF8 | 0.39646157 | 4.08E-43 | 5.77E-39 QQ |
| SARAF | 0.39464365 | 4.35E-171 | 6.15E-167 QQ |
| GBF1 | 0.39447441 | 6.66E-50 | 9.43E-46 QQ |
| NRN1 | 0.39258313 | 2.66E-125 | 3.76E-121 QQ |
| GADD45B | 0.39220471 | 1.96E-11 | 2.77E-07 QQ |
| CAMLG | 0.39158964 | 1.86E-186 | 2.64E-182 QQ |
| IDS | 0.38814212 | 1.19E-98 | 1.69E-94 QQ |
| TOB1 | 0.38671957 | 3.90E-72 | 5.52E-68 QQ |
| GUK1 | 0.3866655 | 2.04E-229 | 2.89E-225 QQ |
| MEGF9 | 0.38598693 | 8.49E-48 | 1.20E-43 QQ |
| IDI1 | 0.38592178 | 8.19E-30 | 1.16E-25 QQ |

|  |  |  |  |  |
| --- | --- | --- | --- | --- |
| CIRBP | 0.38461449 | 1.45E-214 | 2.05E-210 | QQ |
| BEX1 | 0.38389358 | 9.89E-23 | 1.40E-18 | QQ |
| SNHG29 | 0.38381946 | 6.10E-302 | 8.63E-298 | QQ |
| SH3BGRL | 0.38275909 | 1.63E-148 | 2.30E-144 | QQ |
| LRMDA | 0.38189734 | 6.64E-79 | 9.39E-75 | QQ |
| ATP10D | 0.38111631 | 8.33E-68 | 1.18E-63 | QQ |
| GPM6B | 0.38021364 | 5.26E-49 | 7.45E-45 | QQ |
| HSPB2 | 0.38017482 | 2.11E-86 | 2.99E-82 | QQ |
| C1orf122 | 0.37940103 | 7.63E-103 | 1.08E-98 | QQ |
| BORCS7 | 0.37877636 | 8.25E-106 | 1.17E-101 | QQ |
| MPV17 | 0.37874744 | 1.44E-98 | 2.04E-94 | QQ |
| DAB2 | 0.3783804 | 1.83E-62 | 2.59E-58 | QQ |
| MAP1A | 0.37838027 | 1.06E-60 | 1.50E-56 | QQ |
| RABAC1 | 0.37757318 | 3.89E-174 | 5.51E-170 | QQ |
| RHOQ | 0.37715561 | 1.97E-74 | 2.79E-70 | QQ |
| UBE2H | 0.37607639 | 7.88E-68 | 1.11E-63 | QQ |
| ZBTB20 | 0.37600429 | 1.17E-58 | 1.65E-54 | QQ |
| ABHD14B | 0.37566447 | 1.74E-127 | 2.46E-123 | QQ |
| UGCG | 0.37564747 | 2.78E-66 | 3.94E-62 | QQ |
| NFKBIZ | 0.37406153 | 3.59E-63 | 5.08E-59 | QQ |
| SLC25A6 | 0.37344323 | 0 | 0 | QQ |
| NEXN | 0.37201298 | 1.33E-54 | 1.88E-50 | QQ |
| GABARAPL2 | 0.36959392 | 9.82E-169 | 1.39E-164 | QQ |
| VEGFA | 0.36932074 | 1.90E-64 | 2.69E-60 | QQ |
| FXR1 | 0.36835881 | 7.46E-129 | 1.05E-124 | QQ |
| HIGD2A | 0.36774836 | 2.62E-207 | 3.71E-203 | QQ |
| PHYH | 0.36770844 | 2.52E-80 | 3.57E-76 | QQ |
| CHCHD6 | 0.36766818 | 1.30E-79 | 1.84E-75 | QQ |
| PLK2 | 0.36747699 | 6.14E-73 | 8.68E-69 | QQ |
| TPT1 | 0.36729082 | 0 | 0 | QQ |
| FOXD3-AS1 | 0.36684204 | 7.33E-80 | 1.04E-75 | QQ |
| DAAM2 | 0.36615765 | 6.90E-73 | 9.77E-69 | QQ |
| RPS4Y1 | 0.36532086 | 0 | 0 | QQ |
| DUSP13 | 0.36530298 | 2.65E-68 | 3.75E-64 | QQ |
| ABHD2 | 0.36448525 | 5.02E-43 | 7.10E-39 | QQ |
| STN1 | 0.36400941 | 4.54E-105 | 6.42E-101 | QQ |
| MXI1 | 0.36292926 | 1.48E-88 | 2.09E-84 | QQ |
| HBP1 | 0.36292463 | 2.26E-83 | 3.20E-79 | QQ |
| ECH1 | 0.35993356 | 3.56E-79 | 5.03E-75 | QQ |
| ELL2 | 0.35922293 | 1.29E-53 | 1.82E-49 | QQ |
| DAAM1 | 0.35884468 | 1.13E-48 | 1.59E-44 | QQ |
| SLC6A8 | 0.35869862 | 5.84E-63 | 8.26E-59 | QQ |
| POLD4 | 0.35678027 | 1.75E-96 | 2.48E-92 | QQ |
| DYRK3 | 0.35459554 | 6.40E-38 | 9.05E-34 | QQ |
| FUCA1 | 0.35431532 | 2.07E-114 | 2.93E-110 | QQ |
| GCC2 | 0.3541861 | 1.81E-66 | 2.56E-62 | QQ |

|  |  |  |  |  |
| --- | --- | --- | --- | --- |
| CD63 | 0.35273221 | 0 | 0 | QQ |
| SYNGR1 | 0.35272286 | 5.52E-64 | 7.81E-60 | QQ |
| SCAND1 | 0.35129324 | 1.53E-122 | 2.17E-118 | QQ |
| TRIP6 | 0.35123016 | 6.14E-83 | 8.68E-79 | QQ |
| FDXR | 0.35025204 | 1.11E-34 | 1.58E-30 | QQ |
| CCDC85B | 0.34992646 | 1.90E-118 | 2.69E-114 | QQ |
| PJA2 | 0.34953767 | 4.84E-70 | 6.85E-66 | QQ |
| DUBR | 0.34949507 | 1.58E-66 | 2.24E-62 | QQ |
| CCNL1 | 0.34920461 | 2.59E-58 | 3.67E-54 | QQ |
| ATP6V1F | 0.34890155 | 8.61E-120 | 1.22E-115 | QQ |
| TCF12 | 0.34855772 | 2.46E-46 | 3.48E-42 | QQ |
| MEGF10 | 0.34855702 | 1.25E-43 | 1.77E-39 | QQ |
| BBX | 0.34847084 | 3.39E-118 | 4.80E-114 | QQ |
| MRPL34 | 0.34834932 | 9.95E-147 | 1.41E-142 | QQ |
| SLC29A1 | 0.34767164 | 7.34E-19 | 1.04E-14 | QQ |
| AVPI1 | 0.34692123 | 6.79E-74 | 9.61E-70 | QQ |
| LEPROT | 0.34606484 | 5.87E-83 | 8.31E-79 | QQ |
| RPL3 | 0.3458361 | 0 | 0 | QQ |
| PRRX1 | 0.34541196 | 1.86E-59 | 2.63E-55 | QQ |
| SRP14 | 0.34540217 | 0 | 0 | QQ |
| LINC02397 | 0.34451796 | 7.20E-98 | 1.02E-93 | QQ |
| CHMP2A | 0.34372927 | 3.45E-187 | 4.89E-183 | QQ |
| FAM174A | 0.34334059 | 1.31E-70 | 1.85E-66 | QQ |
| HSPA1A | 0.34332108 | 2.00E-08 | 0.00028353 | QQ |
| CELF2 | 0.3431383 | 9.25E-59 | 1.31E-54 | QQ |
| FGF1 | 0.34243199 | 3.57E-67 | 5.05E-63 | QQ |
| CSRP3 | 0.34084799 | 2.60E-43 | 3.68E-39 | QQ |
| IFITM3 | 0.33915202 | 2.70E-174 | 3.82E-170 | QQ |
| LGALS3 | 0.33908386 | 1.61E-216 | 2.28E-212 | QQ |
| CMBL | 0.33737392 | 5.47E-70 | 7.73E-66 | QQ |
| DNASE2 | 0.33726522 | 4.94E-65 | 6.99E-61 | QQ |
| PCDH9 | 0.33641978 | 1.90E-35 | 2.69E-31 | QQ |
| ZNF622 | 0.33628205 | 1.80E-66 | 2.54E-62 | QQ |
| GTF2B | 0.33559873 | 1.26E-71 | 1.79E-67 | QQ |
| SNHG14 | 0.3355642 | 2.42E-48 | 3.43E-44 | QQ |
| C2 | 0.33500812 | 6.26E-70 | 8.85E-66 | QQ |
| YIPF6 | 0.33467241 | 8.94E-71 | 1.26E-66 | QQ |
| C9orf16 | 0.3346169 | 5.14E-131 | 7.27E-127 | QQ |
| NDUFB5 | 0.33422341 | 5.46E-140 | 7.73E-136 | QQ |
| FAM210B | 0.33400912 | 8.00E-79 | 1.13E-74 | QQ |
| TMEM47 | 0.33386739 | 5.19E-48 | 7.34E-44 | QQ |
| TCEAL4 | 0.33312518 | 9.34E-167 | 1.32E-162 | QQ |
| VAMP2 | 0.33294056 | 6.03E-93 | 8.53E-89 | QQ |
| RNASEK | 0.33264981 | 1.72E-171 | 2.44E-167 | QQ |
| C11orf1 | 0.33130004 | 1.88E-55 | 2.66E-51 | QQ |
| SCX | 0.33104825 | 6.42E-61 | 9.08E-57 | QQ |

|  |  |  |  |
| --- | --- | --- | --- |
| RHOD | 0.33087692 | 3.79E-64 | 5.36E-60 QQ |
| C15orf48 | 0.3304741 | 6.53E-24 | 9.24E-20 QQ |
| CDH15 | 0.33005929 | 1.52E-49 | 2.16E-45 QQ |
| LAMB2 | 0.32975452 | 1.10E-48 | 1.56E-44 QQ |
| PCDH7 | 0.32971743 | 1.07E-40 | 1.51E-36 QQ |
| PYCARD | 0.3295821 | 1.65E-109 | 2.33E-105 QQ |
| TMEM150A | 0.32955422 | 1.48E-78 | 2.09E-74 QQ |
| CIR1 | 0.32937806 | 7.57E-88 | 1.07E-83 QQ |
| EAPP | 0.32924958 | 3.13E-114 | 4.43E-110 QQ |
| CTSA | 0.32918705 | 7.64E-73 | 1.08E-68 QQ |
| COX7A2L | 0.32866449 | 1.12E-193 | 1.58E-189 QQ |
| SLC2A3 | 0.32772386 | 3.33E-35 | 4.71E-31 QQ |
| SNED1 | 0.32728569 | 8.48E-46 | 1.20E-41 QQ |
| LAMTOR4 | 0.32724152 | 8.15E-172 | 1.15E-167 QQ |
| MAFF | 0.32684095 | 3.07E-47 | 4.35E-43 QQ |
| SVIL | 0.3262105 | 2.08E-49 | 2.95E-45 QQ |
| MGST1 | 0.32598443 | 8.75E-38 | 1.24E-33 QQ |
| TOMM20 | 0.32489514 | 3.12E-246 | 4.42E-242 QQ |
| CHPF | 0.32472297 | 8.85E-64 | 1.25E-59 QQ |
| BNIP3L | 0.32271293 | 1.10E-93 | 1.55E-89 QQ |
| AC069360.1 | 0.32250576 | 4.23E-50 | 5.98E-46 QQ |
| PLCE1 | 0.32241597 | 6.06E-30 | 8.58E-26 QQ |
| SLC66A1L | 0.32221492 | 5.48E-59 | 7.75E-55 QQ |
| PLEKHA1 | 0.32200263 | 7.71E-56 | 1.09E-51 QQ |
| PIGP | 0.32170761 | 1.04E-65 | 1.47E-61 QQ |
| CCND3 | 0.32128492 | 1.59E-68 | 2.24E-64 QQ |
| EID1 | 0.32115288 | 2.98E-272 | 4.21E-268 QQ |
| RAB9A | 0.32090262 | 1.02E-69 | 1.45E-65 QQ |
| FAM49B | 0.32057255 | 1.67E-46 | 2.36E-42 QQ |
| NBDY | 0.32035794 | 1.61E-145 | 2.28E-141 QQ |
| CALCOCO1 | 0.32031048 | 8.95E-55 | 1.27E-50 QQ |
| SLC38A6 | 0.32019585 | 5.41E-79 | 7.65E-75 QQ |
| CFH | 0.32018171 | 4.94E-59 | 6.99E-55 QQ |
| AFF4 | 0.32016273 | 1.14E-54 | 1.61E-50 QQ |
| GZF1 | 0.31976009 | 1.69E-26 | 2.39E-22 QQ |
| CITED2 | 0.31943621 | 4.44E-54 | 6.28E-50 QQ |
| KIFAP3 | 0.31942736 | 6.26E-59 | 8.86E-55 QQ |
| WDR45 | 0.31910448 | 1.98E-74 | 2.80E-70 QQ |
| C4orf3 | 0.31787154 | 2.39E-96 | 3.38E-92 QQ |
| PON2 | 0.31737373 | 1.57E-78 | 2.22E-74 QQ |
| P4HTM | 0.3171964 | 4.35E-59 | 6.15E-55 QQ |
| AC100810.1 | 0.31718554 | 2.95E-71 | 4.18E-67 QQ |
| MFGE8 | 0.3170971 | 3.93E-23 | 5.56E-19 QQ |
| ZFP36L2 | 0.31674906 | 1.16E-54 | 1.64E-50 QQ |
| UBC | 0.31663012 | 4.68E-124 | 6.62E-120 QQ |
| ANXA4 | 0.3163457 | 3.47E-84 | 4.91E-80 QQ |

|  |  |  |  |
| --- | --- | --- | --- |
| TMEM108 | 0.31608937 | 2.34E-55 | 3.31E-51 QQ |
| TM2D1 | 0.31589833 | 1.08E-68 | 1.53E-64 QQ |
| NAT14 | 0.31531859 | 1.53E-83 | 2.17E-79 QQ |
| AKAP13 | 0.31484983 | 1.26E-46 | 1.78E-42 QQ |
| JARID2 | 0.31445545 | 6.31E-37 | 8.93E-33 QQ |
| WBP2 | 0.3143636 | 6.48E-72 | 9.16E-68 QQ |
| PGPEP1 | 0.31434207 | 2.64E-66 | 3.73E-62 QQ |
| PLA2G4C | 0.31344932 | 1.95E-75 | 2.76E-71 QQ |
| CPQ | 0.31316098 | 4.74E-46 | 6.71E-42 QQ |
| ENO3 | 0.31305907 | 7.51E-08 | 0.00106288 QQ |
| DNAJB9 | 0.31260355 | 1.34E-36 | 1.90E-32 QQ |
| DNAJB4 | 0.31255815 | 1.01E-16 | 1.43E-12 QQ |
| ABHD14A | 0.31164938 | 6.09E-91 | 8.62E-87 QQ |
| LYST | 0.31141871 | 6.81E-47 | 9.63E-43 QQ |
| TMSB4X | 0.31095458 | 1.57E-49 | 2.22E-45 QQ |
| BEX4 | 0.3106765 | 4.62E-111 | 6.54E-107 QQ |
| ANKRD28 | 0.30971611 | 4.84E-52 | 6.85E-48 QQ |
| EIF3E | 0.30952648 | 7.67E-246 | 1.09E-241 QQ |
| FNDC3A | 0.30925144 | 3.02E-42 | 4.27E-38 QQ |
| TNFSF4 | 0.30924277 | 3.78E-74 | 5.35E-70 QQ |
| BCL2L11 | 0.30905181 | 7.64E-29 | 1.08E-24 QQ |
| ZNF106 | 0.3085605 | 5.92E-13 | 8.37E-09 QQ |
| ZNF524 | 0.30818216 | 2.49E-45 | 3.52E-41 QQ |
| SNHG8 | 0.30800153 | 1.40E-80 | 1.98E-76 QQ |
| FHIT | 0.30743735 | 4.99E-49 | 7.06E-45 QQ |
| MAD1L1 | 0.30700657 | 2.90E-60 | 4.10E-56 QQ |
| IDI2-AS1 | 0.30696218 | 1.87E-103 | 2.64E-99 QQ |
| GBP2 | 0.30670404 | 3.71E-43 | 5.24E-39 QQ |
| ATP6V0B | 0.30662051 | 6.65E-62 | 9.41E-58 QQ |
| CRBN | 0.30577681 | 8.23E-74 | 1.16E-69 QQ |
| SSC5D | 0.30456217 | 3.01E-51 | 4.25E-47 QQ |
| KLC1 | 0.3044525 | 3.26E-69 | 4.62E-65 QQ |
| TTC3 | 0.30419292 | 9.25E-51 | 1.31E-46 QQ |
| LAMTOR5 | 0.30398873 | 9.63E-224 | 1.36E-219 QQ |
| SRPX2 | 0.30336218 | 4.32E-30 | 6.12E-26 QQ |
| NOP53 | 0.30335567 | 2.61E-163 | 3.69E-159 QQ |
| AC090204.1 | 0.30329629 | 2.10E-51 | 2.98E-47 QQ |
| CHGB | 0.30279718 | 3.82E-38 | 5.41E-34 QQ |
| BLOC1S2 | 0.30228025 | 2.88E-69 | 4.07E-65 QQ |
| GNG7 | 0.30181535 | 1.48E-75 | 2.10E-71 QQ |
| NFKBIA | 0.30145671 | 7.71E-55 | 1.09E-50 QQ |
| SCO2 | 0.3005824 | 2.82E-56 | 3.99E-52 QQ |
| NR4A3 | 0.30016359 | 1.02E-46 | 1.45E-42 QQ |
| NSMCE3 | 0.29983193 | 1.61E-62 | 2.28E-58 QQ |
| TMEM134 | 0.29969417 | 1.41E-76 | 1.99E-72 QQ |
| LGALS3BP | 0.29914454 | 1.42E-72 | 2.00E-68 QQ |

|  |  |  |  |
| --- | --- | --- | --- |
| SCN5A | 0.29909847 | 9.64E-30 | 1.36E-25 QQ |
| GSTK1 | 0.29883814 | 6.52E-73 | 9.22E-69 QQ |
| POLR3GL | 0.29701784 | 1.30E-62 | 1.83E-58 QQ |
| CPE | 0.29696489 | 1.62E-42 | 2.29E-38 QQ |
| SLC35F5 | 0.2967505 | 2.77E-30 | 3.92E-26 QQ |
| TFAP2A | 0.29674465 | 7.73E-54 | 1.09E-49 QQ |
| NFE2L2 | 0.29589697 | 1.83E-56 | 2.59E-52 QQ |
| RNASET2 | 0.29496323 | 1.03E-58 | 1.45E-54 QQ |
| PEBP1 | 0.29491842 | 7.93E-161 | 1.12E-156 QQ |
| DBP | 0.29366361 | 1.89E-71 | 2.68E-67 QQ |
| OCEL1 | 0.29251775 | 9.40E-60 | 1.33E-55 QQ |
| SUPT4H1 | 0.29239561 | 1.99E-113 | 2.81E-109 QQ |
| MPHOSPH8 | 0.2921917 | 9.65E-73 | 1.36E-68 QQ |
| TAGLN | 0.29177819 | 6.00E-08 | 0.00084928 QQ |
| VPS28 | 0.29164995 | 2.07E-151 | 2.92E-147 QQ |
| JSRP1 | 0.2905097 | 1.41E-18 | 1.99E-14 QQ |
| AOPEP | 0.29041876 | 4.78E-22 | 6.76E-18 QQ |
| RAB13 | 0.29010585 | 6.77E-90 | 9.58E-86 QQ |
| SLC22A18 | 0.28997883 | 6.86E-50 | 9.70E-46 QQ |
| HOOK2 | 0.28976253 | 6.58E-49 | 9.31E-45 QQ |
| NACA | 0.28924644 | 0 | 0 QQ |
| PEG10 | 0.28841574 | 7.96E-39 | 1.13E-34 QQ |
| H19 | 0.28818058 | 8.62E-24 | 1.22E-19 QQ |
| SGCA | 0.28775499 | 1.65E-21 | 2.33E-17 QQ |
| GLI4 | 0.28701215 | 8.58E-51 | 1.21E-46 QQ |
| FBXL15 | 0.28661926 | 4.04E-59 | 5.72E-55 QQ |
| JMJD1C | 0.28633652 | 1.35E-33 | 1.92E-29 QQ |
| SYF2 | 0.28612253 | 1.82E-85 | 2.58E-81 QQ |
| PROS1 | 0.28573413 | 1.91E-36 | 2.71E-32 QQ |
| N4BP2L2 | 0.28543779 | 1.47E-56 | 2.07E-52 QQ |
| CHMP5 | 0.28535306 | 3.85E-89 | 5.44E-85 QQ |
| PFN2 | 0.28527429 | 5.49E-74 | 7.77E-70 QQ |
| ATP6V1E1 | 0.28524395 | 5.47E-54 | 7.74E-50 QQ |
| GLUL | 0.28504686 | 1.05E-76 | 1.49E-72 QQ |
| OSR2 | 0.28441656 | 5.66E-21 | 8.01E-17 QQ |
| CD46 | 0.28436782 | 8.07E-56 | 1.14E-51 QQ |
| LDLRAD3 | 0.2840743 | 7.26E-28 | 1.03E-23 QQ |
| AHR | 0.28253252 | 5.49E-23 | 7.77E-19 QQ |
| ORAI3 | 0.2820064 | 7.21E-55 | 1.02E-50 QQ |
| SPRY1 | 0.28142305 | 7.27E-26 | 1.03E-21 QQ |
| TSPAN5 | 0.28129619 | 4.92E-47 | 6.97E-43 QQ |
| RPS4X | 0.28113606 | 0 | 0 QQ |
| TPP1 | 0.28078126 | 3.86E-38 | 5.46E-34 QQ |
| NEB | 0.28058637 | 1.07E-35 | 1.52E-31 QQ |
| SCRN2 | 0.28055779 | 3.57E-51 | 5.05E-47 QQ |
| MSRB2 | 0.2804717 | 7.09E-60 | 1.00E-55 QQ |

|  |  |  |  |
| --- | --- | --- | --- |
| CADPS | 0.28036233 | 6.49E-45 | 9.18E-41 QQ |
| TMEM163 | 0.28025126 | 1.06E-48 | 1.50E-44 QQ |
| AZIN2 | 0.27995188 | 6.94E-42 | 9.82E-38 QQ |
| PKIG | 0.27968445 | 1.31E-38 | 1.86E-34 QQ |
| DYRK1B | 0.27967104 | 7.03E-59 | 9.95E-55 QQ |
| SLC25A23 | 0.27870525 | 3.94E-57 | 5.57E-53 QQ |
| TMED4 | 0.27795536 | 6.92E-58 | 9.79E-54 QQ |
| UBE2B | 0.27781769 | 2.65E-100 | 3.76E-96 QQ |
| TMEM147 | 0.27745348 | 1.29E-93 | 1.82E-89 QQ |
| RHBDF1 | 0.27738208 | 1.10E-38 | 1.55E-34 QQ |
| COL18A1 | 0.27725554 | 4.75E-30 | 6.72E-26 QQ |
| FAM177A1 | 0.27721096 | 2.33E-45 | 3.29E-41 QQ |
| MAFG | 0.27668863 | 1.52E-27 | 2.16E-23 QQ |
| ZNF32 | 0.27656979 | 2.03E-49 | 2.87E-45 QQ |
| DUSP14 | 0.2757221 | 2.41E-36 | 3.41E-32 QQ |
| TSPAN6 | 0.27492886 | 6.97E-38 | 9.86E-34 QQ |
| AP1S2 | 0.27480439 | 1.40E-65 | 1.99E-61 QQ |
| CHMP2B | 0.27400094 | 2.13E-57 | 3.01E-53 QQ |
| CERT1 | 0.27313812 | 1.79E-31 | 2.53E-27 QQ |
| SURF1 | 0.27284027 | 1.36E-66 | 1.93E-62 QQ |
| FAM20C | 0.27255986 | 1.22E-36 | 1.73E-32 QQ |
| MDM2 | 0.27232312 | 3.09E-16 | 4.38E-12 QQ |
| ABCA1 | 0.27139372 | 1.49E-79 | 2.10E-75 QQ |
| TP53TG1 | 0.27075483 | 9.61E-40 | 1.36E-35 QQ |
| TNFRSF10B | 0.27060561 | 3.49E-39 | 4.94E-35 QQ |
| SMPDL3A | 0.27057532 | 2.41E-36 | 3.41E-32 QQ |
| CEMIP | 0.27042739 | 3.44E-22 | 4.86E-18 QQ |
| METRN | 0.27038822 | 2.90E-21 | 4.10E-17 QQ |
| B2M | 0.26966089 | 1.64E-176 | 2.33E-172 QQ |
| HAGH | 0.26940211 | 3.70E-33 | 5.23E-29 QQ |
| RNF7 | 0.26872226 | 1.12E-125 | 1.59E-121 QQ |
| EIF3G | 0.26867448 | 2.66E-163 | 3.77E-159 QQ |
| TEX264 | 0.26796668 | 9.54E-70 | 1.35E-65 QQ |
| TMEM106B | 0.26786213 | 2.36E-55 | 3.34E-51 QQ |
| RAB29 | 0.26771026 | 1.11E-30 | 1.56E-26 QQ |
| VPS41 | 0.26767571 | 8.24E-34 | 1.17E-29 QQ |
| IFT43 | 0.26710088 | 1.05E-50 | 1.48E-46 QQ |
| SDSL | 0.26683862 | 7.22E-26 | 1.02E-21 QQ |
| DDX3X | 0.26666565 | 4.02E-40 | 5.69E-36 QQ |
| CD27-AS1 | 0.26663961 | 2.45E-48 | 3.47E-44 QQ |
| PHPT1 | 0.26655486 | 5.12E-63 | 7.24E-59 QQ |
| ECHDC2 | 0.2664724 | 5.19E-44 | 7.35E-40 QQ |
| RPAIN | 0.26647012 | 4.96E-77 | 7.01E-73 QQ |
| ATRAID | 0.26637071 | 6.72E-89 | 9.50E-85 QQ |
| SIL1 | 0.26614052 | 2.24E-45 | 3.18E-41 QQ |
| TPGS1 | 0.26612183 | 1.82E-72 | 2.58E-68 QQ |

|  |  |  |  |
| --- | --- | --- | --- |
| C5orf38 | 0.26606351 | 1.01E-57 | 1.42E-53 QQ |
| XBP1 | 0.26601976 | 4.78E-15 | 6.76E-11 QQ |
| CCDC106 | 0.26595855 | 2.22E-45 | 3.14E-41 QQ |
| TXLNB | 0.26564199 | 8.12E-56 | 1.15E-51 QQ |
| WLS | 0.26557073 | 7.96E-43 | 1.13E-38 QQ |
| NCS1 | 0.26530811 | 2.43E-45 | 3.44E-41 QQ |
| ANGPTL2 | 0.26504092 | 1.89E-15 | 2.67E-11 QQ |
| CRTAP | 0.2644151 | 6.60E-69 | 9.34E-65 QQ |
| KDELR1 | 0.26409346 | 2.69E-105 | 3.81E-101 QQ |
| GRINA | 0.26400509 | 1.45E-63 | 2.05E-59 QQ |
| PLTP | 0.26389255 | 9.86E-31 | 1.39E-26 QQ |
| MLF1 | 0.2638004 | 6.61E-37 | 9.35E-33 QQ |
| MYLK | 0.26368817 | 1.08E-17 | 1.52E-13 QQ |
| ISG20 | 0.2636806 | 6.63E-14 | 9.39E-10 QQ |
| CCDC107 | 0.26360975 | 4.28E-53 | 6.05E-49 QQ |
| SSBP2 | 0.26355106 | 3.46E-49 | 4.90E-45 QQ |
| NFIL3 | 0.26346864 | 2.11E-20 | 2.98E-16 QQ |
| RILP | 0.26341169 | 1.24E-42 | 1.75E-38 QQ |
| COX4I1 | 0.26331175 | 9.31E-183 | 1.32E-178 QQ |
| MOSPD1 | 0.26330006 | 1.27E-25 | 1.79E-21 QQ |
| STAT3 | 0.26284915 | 1.01E-18 | 1.43E-14 QQ |
| MT-ND1 | 0.26229428 | 3.39E-35 | 4.79E-31 QQ |
| UBL3 | 0.26222921 | 3.43E-18 | 4.85E-14 QQ |
| NMB | 0.26214168 | 1.82E-33 | 2.58E-29 QQ |
| DDX3Y | 0.26192029 | 2.12E-31 | 3.00E-27 QQ |
| DNAJC4 | 0.26174644 | 1.82E-56 | 2.57E-52 QQ |
| CRELD1 | 0.2615224 | 1.15E-46 | 1.63E-42 QQ |
| DDR1 | 0.26141525 | 1.07E-36 | 1.52E-32 QQ |
| TSPAN10 | 0.26132536 | 5.29E-31 | 7.48E-27 QQ |
| APRT | 0.26124125 | 1.01E-104 | 1.43E-100 QQ |
| ANAPC16 | 0.26116255 | 1.96E-64 | 2.77E-60 QQ |
| NME5 | 0.26113371 | 6.94E-40 | 9.82E-36 QQ |
| BAALC | 0.26065274 | 7.88E-81 | 1.11E-76 QQ |
| SIRT2 | 0.26061434 | 2.64E-41 | 3.73E-37 QQ |
| CNPY2 | 0.26059203 | 1.82E-126 | 2.58E-122 QQ |
| FIS1 | 0.26034658 | 5.73E-102 | 8.10E-98 QQ |
| AR | 0.26008993 | 5.90E-37 | 8.35E-33 QQ |
| NAGLU | 0.26002726 | 1.43E-42 | 2.02E-38 QQ |
| HERPUD1 | 0.25998321 | 9.27E-41 | 1.31E-36 QQ |
| TBXA2R | 0.25939453 | 1.60E-38 | 2.26E-34 QQ |
| PPARGC1A | 0.25936203 | 1.37E-33 | 1.95E-29 QQ |
| SLK | 0.2593262 | 4.28E-27 | 6.05E-23 QQ |
| RRAGB | 0.25922251 | 7.26E-38 | 1.03E-33 QQ |
| ZNF281 | 0.2591292 | 5.65E-25 | 8.00E-21 QQ |
| SLC3A2 | 0.25890904 | 4.98E-27 | 7.04E-23 QQ |
| BTG2 | 0.25873209 | 1.23E-37 | 1.74E-33 QQ |

|  |  |  |  |
| --- | --- | --- | --- |
| SHISA2 | 0.25857785 | 2.63E-21 | 3.72E-17 QQ |
| MMP24OS | 0.25830048 | 2.88E-66 | 4.08E-62 QQ |
| MME | 0.25806868 | 6.82E-29 | 9.64E-25 QQ |
| STXBP3 | 0.25805305 | 9.12E-42 | 1.29E-37 QQ |
| KCNS3 | 0.25766387 | 7.46E-55 | 1.05E-50 QQ |
| PPIC | 0.25750535 | 6.60E-42 | 9.34E-38 QQ |
| EIF3K | 0.25692999 | 1.93E-176 | 2.74E-172 QQ |
| SSR4 | 0.25656162 | 7.99E-162 | 1.13E-157 QQ |
| RPL5 | 0.25643863 | 0 | 0 QQ |
| CCNH | 0.25620504 | 1.07E-20 | 1.51E-16 QQ |
| FOSL2 | 0.25581226 | 2.98E-17 | 4.22E-13 QQ |
| SYT14 | 0.25509576 | 4.76E-60 | 6.74E-56 QQ |
| ZNF428 | 0.25504251 | 8.54E-76 | 1.21E-71 QQ |
| SSPN | 0.25498005 | 6.83E-30 | 9.66E-26 QQ |
| PPFIBP1 | 0.2548801 | 2.85E-32 | 4.03E-28 QQ |
| SERINC1 | 0.25476213 | 7.44E-34 | 1.05E-29 QQ |
| KCTD12 | 0.25446827 | 2.25E-56 | 3.18E-52 QQ |
| PFDN5 | 0.25440412 | 3.73E-231 | 5.28E-227 QQ |
| GADD45A | 0.25431416 | 9.23E-30 | 1.31E-25 QQ |
| VEGFB | 0.25418974 | 6.43E-42 | 9.10E-38 QQ |
| EIF4B | 0.25385342 | 7.59E-97 | 1.07E-92 QQ |
| NFE2L1 | 0.25351725 | 9.00E-23 | 1.27E-18 QQ |
| IFRD1 | 0.25349005 | 1.78E-27 | 2.52E-23 QQ |
| MIR22HG | 0.25312727 | 1.53E-19 | 2.16E-15 QQ |
| MAF | 0.25300317 | 3.90E-102 | 5.52E-98 QQ |
| TBC1D23 | 0.25280835 | 3.87E-41 | 5.48E-37 QQ |
| TCEAL1 | 0.25273531 | 3.64E-33 | 5.15E-29 QQ |
| RHOB | 0.25258117 | 3.42E-28 | 4.84E-24 QQ |
| DAG1 | 0.25211997 | 8.46E-20 | 1.20E-15 QQ |
| TMX4 | 0.25191102 | 1.61E-25 | 2.28E-21 QQ |
| HDAC5 | 0.25158448 | 1.20E-35 | 1.69E-31 QQ |
| SMIM14 | 0.2514834 | 7.19E-34 | 1.02E-29 QQ |
| RHOBTB3 | 0.25139249 | 4.63E-38 | 6.54E-34 QQ |
| SATB1 | 0.25127953 | 2.61E-41 | 3.69E-37 QQ |
| SKP1 | 0.25116926 | 7.82E-207 | 1.11E-202 QQ |
| TCEAL9 | 0.25116332 | 6.95E-123 | 9.83E-119 QQ |
| GALNT7 | 0.25113473 | 4.82E-23 | 6.82E-19 QQ |
| SLC16A4 | 0.25104992 | 7.98E-41 | 1.13E-36 QQ |
| REL | 0.250738 | 7.31E-29 | 1.03E-24 QQ |
| ZBED3 | 0.2506859 | 7.85E-34 | 1.11E-29 QQ |
| UBE2C | 1.90442571 | 0 | 0 QP |
| CDK1 | 1.81737978 | 0 | 0 QP |
| CCNB1 | 1.80323618 | 0 | 0 QP |
| HIST1H4C | 1.73988254 | 0 | 0 QP |
| CENPF | 1.73948351 | 0 | 0 QP |
| CDKN3 | 1.70799922 | 0 | 0 QP |

|  |  |  |  |
| --- | --- | --- | --- |
| TOP2A | 1.68534665 | 0 | 0 QP |
| PBK | 1.68069785 | 0 | 0 QP |
| MKI67 | 1.66629082 | 0 | 0 QP |
| NUSAP1 | 1.66246987 | 0 | 0 QP |
| HMGB2 | 1.6369199 | 0 | 0 QP |
| PRC1 | 1.60850076 | 0 | 0 QP |
| PTTG1 | 1.57474443 | 0 | 0 QP |
| ASPM | 1.52641837 | 0 | 0 QP |
| CDCA3 | 1.51351912 | 0 | 0 QP |
| UBE2S | 1.50557863 | 0 | 0 QP |
| CCNB2 | 1.48126917 | 0 | 0 QP |
| NUF2 | 1.4739749 | 0 | 0 QP |
| SMC4 | 1.47088407 | 0 | 0 QP |
| BIRC5 | 1.45883109 | 0 | 0 QP |
| CKS1B | 1.4271336 | 0 | 0 QP |
| ARL6IP1 | 1.41918619 | 0 | 0 QP |
| TPX2 | 1.39194098 | 0 | 0 QP |
| TUBB4B | 1.35633174 | 0 | 0 QP |
| PCLAF | 1.35372685 | 0 | 0 QP |
| DLGAP5 | 1.35249875 | 0 | 0 QP |
| CEP55 | 1.34166734 | 0 | 0 QP |
| TUBA1B | 1.33350961 | 0 | 0 QP |
| CDC20 | 1.3220711 | 0 | 0 QP |
| CENPW | 1.29756105 | 0 | 0 QP |
| TUBA1C | 1.2801727 | 0 | 0 QP |
| MAD2L1 | 1.25348414 | 0 | 0 QP |
| KIF20B | 1.23912549 | 0 | 0 QP |
| GTSE1 | 1.22956218 | 0 | 0 QP |
| KPNA2 | 1.22760691 | 0 | 0 QP |
| PLK1 | 1.22314296 | 0 | 0 QP |
| UBE2T | 1.17337398 | 0 | 0 QP |
| RRM2 | 1.16660194 | 0 | 0 QP |
| CENPA | 1.16480189 | 0 | 0 QP |
| H2AFZ | 1.16390915 | 0 | 0 QP |
| DEPDC1 | 1.16335285 | 0 | 0 QP |
| NEK2 | 1.15839715 | 0 | 0 QP |
| SPC25 | 1.15749617 | 0 | 0 QP |
| AURKB | 1.1492589 | 0 | 0 QP |
| KIF11 | 1.14161892 | 0 | 0 QP |
| NUCB2 | 1.14020178 | 0 | 0 QP |
| HMMR | 1.12841876 | 0 | 0 QP |
| TACC3 | 1.12431204 | 0 | 0 QP |
| TK1 | 1.11666551 | 0 | 0 QP |
| TYMS | 1.10670245 | 0 | 0 QP |
| CENPE | 1.10083385 | 0 | 0 QP |
| ANLN | 1.09795545 | 0 | 0 QP |

|  |  |  |  |
| --- | --- | --- | --- |
| HIST1H1B | 1.08338938 | 0 | 0 QP |
| CKS2 | 1.0828756 | 0 | 0 QP |
| ZWINT | 1.08103319 | 0 | 0 QP |
| H2AFX | 1.06194973 | 0 | 0 QP |
| HMGB3 | 1.06134615 | 0 | 0 QP |
| CCNA2 | 1.06022998 | 0 | 0 QP |
| TROAP | 1.05004002 | 0 | 0 QP |
| CDKN2D | 1.03802846 | 2.25E-298 | 3.18E-294 QP |
| SGO1 | 1.03122559 | 0 | 0 QP |
| FOXM1 | 1.02310948 | 0 | 0 QP |
| HIST1H3D | 1.02240057 | 1.22E-253 | 1.73E-249 QP |
| NCAPG | 1.01821798 | 0 | 0 QP |
| DTYMK | 1.0127408 | 0 | 0 QP |
| MXD3 | 1.00213846 | 0 | 0 QP |
| CENPM | 0.99820538 | 0 | 0 QP |
| KIF23 | 0.99565833 | 0 | 0 QP |
| CCNA1 | 0.97236116 | 0 | 0 QP |
| KIF2C | 0.96860588 | 0 | 0 QP |
| KIFC1 | 0.96353511 | 0 | 0 QP |
| DIAPH3 | 0.94924234 | 0 | 0 QP |
| TUBB | 0.94313076 | 0 | 0 QP |
| TMPO | 0.9395522 | 0 | 0 QP |
| SGO2 | 0.93930637 | 0 | 0 QP |
| RACGAP1 | 0.92876213 | 0 | 0 QP |
| AURKA | 0.9278277 | 0 | 0 QP |
| HJURP | 0.91648652 | 0 | 0 QP |
| CIP2A | 0.90954482 | 0 | 0 QP |
| PHF19 | 0.90202686 | 0 | 0 QP |
| NCAPD2 | 0.89382055 | 0 | 0 QP |
| STMN1 | 0.89041616 | 0 | 0 QP |
| HMGN2 | 0.88953498 | 0 | 0 QP |
| CENPK | 0.88430862 | 0 | 0 QP |
| DDX39A | 0.88002878 | 0 | 0 QP |
| JPT1 | 0.87904711 | 0 | 0 QP |
| POC1A | 0.87699917 | 0 | 0 QP |
| NMU | 0.86529858 | 0 | 0 QP |
| DHFR | 0.86465147 | 0 | 0 QP |
| KNSTRN | 0.86348311 | 0 | 0 QP |
| SMC2 | 0.86134865 | 0 | 0 QP |
| MZT1 | 0.85687943 | 0 | 0 QP |
| RPL39L | 0.85008738 | 0 | 0 QP |
| NDC80 | 0.84744866 | 0 | 0 QP |
| KNL1 | 0.84415916 | 0 | 0 QP |
| KIF4A | 0.84414311 | 0 | 0 QP |
| FOSL1 | 0.84152565 | 0 | 0 QP |
| ASF1B | 0.84013451 | 0 | 0 QP |

|  |  |  |  |
| --- | --- | --- | --- |
| GGH | 0.83976791 | 0 | 0 QP |
| HMGB1 | 0.83574287 | 0 | 0 QP |
| CENPU | 0.83469167 | 0 | 0 QP |
| CKAP2L | 0.8314841 | 0 | 0 QP |
| LMNB1 | 0.82995836 | 0 | 0 QP |
| RAD21 | 0.82430939 | 0 | 0 QP |
| CKAP2 | 0.81084133 | 1.98E-301 | 2.81E-297 QP |
| MIS18BP1 | 0.81064934 | 0 | 0 QP |
| UBALD2 | 0.80816236 | 2.38E-304 | 3.37E-300 QP |
| ECT2 | 0.80720514 | 0 | 0 QP |
| SHCBP1 | 0.80469439 | 0 | 0 QP |
| FEN1 | 0.79529138 | 0 | 0 QP |
| CIT | 0.7905839 | 0 | 0 QP |
| KIF14 | 0.78739142 | 0 | 0 QP |
| CDCA2 | 0.78250543 | 0 | 0 QP |
| LMNB2 | 0.77611767 | 0 | 0 QP |
| CENPN | 0.77426644 | 0 | 0 QP |
| PARBP | 0.77356083 | 0 | 0 QP |
| DEK | 0.77026693 | 0 | 0 QP |
| PIMREG | 0.76846068 | 0 | 0 QP |
| GMNN | 0.7673164 | 0 | 0 QP |
| BUB1 | 0.76650191 | 0 | 0 QP |
| BUB3 | 0.76638537 | 0 | 0 QP |
| ANP32E | 0.75925149 | 0 | 0 QP |
| HIST1H3B | 0.75064134 | 0 | 0 QP |
| ATAD2 | 0.73962186 | 0 | 0 QP |
| MND1 | 0.73928092 | 0 | 0 QP |
| H2AFV | 0.73478091 | 0 | 0 QP |
| C16orf95 | 0.72869244 | 0 | 0 QP |
| FBXO5 | 0.72516296 | 0 | 0 QP |
| TTK | 0.72049653 | 0 | 0 QP |
| ARHGAP11A | 0.71883934 | 0 | 0 QP |
| USP1 | 0.71811091 | 0 | 0 QP |
| C12orf75 | 0.71810129 | 0 | 0 QP |
| TMEM106C | 0.71629889 | 0 | 0 QP |
| DNAJC9 | 0.71496135 | 0 | 0 QP |
| CAVIN3 | 0.71366103 | 0 | 0 QP |
| RAD51AP1 | 0.71148316 | 0 | 0 QP |
| SPC24 | 0.7014185 | 0 | 0 QP |
| PSRC1 | 0.69781999 | 1.28E-304 | 1.81E-300 QP |
| CKAP5 | 0.69763083 | 3.82E-268 | 5.40E-264 QP |
| EZH2 | 0.69635455 | 0 | 0 QP |
| KIF20A | 0.69370915 | 0 | 0 QP |
| TUBB6 | 0.69339038 | 0 | 0 QP |
| HIST1H1C | 0.68801165 | 4.98E-94 | 7.04E-90 QP |
| ORC6 | 0.68756772 | 0 | 0 QP |

|  |  |  |  |
| --- | --- | --- | --- |
| ITGB3BP | 0.68600411 | 0 | 0 QP |
| LRR1 | 0.68005699 | 0 | 0 QP |
| RPA3 | 0.6784711 | 0 | 0 QP |
| NUCKS1 | 0.67616438 | 0 | 0 QP |
| CCDC34 | 0.6756341 | 0 | 0 QP |
| ESCO2 | 0.67481713 | 0 | 0 QP |
| CALM3 | 0.67134073 | 0 | 0 QP |
| DUT | 0.65470438 | 1.55E-209 | 2.20E-205 QP |
| HIST1H1E | 0.65339394 | 2.12E-137 | 3.00E-133 QP |
| CDCA5 | 0.65276898 | 0 | 0 QP |
| RRM1 | 0.64900924 | 1.49E-305 | 2.11E-301 QP |
| MCM7 | 0.64625805 | 3.91E-289 | 5.53E-285 QP |
| CCDC88A | 0.64607703 | 1.12E-286 | 1.58E-282 QP |
| ILF2 | 0.64408511 | 0 | 0 QP |
| HIST1H1A | 0.63556598 | 3.51E-206 | 4.97E-202 QP |
| NUDT1 | 0.63299471 | 0 | 0 QP |
| SKA2 | 0.63282459 | 0 | 0 QP |
| CLSPN | 0.63004909 | 0 | 0 QP |
| RHEB | 0.62999782 | 0 | 0 QP |
| SKA3 | 0.62983841 | 0 | 0 QP |
| C9orf40 | 0.6291924 | 0 | 0 QP |
| CBR3 | 0.62119972 | 0 | 0 QP |
| NCAPH | 0.61948256 | 0 | 0 QP |
| EXOSC8 | 0.61806111 | 0 | 0 QP |
| MNS1 | 0.617419 | 1.83E-301 | 2.58E-297 QP |
| SMS | 0.61678617 | 0 | 0 QP |
| NSD2 | 0.60941293 | 0 | 0 QP |
| NCEH1 | 0.60826886 | 3.35E-237 | 4.74E-233 QP |
| MELK | 0.6068153 | 0 | 0 QP |
| DLEU2 | 0.60468476 | 0 | 0 QP |
| PCNA | 0.60421879 | 2.64E-141 | 3.74E-137 QP |
| FANCI | 0.60379385 | 0 | 0 QP |
| HNRNPA2B1 | 0.60203087 | 0 | 0 QP |
| DBF4 | 0.60131354 | 1.08E-297 | 1.53E-293 QP |
| TCF19 | 0.60026798 | 0 | 0 QP |
| TEX30 | 0.59980138 | 0 | 0 QP |
| GAS2L3 | 0.59597354 | 0 | 0 QP |
| CLEC11A | 0.59521785 | 0 | 0 QP |
| CENPH | 0.59216106 | 0 | 0 QP |
| SNRPG | 0.59136086 | 0 | 0 QP |
| DNMT1 | 0.58910772 | 0 | 0 QP |
| COMMD4 | 0.58751093 | 0 | 0 QP |
| PSIP1 | 0.58653079 | 0 | 0 QP |
| SPDL1 | 0.58627181 | 1.00E-258 | 1.42E-254 QP |
| MYBL2 | 0.58565199 | 0 | 0 QP |
| FAM111A | 0.5825416 | 4.55E-243 | 6.43E-239 QP |

|  |  |  |  |
| --- | --- | --- | --- |
| CLIC1 | 0.58202486 | 0 | 0 QP |
| PRR11 | 0.57890438 | 0 | 0 QP |
| SAE1 | 0.57744204 | 0 | 0 QP |
| NUDT5 | 0.57146051 | 0 | 0 QP |
| C3orf14 | 0.57017611 | 0 | 0 QP |
| TMSB15A | 0.56971686 | 0 | 0 QP |
| YWHAH | 0.56597459 | 0 | 0 QP |
| KIF22 | 0.56593032 | 1.06E-277 | 1.49E-273 QP |
| TEDC1 | 0.56569839 | 0 | 0 QP |
| SNRPD1 | 0.56362232 | 0 | 0 QP |
| HES6 | 0.56138232 | 6.07E-79 | 8.60E-75 QP |
| TUBG1 | 0.56044834 | 0 | 0 QP |
| CDCA8 | 0.55991693 | 0 | 0 QP |
| VRK1 | 0.55920689 | 0 | 0 QP |
| TRIP13 | 0.55643159 | 0 | 0 QP |
| RANBP1 | 0.55546914 | 0 | 0 QP |
| RTKN2 | 0.5542388 | 0 | 0 QP |
| SPAG5 | 0.55232444 | 0 | 0 QP |
| MOK | 0.55195877 | 1.92E-142 | 2.72E-138 QP |
| CDKN2C | 0.55065515 | 1.22E-211 | 1.72E-207 QP |
| ACTG2 | 0.54760162 | 7.23E-228 | 1.02E-223 QP |
| NASP | 0.54720253 | 0 | 0 QP |
| PLGRKT | 0.54707159 | 0 | 0 QP |
| CDC45 | 0.54544881 | 2.99E-231 | 4.23E-227 QP |
| PTMS | 0.54452601 | 1.35E-267 | 1.91E-263 QP |
| DEPDC1B | 0.54115307 | 0 | 0 QP |
| HMGN1 | 0.5411506 | 0 | 0 QP |
| REEP4 | 0.54077019 | 3.58E-223 | 5.07E-219 QP |
| AKR1C3 | 0.53920473 | 4.74E-192 | 6.70E-188 QP |
| LBR | 0.5372354 | 2.18E-239 | 3.08E-235 QP |
| RFC4 | 0.53634952 | 1.04E-221 | 1.47E-217 QP |
| LYAR | 0.53577158 | 4.61E-305 | 6.53E-301 QP |
| LSM5 | 0.53427641 | 0 | 0 QP |
| KIF15 | 0.53414043 | 0 | 0 QP |
| EMC9 | 0.53316194 | 1.32E-224 | 1.86E-220 QP |
| TPRKB | 0.52977882 | 0 | 0 QP |
| NUDT15 | 0.52928022 | 0 | 0 QP |
| HIST1H2AG | 0.52788187 | 1.11E-218 | 1.57E-214 QP |
| CDCA4 | 0.52715987 | 8.21E-239 | 1.16E-234 QP |
| FAM83D | 0.52252022 | 0 | 0 QP |
| SVIP | 0.52213693 | 1.78E-263 | 2.51E-259 QP |
| GIN5 | 0.5203338 | 4.20E-199 | 5.94E-195 QP |
| BCL2L12 | 0.52032221 | 2.35E-301 | 3.33E-297 QP |
| PKMYT1 | 0.51710072 | 0 | 0 QP |
| HNRNPAB | 0.51669442 | 0 | 0 QP |
| NUP37 | 0.51603023 | 7.31E-295 | 1.03E-290 QP |

|  |  |  |  |  |
| --- | --- | --- | --- | --- |
| CDT1 | 0.51514914 | 1.06E-266 | 1.50E-262 | QP |
| MASTL | 0.5142981 | 0 | 0 | QP |
| DRAP1 | 0.51260841 | 0 | 0 | QP |
| MIS18A | 0.51198203 | 4.83E-289 | 6.83E-285 | QP |
| CMC2 | 0.51102112 | 6.18E-275 | 8.74E-271 | QP |
| PBX3 | 0.51089391 | 1.74E-223 | 2.47E-219 | QP |
| BUB1B | 0.50601117 | 0 | 0 | QP |
| HNRNPH3 | 0.50593828 | 0 | 0 | QP |
| WDR34 | 0.50499223 | 4.67E-254 | 6.60E-250 | QP |
| CCDC18 | 0.50279598 | 3.06E-267 | 4.33E-263 | QP |
| RAN | 0.50203855 | 0 | 0 | QP |
| KIF18A | 0.49901451 | 1.32E-200 | 1.87E-196 | QP |
| PPP1R35 | 0.49824597 | 1.42E-209 | 2.01E-205 | QP |
| SEPHS1 | 0.49740993 | 2.51E-270 | 3.56E-266 | QP |
| SNRPB | 0.49697648 | 0 | 0 | QP |
| CNIH4 | 0.49577449 | 0 | 0 | QP |
| SIVA1 | 0.49371085 | 0 | 0 | QP |
| ENPP1 | 0.49281981 | 2.76E-157 | 3.91E-153 | QP |
| FAM111B | 0.49080007 | 8.74E-111 | 1.24E-106 | QP |
| CRNDE | 0.48943097 | 3.82E-237 | 5.40E-233 | QP |
| TPM4 | 0.48931168 | 0 | 0 | QP |
| UHRF1 | 0.48891912 | 2.90E-252 | 4.10E-248 | QP |
| CAPG | 0.48661428 | 1.95E-160 | 2.76E-156 | QP |
| HMG20B | 0.48640838 | 4.67E-186 | 6.60E-182 | QP |
| DAZAP1 | 0.48360159 | 1.08E-292 | 1.53E-288 | QP |
| ACTL6A | 0.48148864 | 0 | 0 | QP |
| MAGOHB | 0.48017264 | 8.74E-290 | 1.24E-285 | QP |
| VPS29 | 0.48000153 | 0 | 0 | QP |
| CDC25B | 0.47965079 | 1.99E-163 | 2.82E-159 | QP |
| C8orf88 | 0.47916504 | 3.33E-269 | 4.71E-265 | QP |
| GIHCG | 0.47893633 | 2.06E-213 | 2.91E-209 | QP |
| LPXN | 0.47811931 | 8.37E-228 | 1.18E-223 | QP |
| SRSF2 | 0.47741502 | 0 | 0 | QP |
| MAZ | 0.47642111 | 7.21E-261 | 1.02E-256 | QP |
| HP1BP3 | 0.47629144 | 1.58E-189 | 2.23E-185 | QP |
| BARD1 | 0.47573551 | 3.95E-210 | 5.59E-206 | QP |
| CAVIN2 | 0.47181017 | 4.89E-164 | 6.92E-160 | QP |
| PAK1 | 0.47001375 | 8.85E-246 | 1.25E-241 | QP |
| SFPQ | 0.46924464 | 1.33E-245 | 1.88E-241 | QP |
| HMGN5 | 0.46821877 | 4.25E-255 | 6.02E-251 | QP |
| FUS | 0.46748017 | 0 | 0 | QP |
| CSE1L | 0.466402 | 1.01E-257 | 1.43E-253 | QP |
| ANP32B | 0.4649029 | 0 | 0 | QP |
| CBX3 | 0.46213715 | 0 | 0 | QP |
| SMTN | 0.46152836 | 6.50E-197 | 9.20E-193 | QP |
| CSRP2 | 0.45999552 | 6.61E-144 | 9.35E-140 | QP |

|  |  |  |  |  |
| --- | --- | --- | --- | --- |
| PGP | 0.45635169 | 7.37E-278 | 1.04E-273 | QP |
| SAC3D1 | 0.45606496 | 3.57E-211 | 5.05E-207 | QP |
| CCT5 | 0.45586425 | 0 | 0 | QP |
| ALYREF | 0.45519389 | 1.02E-251 | 1.44E-247 | QP |
| S100A16 | 0.45480469 | 1.93E-281 | 2.73E-277 | QP |
| HNRNPD | 0.4540639 | 0 | 0 | QP |
| PSMC3 | 0.45230533 | 0 | 0 | QP |
| FANCG | 0.45206656 | 1.22E-242 | 1.73E-238 | QP |
| NT5DC2 | 0.45106141 | 5.51E-257 | 7.80E-253 | QP |
| IKBIP | 0.44990594 | 1.52E-227 | 2.14E-223 | QP |
| CENPQ | 0.44831602 | 1.03E-210 | 1.46E-206 | QP |
| CBX1 | 0.44573868 | 4.47E-207 | 6.32E-203 | QP |
| HPRT1 | 0.44383063 | 2.47E-244 | 3.50E-240 | QP |
| HNRNPA3 | 0.44312211 | 0 | 0 | QP |
| HELLS | 0.44105359 | 2.69E-124 | 3.80E-120 | QP |
| KPNB1 | 0.43871055 | 2.95E-253 | 4.18E-249 | QP |
| SRSF7 | 0.43866717 | 4.80E-260 | 6.80E-256 | QP |
| HSPB11 | 0.43828736 | 4.80E-170 | 6.79E-166 | QP |
| PTGES3 | 0.43810774 | 0 | 0 | QP |
| SNRPA | 0.4380067 | 6.40E-249 | 9.06E-245 | QP |
| TOMM40 | 0.43701056 | 3.52E-264 | 4.98E-260 | QP |
| HNRNPR | 0.43422799 | 9.12E-283 | 1.29E-278 | QP |
| COX8A | 0.43291491 | 0 | 0 | QP |
| MARCKS | 0.43219401 | 3.04E-165 | 4.29E-161 | QP |
| DDX39B | 0.43217027 | 2.13E-260 | 3.01E-256 | QP |
| RHNO1 | 0.43152202 | 1.73E-218 | 2.45E-214 | QP |
| CDC25C | 0.43002333 | 0 | 0 | QP |
| BCL2A1 | 0.42849136 | 3.94E-110 | 5.57E-106 | QP |
| ARHGDI | 0.4283645 | 1.98E-176 | 2.80E-172 | QP |
| NUDCD2 | 0.42755918 | 4.27E-206 | 6.03E-202 | QP |
| C19orf48 | 0.42732982 | 1.39E-198 | 1.97E-194 | QP |
| PA2G4 | 0.42726236 | 0 | 0 | QP |
| LGALS1 | 0.42724554 | 1.08E-290 | 1.52E-286 | QP |
| MAGOH | 0.4252595 | 1.90E-307 | 2.69E-303 | QP |
| RUVBL2 | 0.42325815 | 7.45E-232 | 1.05E-227 | QP |
| HNRNPM | 0.42312059 | 0 | 0 | QP |
| RANGAP1 | 0.42026806 | 3.74E-182 | 5.30E-178 | QP |
| FAM102B | 0.42006489 | 1.31E-206 | 1.85E-202 | QP |
| CACYBP | 0.41745793 | 8.32E-261 | 1.18E-256 | QP |
| AXL | 0.41616359 | 6.30E-209 | 8.91E-205 | QP |
| MAD2L2 | 0.41606064 | 2.40E-208 | 3.40E-204 | QP |
| EIF5A | 0.41550127 | 2.91E-275 | 4.12E-271 | QP |
| UAP1 | 0.4153924 | 5.52E-170 | 7.81E-166 | QP |
| NGFR | 0.41518232 | 3.51E-178 | 4.97E-174 | QP |
| SNRPF | 0.4141896 | 9.79E-268 | 1.39E-263 | QP |
| CENPX | 0.41401557 | 6.57E-151 | 9.30E-147 | QP |

|  |  |  |  |  |
| --- | --- | --- | --- | --- |
| PLEKHJ1 | 0.41295845 | 1.54E-243 | 2.18E-239 | QP |
| NRM | 0.41257707 | 6.53E-179 | 9.24E-175 | QP |
| PLP2 | 0.4123646 | 1.01E-266 | 1.44E-262 | QP |
| HDGF | 0.41211113 | 6.00E-248 | 8.48E-244 | QP |
| SSRP1 | 0.41006519 | 1.72E-240 | 2.43E-236 | QP |
| RBMX | 0.40912907 | 2.35E-274 | 3.33E-270 | QP |
| E2F1 | 0.40780143 | 1.71E-108 | 2.41E-104 | QP |
| MCM4 | 0.40694652 | 1.50E-138 | 2.12E-134 | QP |
| H1FX | 0.40662761 | 9.79E-190 | 1.39E-185 | QP |
| KIF5B | 0.40569712 | 1.75E-127 | 2.48E-123 | QP |
| SRSF3 | 0.40330088 | 8.25E-305 | 1.17E-300 | QP |
| LSM4 | 0.40283578 | 0 | 0 | QP |
| MCM10 | 0.40257738 | 9.16E-220 | 1.30E-215 | QP |
| HAO1 | 0.40205837 | 4.17E-193 | 5.90E-189 | QP |
| ACTN4 | 0.40013836 | 1.15E-248 | 1.62E-244 | QP |
| LSM8 | 0.39989959 | 1.89E-276 | 2.67E-272 | QP |
| ACTB | 0.39924892 | 2.33E-282 | 3.30E-278 | QP |
| RFC3 | 0.39798224 | 3.60E-213 | 5.10E-209 | QP |
| XRCC6 | 0.39780884 | 1.44E-306 | 2.03E-302 | QP |
| ELOVL3 | 0.39734921 | 1.04E-31 | 1.46E-27 | QP |
| PRIM1 | 0.3968525 | 1.84E-144 | 2.60E-140 | QP |
| CKLF | 0.39560565 | 4.89E-192 | 6.93E-188 | QP |
| EEF1AKMT2 | 0.39506081 | 1.45E-160 | 2.06E-156 | QP |
| PAXX | 0.39478577 | 3.23E-240 | 4.57E-236 | QP |
| HAT1 | 0.39415122 | 8.64E-175 | 1.22E-170 | QP |
| TIMM10 | 0.39382644 | 1.54E-219 | 2.19E-215 | QP |
| CENPL | 0.39345228 | 4.09E-220 | 5.78E-216 | QP |
| RAD51 | 0.39332967 | 3.83E-258 | 5.41E-254 | QP |
| CHAF1A | 0.39267799 | 2.80E-171 | 3.96E-167 | QP |
| TNFRSF12A | 0.39208811 | 3.27E-196 | 4.63E-192 | QP |
| NOP58 | 0.39153377 | 1.13E-200 | 1.60E-196 | QP |
| DBI | 0.3908412 | 0 | 0 | QP |
| PPP1CA | 0.39020688 | 1.23E-290 | 1.74E-286 | QP |
| LSM3 | 0.38970933 | 0 | 0 | QP |
| MCM3 | 0.38894023 | 6.45E-88 | 9.12E-84 | QP |
| NCAPG2 | 0.38766206 | 2.72E-247 | 3.84E-243 | QP |
| NCAPH2 | 0.38746234 | 2.63E-183 | 3.72E-179 | QP |
| LSM6 | 0.38724338 | 7.52E-194 | 1.06E-189 | QP |
| GLRX5 | 0.38638435 | 3.97E-239 | 5.61E-235 | QP |
| IDH2 | 0.38578974 | 1.41E-89 | 2.00E-85 | QP |
| TFDP1 | 0.38545603 | 2.61E-177 | 3.69E-173 | QP |
| HIST2H2AC | 0.38457006 | 5.84E-41 | 8.26E-37 | QP |
| NT5E | 0.38375534 | 2.93E-172 | 4.15E-168 | QP |
| NUDT21 | 0.38352042 | 7.90E-196 | 1.12E-191 | QP |
| CHEK1 | 0.3824966 | 6.18E-211 | 8.75E-207 | QP |
| SYNE2 | 0.38047818 | 4.34E-114 | 6.15E-110 | QP |

|  |  |  |  |  |
| --- | --- | --- | --- | --- |
| PLAUR | 0.3801037 | 2.41E-113 | 3.41E-109 | QP |
| POLE3 | 0.37925979 | 5.07E-163 | 7.17E-159 | QP |
| EZR | 0.37848246 | 8.96E-154 | 1.27E-149 | QP |
| LRRC59 | 0.37810478 | 5.13E-198 | 7.25E-194 | QP |
| SAPCD2 | 0.37654055 | 7.31E-267 | 1.03E-262 | QP |
| CDC6 | 0.37477911 | 2.05E-155 | 2.89E-151 | QP |
| CBX5 | 0.37439461 | 4.45E-201 | 6.30E-197 | QP |
| TAF9 | 0.37403968 | 4.66E-235 | 6.60E-231 | QP |
| ILK | 0.37282518 | 3.80E-173 | 5.38E-169 | QP |
| TMEM19 | 0.37274088 | 7.31E-164 | 1.03E-159 | QP |
| MRPL16 | 0.37231481 | 1.82E-171 | 2.58E-167 | QP |
| SSNA1 | 0.3719851 | 4.42E-247 | 6.25E-243 | QP |
| CASP3 | 0.37195787 | 4.61E-97 | 6.52E-93 | QP |
| E2F8 | 0.37181179 | 9.99E-188 | 1.41E-183 | QP |
| SRP9 | 0.37085107 | 0 | 0 | QP |
| MAP1B | 0.36930216 | 4.30E-143 | 6.09E-139 | QP |
| OIP5 | 0.36894281 | 1.17E-299 | 1.65E-295 | QP |
| PXMP2 | 0.36694575 | 1.39E-132 | 1.96E-128 | QP |
| BOLA3 | 0.36628998 | 7.45E-183 | 1.05E-178 | QP |
| ZNF511 | 0.36547196 | 1.53E-227 | 2.16E-223 | QP |
| IRAK1 | 0.36298823 | 4.98E-153 | 7.04E-149 | QP |
| USP13 | 0.36284737 | 2.22E-130 | 3.14E-126 | QP |
| PFN1 | 0.36211777 | 0 | 0 | QP |
| SNRPA1 | 0.36193107 | 4.51E-194 | 6.39E-190 | QP |
| ZWILCH | 0.36170611 | 4.76E-198 | 6.74E-194 | QP |
| BCL7C | 0.36159439 | 1.46E-211 | 2.07E-207 | QP |
| CARHSP1 | 0.36136151 | 8.99E-191 | 1.27E-186 | QP |
| HNRNPUL2 | 0.35951292 | 7.23E-146 | 1.02E-141 | QP |
| GPSM2 | 0.35942514 | 1.01E-101 | 1.43E-97 | QP |
| PHGDH | 0.35904477 | 2.41E-160 | 3.41E-156 | QP |
| SYNCRIP | 0.35834188 | 7.44E-171 | 1.05E-166 | QP |
| BRCA2 | 0.35685439 | 6.60E-206 | 9.34E-202 | QP |
| YWHAE | 0.35684755 | 3.06E-191 | 4.33E-187 | QP |
| SNRPE | 0.35676621 | 4.22E-243 | 5.97E-239 | QP |
| ARPC5L | 0.35675346 | 2.90E-200 | 4.10E-196 | QP |
| CNTRL | 0.35592768 | 8.61E-159 | 1.22E-154 | QP |
| RFC5 | 0.35579861 | 2.40E-177 | 3.39E-173 | QP |
| BTG3 | 0.3556862 | 3.08E-146 | 4.35E-142 | QP |
| EHD1 | 0.35516828 | 4.90E-146 | 6.93E-142 | QP |
| RBM8A | 0.35456273 | 1.09E-304 | 1.54E-300 | QP |
| CDK2 | 0.35353494 | 3.15E-185 | 4.46E-181 | QP |
| VASP | 0.3530558 | 8.06E-130 | 1.14E-125 | QP |
| TCP1 | 0.35138917 | 4.67E-171 | 6.61E-167 | QP |
| SNX7 | 0.35114743 | 3.09E-178 | 4.37E-174 | QP |
| SMC3 | 0.35039428 | 4.72E-137 | 6.67E-133 | QP |
| FBL | 0.35023662 | 2.73E-219 | 3.87E-215 | QP |

|  |  |  |  |
| --- | --- | --- | --- |
| ERH | 0.34970104 | 0 | 0 QP |
| BRCA1 | 0.34907752 | 2.13E-162 | 3.02E-158 QP |
| PPP1R14B | 0.34847741 | 7.66E-282 | 1.08E-277 QP |
| BLM | 0.34801374 | 2.62E-209 | 3.71E-205 QP |
| SUMO3 | 0.34770014 | 2.20E-209 | 3.11E-205 QP |
| DTL | 0.34742722 | 1.06E-146 | 1.50E-142 QP |
| STIP1 | 0.34687987 | 4.85E-182 | 6.87E-178 QP |
| ENY2 | 0.34662445 | 1.94E-307 | 2.74E-303 QP |
| CD82 | 0.34633616 | 5.36E-98 | 7.59E-94 QP |
| CEP78 | 0.34631791 | 6.99E-156 | 9.89E-152 QP |
| CXCL3 | 0.34573833 | 6.46E-56 | 9.13E-52 QP |
| TM4SF1 | 0.34540116 | 5.90E-62 | 8.35E-58 QP |
| HSP90B1 | 0.34492974 | 2.25E-161 | 3.19E-157 QP |
| SRPK1 | 0.34428055 | 6.37E-160 | 9.02E-156 QP |
| KHDRBS1 | 0.34418802 | 1.50E-198 | 2.13E-194 QP |
| HIST1H2AH | 0.34367583 | 4.55E-156 | 6.44E-152 QP |
| SLBP | 0.34291356 | 3.04E-81 | 4.30E-77 QP |
| MED30 | 0.34264781 | 1.47E-146 | 2.09E-142 QP |
| BANF1 | 0.34221989 | 1.02E-278 | 1.44E-274 QP |
| SKA1 | 0.34211815 | 3.30E-299 | 4.66E-295 QP |
| ID1 | 0.34195835 | 1.48E-58 | 2.09E-54 QP |
| UBE2N | 0.3418389 | 2.58E-198 | 3.66E-194 QP |
| TPM3 | 0.34159844 | 6.92E-188 | 9.79E-184 QP |
| SRRT | 0.34095563 | 5.36E-144 | 7.59E-140 QP |
| HNRNPUL1 | 0.34029461 | 2.99E-164 | 4.23E-160 QP |
| ING2 | 0.34008541 | 4.39E-152 | 6.22E-148 QP |
| HIRIP3 | 0.33947567 | 9.70E-133 | 1.37E-128 QP |
| PTBP1 | 0.33904835 | 2.10E-157 | 2.97E-153 QP |
| RNASEH2B | 0.33857457 | 4.04E-143 | 5.72E-139 QP |
| GUSB | 0.33749066 | 8.23E-149 | 1.16E-144 QP |
| HINT1 | 0.33733894 | 0 | 0 QP |
| EFHD2 | 0.33719342 | 1.32E-147 | 1.87E-143 QP |
| HYLS1 | 0.33566315 | 2.79E-155 | 3.95E-151 QP |
| EXO1 | 0.33561336 | 2.58E-199 | 3.64E-195 QP |
| WDR62 | 0.33469285 | 4.28E-267 | 6.05E-263 QP |
| HAUS8 | 0.33456769 | 2.33E-199 | 3.30E-195 QP |
| RALY | 0.33349806 | 4.01E-159 | 5.67E-155 QP |
| COQ2 | 0.33263108 | 1.65E-174 | 2.34E-170 QP |
| PSMD9 | 0.3323612 | 1.38E-154 | 1.95E-150 QP |
| SEPTIN10 | 0.33174143 | 5.54E-136 | 7.85E-132 QP |
| PPIH | 0.3312539 | 4.42E-156 | 6.25E-152 QP |
| TRIM59 | 0.33025654 | 1.23E-114 | 1.74E-110 QP |
| MYOD1 | 0.32940653 | 7.99E-39 | 1.13E-34 QP |
| PSMD14 | 0.3292954 | 4.39E-171 | 6.21E-167 QP |
| ARPC2 | 0.32924757 | 5.53E-261 | 7.82E-257 QP |
| DMBT1 | 0.32874761 | 4.63E-122 | 6.55E-118 QP |

|  |  |  |  |  |
| --- | --- | --- | --- | --- |
| MT2A | 0.32842074 | 3.93E-136 | 5.56E-132 | QP |
| TRIM28 | 0.32836093 | 5.25E-185 | 7.42E-181 | QP |
| ITGB1BP1 | 0.32802152 | 3.18E-188 | 4.50E-184 | QP |
| RFC2 | 0.32799693 | 1.61E-89 | 2.27E-85 | QP |
| PSMC3IP | 0.32735087 | 2.31E-150 | 3.26E-146 | QP |
| SCLT1 | 0.32651096 | 5.45E-141 | 7.72E-137 | QP |
| MCM5 | 0.32646746 | 3.45E-76 | 4.88E-72 | QP |
| HDAC2 | 0.32616161 | 1.89E-136 | 2.67E-132 | QP |
| FANCD2 | 0.32542485 | 3.15E-231 | 4.46E-227 | QP |
| PAK4 | 0.32483991 | 3.90E-126 | 5.52E-122 | QP |
| PSMD2 | 0.32427336 | 3.91E-158 | 5.53E-154 | QP |
| ATAD5 | 0.32364145 | 4.35E-157 | 6.16E-153 | QP |
| FGF5 | 0.32362131 | 2.41E-97 | 3.42E-93 | QP |
| CCNE2 | 0.32328383 | 4.24E-110 | 6.00E-106 | QP |
| SUPT16H | 0.32325171 | 8.62E-104 | 1.22E-99 | QP |
| ACYP1 | 0.32295985 | 4.66E-120 | 6.59E-116 | QP |
| MT1E | 0.3226611 | 4.31E-113 | 6.10E-109 | QP |
| FH | 0.32241448 | 3.34E-146 | 4.73E-142 | QP |
| HIST1H2AL | 0.32232829 | 4.35E-177 | 6.16E-173 | QP |
| LIG1 | 0.32227084 | 2.79E-101 | 3.95E-97 | QP |
| PIH1D1 | 0.32176736 | 4.78E-181 | 6.76E-177 | QP |
| PAFAH1B3 | 0.32166371 | 6.05E-111 | 8.57E-107 | QP |
| PPIF | 0.32164371 | 3.04E-121 | 4.30E-117 | QP |
| DKC1 | 0.32152795 | 6.15E-137 | 8.70E-133 | QP |
| PPM1G | 0.32150629 | 2.88E-172 | 4.07E-168 | QP |
| ERCC6L | 0.3214759 | 1.65E-245 | 2.33E-241 | QP |
| TMEM237 | 0.32115739 | 2.93E-134 | 4.14E-130 | QP |
| ECM1 | 0.32066068 | 2.09E-128 | 2.96E-124 | QP |
| HNRNPL | 0.31982633 | 5.36E-142 | 7.59E-138 | QP |
| SAP30 | 0.31944393 | 1.21E-101 | 1.71E-97 | QP |
| VBP1 | 0.31931987 | 4.38E-173 | 6.20E-169 | QP |
| SLC25A5 | 0.31861529 | 1.45E-277 | 2.05E-273 | QP |
| PDAP1 | 0.31793958 | 2.63E-189 | 3.73E-185 | QP |
| FUBP1 | 0.31776003 | 1.06E-110 | 1.50E-106 | QP |
| ENO1 | 0.31699493 | 4.58E-205 | 6.49E-201 | QP |
| FDPS | 0.31594262 | 3.69E-164 | 5.22E-160 | QP |
| TMPO-AS1 | 0.31589592 | 6.41E-241 | 9.07E-237 | QP |
| INCENP | 0.31532802 | 1.00E-183 | 1.42E-179 | QP |
| STIL | 0.3147906 | 7.34E-222 | 1.04E-217 | QP |
| KMT5A | 0.31445347 | 8.42E-99 | 1.19E-94 | QP |
| CENPO | 0.31355663 | 3.43E-219 | 4.85E-215 | QP |
| AP1M1 | 0.31297512 | 2.42E-126 | 3.43E-122 | QP |
| ARPC1A | 0.31265787 | 2.12E-188 | 3.00E-184 | QP |
| WDR1 | 0.31253716 | 1.33E-145 | 1.88E-141 | QP |
| IQGAP3 | 0.31243136 | 8.71E-254 | 1.23E-249 | QP |
| PARP2 | 0.31207895 | 1.97E-124 | 2.79E-120 | QP |

|  |  |  |  |  |
| --- | --- | --- | --- | --- |
| TOPBP1 | 0.311687 | 4.92E-118 | 6.96E-114 | QP |
| MAPKAP1 | 0.31158894 | 3.15E-121 | 4.46E-117 | QP |
| H3F3B | 0.30915124 | 6.52E-172 | 9.23E-168 | QP |
| RAB8A | 0.30905813 | 9.43E-131 | 1.33E-126 | QP |
| HNRNPF | 0.3090394 | 5.45E-156 | 7.71E-152 | QP |
| FLOT1 | 0.30886875 | 1.69E-130 | 2.39E-126 | QP |
| SNX5 | 0.30798049 | 2.08E-122 | 2.94E-118 | QP |
| EHD4 | 0.30791956 | 1.65E-110 | 2.33E-106 | QP |
| EMB | 0.30782135 | 3.72E-124 | 5.27E-120 | QP |
| UQCC3 | 0.30725713 | 6.42E-139 | 9.08E-135 | QP |
| S100A2 | 0.30703855 | 2.25E-151 | 3.19E-147 | QP |
| TMEM173 | 0.30672039 | 4.75E-115 | 6.72E-111 | QP |
| CFL1 | 0.3064043 | 3.27E-264 | 4.63E-260 | QP |
| PRRG4 | 0.3062994 | 3.64E-138 | 5.16E-134 | QP |
| TPGS2 | 0.30576463 | 3.84E-122 | 5.43E-118 | QP |
| MBD3 | 0.3056922 | 4.24E-118 | 6.00E-114 | QP |
| ODF2 | 0.30567279 | 3.70E-144 | 5.23E-140 | QP |
| HSP90AA1 | 0.30521537 | 9.09E-152 | 1.29E-147 | QP |
| EMD | 0.30445293 | 1.99E-145 | 2.82E-141 | QP |
| KIF18B | 0.30440608 | 1.21E-247 | 1.71E-243 | QP |
| UQCC2 | 0.30430253 | 7.45E-133 | 1.05E-128 | QP |
| ZNF22 | 0.30406733 | 8.66E-120 | 1.23E-115 | QP |
| ARPC5 | 0.30395759 | 1.07E-178 | 1.52E-174 | QP |
| FAM122B | 0.30393765 | 1.38E-140 | 1.95E-136 | QP |
| HELLPAR | 0.30332742 | 3.79E-162 | 5.36E-158 | QP |
| NRGN | 0.30296687 | 7.06E-181 | 9.99E-177 | QP |
| HADH | 0.30287792 | 5.44E-104 | 7.70E-100 | QP |
| POP7 | 0.30239033 | 2.56E-141 | 3.63E-137 | QP |
| TPI1 | 0.30187559 | 2.77E-228 | 3.93E-224 | QP |
| TALDO1 | 0.30112765 | 1.68E-174 | 2.37E-170 | QP |
| COTL1 | 0.30093969 | 3.08E-108 | 4.36E-104 | QP |
| RAC2 | 0.30087632 | 7.20E-89 | 1.02E-84 | QP |
| PAQR4 | 0.29984741 | 9.92E-103 | 1.40E-98 | QP |
| MANCR | 0.299628 | 7.86E-90 | 1.11E-85 | QP |
| ARF6 | 0.29959279 | 1.79E-160 | 2.53E-156 | QP |
| CMSS1 | 0.29912703 | 4.59E-129 | 6.50E-125 | QP |
| NOP56 | 0.29901286 | 2.53E-123 | 3.58E-119 | QP |
| UBR7 | 0.29873779 | 1.26E-105 | 1.79E-101 | QP |
| QSER1 | 0.29787099 | 1.34E-93 | 1.89E-89 | QP |
| CENPS | 0.29763827 | 1.43E-129 | 2.03E-125 | QP |
| PCMT1 | 0.29763534 | 2.41E-144 | 3.41E-140 | QP |
| MZT2B | 0.29756613 | 8.52E-251 | 1.21E-246 | QP |
| EIF2S1 | 0.29623863 | 2.28E-139 | 3.23E-135 | QP |
| ALCAM | 0.29623313 | 2.41E-111 | 3.41E-107 | QP |
| MPP6 | 0.29599626 | 5.18E-108 | 7.33E-104 | QP |
| SSR3 | 0.29569505 | 4.13E-120 | 5.85E-116 | QP |

|  |  |  |  |  |
| --- | --- | --- | --- | --- |
| CDC123 | 0.29547931 | 4.02E-141 | 5.69E-137 | QP |
| PNP | 0.29495263 | 2.70E-86 | 3.82E-82 | QP |
| C19orf33 | 0.29447227 | 1.86E-96 | 2.63E-92 | QP |
| CDK6 | 0.293032 | 3.58E-86 | 5.07E-82 | QP |
| RPP30 | 0.29270711 | 2.04E-119 | 2.88E-115 | QP |
| ZNF367 | 0.29255405 | 6.18E-140 | 8.75E-136 | QP |
| DSN1 | 0.29242829 | 9.45E-114 | 1.34E-109 | QP |
| NUP107 | 0.29239239 | 2.53E-110 | 3.58E-106 | QP |
| TOMM5 | 0.29232874 | 2.92E-204 | 4.13E-200 | QP |
| ILF3 | 0.29210646 | 6.68E-129 | 9.45E-125 | QP |
| ANAPC11 | 0.29191555 | 9.50E-239 | 1.34E-234 | QP |
| YES1 | 0.29133607 | 1.17E-77 | 1.65E-73 | QP |
| RASA1 | 0.29125814 | 1.04E-101 | 1.47E-97 | QP |
| PRKDC | 0.29093867 | 8.70E-99 | 1.23E-94 | QP |
| HMGN3 | 0.2907417 | 2.48E-153 | 3.50E-149 | QP |
| APOBEC3B | 0.29031814 | 8.16E-233 | 1.15E-228 | QP |
| HNRNPK | 0.28967617 | 4.42E-219 | 6.26E-215 | QP |
| RHOA | 0.28930507 | 1.16E-258 | 1.64E-254 | QP |
| KIAA0586 | 0.28926418 | 2.28E-98 | 3.22E-94 | QP |
| FANCB | 0.28906222 | 1.30E-162 | 1.83E-158 | QP |
| EBNA1BP2 | 0.28899885 | 4.76E-118 | 6.74E-114 | QP |
| CDC25A | 0.28875466 | 1.41E-183 | 1.99E-179 | QP |
| STEAP1 | 0.28841316 | 3.35E-85 | 4.74E-81 | QP |
| DKK1 | 0.28832345 | 1.91E-14 | 2.70E-10 | QP |
| NAP1L4 | 0.28769705 | 1.17E-135 | 1.66E-131 | QP |
| PGAM1 | 0.28767575 | 1.06E-155 | 1.50E-151 | QP |
| SMC1A | 0.28691189 | 2.89E-89 | 4.09E-85 | QP |
| EIF4A3 | 0.28690362 | 1.26E-138 | 1.78E-134 | QP |
| MCUB | 0.28671793 | 7.81E-110 | 1.10E-105 | QP |
| MTHFD2 | 0.28661384 | 2.20E-128 | 3.11E-124 | QP |
| ACAT2 | 0.28603055 | 1.15E-75 | 1.63E-71 | QP |
| C1QTNF2 | 0.28554981 | 4.22E-100 | 5.97E-96 | QP |
| GIN5 | 0.28535853 | 5.94E-145 | 8.40E-141 | QP |
| VANGL1 | 0.28530021 | 2.08E-77 | 2.95E-73 | QP |
| NECTIN2 | 0.28519943 | 9.25E-79 | 1.31E-74 | QP |
| GTF2A2 | 0.2850456 | 1.40E-145 | 1.98E-141 | QP |
| NANS | 0.28500564 | 1.62E-112 | 2.29E-108 | QP |
| PSMA4 | 0.28481966 | 1.33E-215 | 1.88E-211 | QP |
| PRKRA | 0.28478013 | 7.71E-90 | 1.09E-85 | QP |
| PSMA7 | 0.2839489 | 1.66E-254 | 2.34E-250 | QP |
| RBM14 | 0.28340823 | 2.85E-112 | 4.03E-108 | QP |
| LSM2 | 0.28323261 | 1.09E-137 | 1.54E-133 | QP |
| MRPL11 | 0.28321267 | 5.73E-159 | 8.10E-155 | QP |
| ACTG1 | 0.28304655 | 1.54E-128 | 2.17E-124 | QP |
| MCMBP | 0.28236304 | 3.30E-91 | 4.67E-87 | QP |
| NRIP3 | 0.28234314 | 1.03E-98 | 1.45E-94 | QP |

|  |  |  |  |  |
| --- | --- | --- | --- | --- |
| EIF4EBP1 | 0.28181056 | 6.68E-104 | 9.45E-100 | QP |
| GANAB | 0.28153896 | 5.44E-113 | 7.69E-109 | QP |
| HSPA2 | 0.28105399 | 1.14E-100 | 1.62E-96 | QP |
| RECQL | 0.28101445 | 6.54E-113 | 9.25E-109 | QP |
| NFYB | 0.28096391 | 1.32E-93 | 1.86E-89 | QP |
| mar-03 | 0.28026346 | 6.79E-103 | 9.61E-99 | QP |
| POLA2 | 0.28001944 | 1.50E-115 | 2.12E-111 | QP |
| LRRCC1 | 0.27991093 | 9.12E-102 | 1.29E-97 | QP |
| CHAC2 | 0.27920734 | 5.15E-120 | 7.29E-116 | QP |
| ADK | 0.2790561 | 1.14E-112 | 1.62E-108 | QP |
| HSPA14 | 0.27865731 | 7.09E-106 | 1.00E-101 | QP |
| TRA2B | 0.27844462 | 2.74E-103 | 3.87E-99 | QP |
| SUN2 | 0.27821233 | 1.60E-103 | 2.26E-99 | QP |
| MYEOV | 0.27785952 | 8.26E-101 | 1.17E-96 | QP |
| C11orf24 | 0.27727465 | 1.34E-91 | 1.89E-87 | QP |
| MYBL1 | 0.27705914 | 9.08E-84 | 1.28E-79 | QP |
| CDK2AP2 | 0.27668386 | 5.49E-94 | 7.77E-90 | QP |
| LYPLA1 | 0.27562946 | 5.36E-103 | 7.58E-99 | QP |
| UBE2I | 0.27553832 | 5.28E-185 | 7.47E-181 | QP |
| TRAIP | 0.27534434 | 1.31E-193 | 1.85E-189 | QP |
| PSME2 | 0.27501806 | 9.68E-110 | 1.37E-105 | QP |
| SNRNP40 | 0.27479111 | 3.33E-105 | 4.71E-101 | QP |
| HNRNPA0 | 0.27460103 | 8.10E-149 | 1.15E-144 | QP |
| CDCA7 | 0.27435547 | 7.77E-93 | 1.10E-88 | QP |
| COPS3 | 0.27413957 | 4.20E-104 | 5.95E-100 | QP |
| POLD3 | 0.27359268 | 2.43E-73 | 3.43E-69 | QP |
| ASF1A | 0.27338297 | 5.29E-88 | 7.48E-84 | QP |
| MBD2 | 0.27245477 | 5.75E-114 | 8.14E-110 | QP |
| C1QL1 | 0.27179901 | 1.21E-87 | 1.71E-83 | QP |
| EBP | 0.27157727 | 4.68E-103 | 6.63E-99 | QP |
| ARHGDI | 0.27139263 | 1.15E-129 | 1.63E-125 | QP |
| LDHA | 0.27097909 | 2.36E-155 | 3.33E-151 | QP |
| PHF5A | 0.27084009 | 5.69E-112 | 8.06E-108 | QP |
| ECI2 | 0.2708352 | 9.18E-121 | 1.30E-116 | QP |
| ADRM1 | 0.27050888 | 8.32E-132 | 1.18E-127 | QP |
| THOP1 | 0.27033036 | 7.39E-107 | 1.05E-102 | QP |
| TNFAIP8L1 | 0.27024464 | 2.07E-69 | 2.93E-65 | QP |
| TXNDC12 | 0.26914193 | 2.33E-92 | 3.29E-88 | QP |
| FANCA | 0.26896671 | 1.66E-134 | 2.35E-130 | QP |
| SASS6 | 0.2684482 | 4.19E-128 | 5.93E-124 | QP |
| ACOT9 | 0.26836846 | 1.08E-114 | 1.53E-110 | QP |
| DONSON | 0.26820661 | 1.50E-126 | 2.12E-122 | QP |
| TTF2 | 0.2676394 | 1.02E-116 | 1.44E-112 | QP |
| AKT1 | 0.26756808 | 7.60E-95 | 1.08E-90 | QP |
| RHOC | 0.26739268 | 7.75E-108 | 1.10E-103 | QP |
| DHCR24 | 0.26721099 | 3.55E-83 | 5.02E-79 | QP |

|  |  |  |  |
| --- | --- | --- | --- |
| GMPS | 0.26709703 | 5.46E-78 | 7.73E-74 QP |
| ITGAE | 0.26697669 | 1.78E-101 | 2.52E-97 QP |
| MYCBP | 0.26677095 | 9.54E-94 | 1.35E-89 QP |
| PITPNC1 | 0.26664418 | 1.95E-80 | 2.76E-76 QP |
| PRIM2 | 0.26646829 | 3.53E-115 | 4.99E-111 QP |
| ANP32A | 0.26623234 | 4.96E-129 | 7.02E-125 QP |
| EWSR1 | 0.26581914 | 1.56E-106 | 2.20E-102 QP |
| ADGRE5 | 0.26570754 | 1.60E-54 | 2.27E-50 QP |
| RCCD1 | 0.26508477 | 2.20E-101 | 3.11E-97 QP |
| MCM6 | 0.26497053 | 1.71E-60 | 2.42E-56 QP |
| SPA17 | 0.26486347 | 4.40E-90 | 6.22E-86 QP |
| ACTN1 | 0.26465767 | 2.22E-116 | 3.14E-112 QP |
| FOPNL | 0.26461147 | 3.81E-71 | 5.39E-67 QP |
| HNRNPC | 0.26451903 | 1.17E-163 | 1.65E-159 QP |
| AL590617.2 | 0.2640684 | 3.11E-93 | 4.41E-89 QP |
| PLS3 | 0.26392849 | 2.37E-78 | 3.36E-74 QP |
| G3BP1 | 0.26387841 | 7.83E-107 | 1.11E-102 QP |
| CYCS | 0.26353069 | 3.11E-192 | 4.40E-188 QP |
| NCAPD3 | 0.26306297 | 2.89E-122 | 4.08E-118 QP |
| RMI2 | 0.26271709 | 6.56E-97 | 9.27E-93 QP |
| EPB41L2 | 0.26264751 | 7.81E-93 | 1.10E-88 QP |
| EXOSC3 | 0.26239544 | 1.43E-99 | 2.02E-95 QP |
| NET1 | 0.26238857 | 4.48E-105 | 6.34E-101 QP |
| TMEM97 | 0.2610202 | 1.54E-81 | 2.17E-77 QP |
| ENSA | 0.26088522 | 4.51E-119 | 6.38E-115 QP |
| LSM7 | 0.26086978 | 1.62E-142 | 2.29E-138 QP |
| NUDC | 0.26074022 | 9.42E-137 | 1.33E-132 QP |
| TDP1 | 0.26054363 | 1.27E-101 | 1.79E-97 QP |
| CCHCR1 | 0.26047582 | 2.59E-89 | 3.66E-85 QP |
| TCEA1 | 0.25996485 | 1.78E-132 | 2.52E-128 QP |
| HAS2 | 0.25969158 | 1.11E-37 | 1.56E-33 QP |
| RNPS1 | 0.2593429 | 4.84E-122 | 6.85E-118 QP |
| UNG | 0.2592837 | 3.33E-42 | 4.72E-38 QP |
| BZW1 | 0.2591893 | 8.53E-125 | 1.21E-120 QP |
| YEATS4 | 0.25906278 | 3.81E-75 | 5.38E-71 QP |
| COX20 | 0.25896741 | 1.04E-89 | 1.47E-85 QP |
| THOC3 | 0.25889308 | 4.95E-89 | 7.00E-85 QP |
| KHSRP | 0.25884094 | 9.29E-89 | 1.31E-84 QP |
| RPS6KA5 | 0.25816834 | 4.53E-78 | 6.40E-74 QP |
| SUZ12 | 0.25803524 | 3.37E-76 | 4.77E-72 QP |
| MRPL51 | 0.2578899 | 4.38E-237 | 6.19E-233 QP |
| CCNF | 0.25756521 | 4.52E-177 | 6.40E-173 QP |
| DESI2 | 0.25747417 | 4.40E-82 | 6.22E-78 QP |
| PARP1 | 0.25732577 | 9.27E-85 | 1.31E-80 QP |
| SH3GL1 | 0.25718605 | 4.06E-86 | 5.74E-82 QP |
| RAD1 | 0.25688451 | 1.29E-83 | 1.83E-79 QP |

|  |  |  |  |
| --- | --- | --- | --- |
| DR1 | 0.25675341 | 6.37E-87 | 9.01E-83 QP |
| FAM24B | 0.25660353 | 1.24E-58 | 1.76E-54 QP |
| UBE2A | 0.25633682 | 7.24E-111 | 1.02E-106 QP |
| NSMCE4A | 0.25605175 | 1.02E-90 | 1.44E-86 QP |
| GNG4 | 0.25596452 | 7.47E-93 | 1.06E-88 QP |
| DIAPH1 | 0.25587852 | 8.66E-86 | 1.23E-81 QP |
| CDC5L | 0.25586382 | 8.86E-104 | 1.25E-99 QP |
| SNRPC | 0.25543989 | 1.27E-133 | 1.79E-129 QP |
| CEP97 | 0.25517524 | 6.74E-103 | 9.53E-99 QP |
| TBL1XR1 | 0.25459926 | 2.15E-69 | 3.04E-65 QP |
| HAUS6 | 0.25453665 | 1.46E-95 | 2.06E-91 QP |
| MSN | 0.25442761 | 1.98E-87 | 2.80E-83 QP |
| CEP57 | 0.25419482 | 5.90E-82 | 8.35E-78 QP |
| CENPI | 0.25399797 | 2.54E-176 | 3.60E-172 QP |
| ID4 | 0.2539663 | 2.83E-38 | 4.01E-34 QP |
| EIF5 | 0.25356482 | 6.60E-120 | 9.34E-116 QP |
| CTNNAL1 | 0.25287749 | 3.91E-82 | 5.53E-78 QP |
| IGFBP3 | 0.25276402 | 1.20E-49 | 1.70E-45 QP |
| WDR76 | 0.25268924 | 1.13E-91 | 1.59E-87 QP |
| ANXA7 | 0.25245924 | 7.96E-86 | 1.13E-81 QP |
| ADAMTS1 | 0.25181188 | 3.00E-38 | 4.25E-34 QP |
| TMA7 | 0.25174007 | 3.16E-124 | 4.48E-120 QP |
| ANAPC7 | 0.2502995 | 7.70E-78 | 1.09E-73 QP |
| TXNIP | 1.17393532 | 2.00E-206 | 2.83E-202 PQ |
| HLA-DRB5 | 0.92399654 | 6.65E-138 | 9.41E-134 PQ |
| IFITM3 | 0.9131442 | 2.08E-227 | 2.94E-223 PQ |
| NDRG1 | 0.85323179 | 5.85E-122 | 8.27E-118 PQ |
| DKK3 | 0.79719267 | 1.69E-110 | 2.40E-106 PQ |
| HLA-DRB1 | 0.78672691 | 1.49E-51 | 2.11E-47 PQ |
| HLA-DRA | 0.78613575 | 8.11E-51 | 1.15E-46 PQ |
| ST3GAL5 | 0.78243004 | 6.03E-97 | 8.53E-93 PQ |
| HLA-DQB1 | 0.77637027 | 8.36E-141 | 1.18E-136 PQ |
| CTSH | 0.76868436 | 2.89E-52 | 4.09E-48 PQ |
| COL6A2 | 0.76198141 | 3.90E-137 | 5.52E-133 PQ |
| CD74 | 0.74663622 | 7.61E-54 | 1.08E-49 PQ |
| SCG2 | 0.71891339 | 1.61E-51 | 2.28E-47 PQ |
| S100A1 | 0.68372082 | 1.27E-80 | 1.80E-76 PQ |
| COL6A1 | 0.66310112 | 1.11E-111 | 1.58E-107 PQ |
| SERPINE2 | 0.66184984 | 1.54E-56 | 2.18E-52 PQ |
| ZFP36 | 0.64378683 | 1.23E-38 | 1.74E-34 PQ |
| GPNMB | 0.63846413 | 2.91E-90 | 4.12E-86 PQ |
| SERPINE1 | 0.62512265 | 2.33E-18 | 3.30E-14 PQ |
| TNFRSF21 | 0.62069566 | 1.35E-71 | 1.92E-67 PQ |
| PSAP | 0.62064742 | 3.13E-159 | 4.43E-155 PQ |
| BBC3 | 0.607633 | 4.95E-176 | 7.00E-172 PQ |
| RPS10 | 0.60188808 | 7.08E-202 | 1.00E-197 PQ |

|  |  |  |  |
| --- | --- | --- | --- |
| DUSP4 | 0.59177154 | 2.51E-110 | 3.55E-106 PQ |
| HLA-DPA1 | 0.58191651 | 3.33E-36 | 4.71E-32 PQ |
| KCTD12 | 0.58141118 | 5.54E-119 | 7.84E-115 PQ |
| JUNB | 0.57995375 | 1.80E-29 | 2.54E-25 PQ |
| MARCKSL1 | 0.57711986 | 1.27E-62 | 1.80E-58 PQ |
| IER3 | 0.57563614 | 6.06E-57 | 8.58E-53 PQ |
| MALAT1 | 0.55767244 | 2.71E-65 | 3.83E-61 PQ |
| SLC20A1 | 0.55578626 | 1.77E-93 | 2.51E-89 PQ |
| CTSB | 0.5545917 | 1.84E-133 | 2.61E-129 PQ |
| NPC2 | 0.55021485 | 3.97E-110 | 5.61E-106 PQ |
| PHLDA3 | 0.54976881 | 2.38E-137 | 3.36E-133 PQ |
| AHNAK2 | 0.54971005 | 1.24E-77 | 1.75E-73 PQ |
| MT-ND3 | 0.54678884 | 2.29E-56 | 3.24E-52 PQ |
| A1BG | 0.54348959 | 4.20E-270 | 5.94E-266 PQ |
| FBXO32 | 0.54185372 | 9.58E-55 | 1.36E-50 PQ |
| MTRNR2L12 | 0.53977207 | 8.76E-33 | 1.24E-28 PQ |
| CCDC85B | 0.53720134 | 6.44E-175 | 9.11E-171 PQ |
| ITGB4 | 0.5318023 | 3.05E-280 | 4.32E-276 PQ |
| FN1 | 0.52831805 | 6.26E-85 | 8.86E-81 PQ |
| MGP | 0.51090165 | 4.23E-16 | 5.98E-12 PQ |
| SLC1A3 | 0.50889722 | 3.67E-74 | 5.19E-70 PQ |
| TRNP1 | 0.50739629 | 8.40E-114 | 1.19E-109 PQ |
| DUSP23 | 0.50200463 | 9.60E-103 | 1.36E-98 PQ |
| CADPS | 0.49272303 | 8.87E-108 | 1.25E-103 PQ |
| DUBR | 0.49018367 | 3.12E-110 | 4.41E-106 PQ |
| TIMP3 | 0.48719321 | 1.04E-10 | 1.48E-06 PQ |
| GAS5 | 0.47839082 | 7.60E-108 | 1.08E-103 PQ |
| PNRC1 | 0.47800566 | 4.81E-99 | 6.80E-95 PQ |
| NRN1 | 0.47670508 | 2.32E-109 | 3.28E-105 PQ |
| SNHG32 | 0.47044631 | 4.91E-85 | 6.95E-81 PQ |
| GADD45A | 0.46941614 | 3.31E-52 | 4.68E-48 PQ |
| MMP14 | 0.4675811 | 1.25E-42 | 1.77E-38 PQ |
| SFRP1 | 0.46751526 | 6.67E-25 | 9.43E-21 PQ |
| GYPC | 0.46465513 | 6.06E-144 | 8.57E-140 PQ |
| IFITM2 | 0.45444969 | 6.00E-95 | 8.49E-91 PQ |
| SSBP4 | 0.44905862 | 7.18E-99 | 1.02E-94 PQ |
| NUPR1 | 0.44815581 | 8.52E-58 | 1.21E-53 PQ |
| HLA-DPB1 | 0.4466758 | 4.47E-31 | 6.33E-27 PQ |
| DUSP6 | 0.43843267 | 1.85E-52 | 2.62E-48 PQ |
| GABARAPL1 | 0.43706712 | 3.85E-85 | 5.44E-81 PQ |
| GXYLT2 | 0.4369616 | 5.38E-68 | 7.61E-64 PQ |
| BMP4 | 0.43536982 | 1.19E-77 | 1.69E-73 PQ |
| DBP | 0.43441806 | 3.92E-87 | 5.54E-83 PQ |
| LRP1 | 0.43383612 | 1.42E-87 | 2.01E-83 PQ |
| CDKN2B | 0.43346872 | 6.01E-101 | 8.51E-97 PQ |
| POU2F2 | 0.43171041 | 2.08E-48 | 2.94E-44 PQ |

|  |  |  |  |
| --- | --- | --- | --- |
| CYSTM1 | 0.43000732 | 1.14E-72 | 1.61E-68 PQ |
| TMEM158 | 0.42985947 | 2.10E-38 | 2.97E-34 PQ |
| EGR1 | 0.42895622 | 7.47E-13 | 1.06E-08 PQ |
| NEU1 | 0.42699281 | 2.10E-95 | 2.97E-91 PQ |
| SDC2 | 0.42176191 | 1.56E-40 | 2.20E-36 PQ |
| SUGCT | 0.4209732 | 1.21E-49 | 1.71E-45 PQ |
| SNHG5 | 0.42088168 | 1.22E-78 | 1.73E-74 PQ |
| PRSS23 | 0.41982398 | 1.19E-27 | 1.69E-23 PQ |
| PLD3 | 0.41370033 | 9.55E-85 | 1.35E-80 PQ |
| HLA-E | 0.41152393 | 1.82E-96 | 2.58E-92 PQ |
| PAPPA | 0.41115342 | 1.22E-114 | 1.73E-110 PQ |
| HLA-DMA | 0.41112307 | 2.89E-52 | 4.09E-48 PQ |
| MMP2 | 0.41048131 | 3.03E-66 | 4.28E-62 PQ |
| RGS4 | 0.40781385 | 2.55E-69 | 3.61E-65 PQ |
| HOXB6 | 0.40722187 | 1.74E-65 | 2.46E-61 PQ |
| KRT8 | 0.40684193 | 1.91E-97 | 2.71E-93 PQ |
| DDIT4 | 0.40269506 | 5.99E-57 | 8.47E-53 PQ |
| DUSP1 | 0.40204306 | 2.41E-20 | 3.40E-16 PQ |
| GRN | 0.40172247 | 1.56E-82 | 2.21E-78 PQ |
| TYMP | 0.39954501 | 1.54E-77 | 2.18E-73 PQ |
| PPL | 0.39830418 | 5.15E-107 | 7.29E-103 PQ |
| YPEL3 | 0.39643913 | 1.41E-75 | 2.00E-71 PQ |
| OLFML2A | 0.3920514 | 9.53E-54 | 1.35E-49 PQ |
| ACTC1 | 0.391285 | 2.42E-11 | 3.43E-07 PQ |
| MLPH | 0.38979173 | 9.68E-68 | 1.37E-63 PQ |
| GLUL | 0.38884568 | 2.10E-43 | 2.97E-39 PQ |
| FAM20C | 0.38451541 | 7.22E-44 | 1.02E-39 PQ |
| GSN | 0.38213765 | 1.16E-101 | 1.64E-97 PQ |
| ITGA3 | 0.38062003 | 7.90E-55 | 1.12E-50 PQ |
| SPRY1 | 0.38026729 | 2.34E-19 | 3.32E-15 PQ |
| FRMD4A | 0.38009218 | 1.09E-58 | 1.55E-54 PQ |
| SPARC | 0.37964216 | 1.21E-80 | 1.72E-76 PQ |
| HLA-DMB | 0.37497485 | 7.68E-68 | 1.09E-63 PQ |
| ZFAS1 | 0.37306979 | 1.33E-122 | 1.88E-118 PQ |
| TSC22D1 | 0.37123602 | 2.57E-49 | 3.64E-45 PQ |
| GRINA | 0.37086693 | 1.45E-99 | 2.06E-95 PQ |
| IGFBP4 | 0.36670304 | 1.03E-28 | 1.46E-24 PQ |
| ANXA4 | 0.36517303 | 2.80E-98 | 3.96E-94 PQ |
| CEBPD | 0.36488695 | 4.17E-33 | 5.90E-29 PQ |
| ITGB5 | 0.36450887 | 7.01E-67 | 9.92E-63 PQ |
| MT-ND4L | 0.36449618 | 8.00E-41 | 1.13E-36 PQ |
| CTSD | 0.36446911 | 2.96E-78 | 4.19E-74 PQ |
| ASAH1 | 0.36018804 | 1.78E-67 | 2.52E-63 PQ |
| LRRC75A | 0.35892239 | 3.60E-48 | 5.09E-44 PQ |
| MMP24OS | 0.35758171 | 3.05E-84 | 4.32E-80 PQ |
| GABARAP | 0.35694775 | 4.08E-120 | 5.78E-116 PQ |

|  |  |  |  |
| --- | --- | --- | --- |
| COL18A1 | 0.35655785 | 1.93E-55 | 2.73E-51 PQ |
| RPL34 | 0.35540621 | 4.62E-187 | 6.54E-183 PQ |
| TSPAN14 | 0.3539654 | 3.87E-85 | 5.48E-81 PQ |
| CTSK | 0.35392371 | 1.06E-22 | 1.49E-18 PQ |
| BNIP3L | 0.35305044 | 1.23E-78 | 1.74E-74 PQ |
| LAMB2 | 0.35252776 | 1.74E-65 | 2.47E-61 PQ |
| TMEM205 | 0.35217913 | 2.12E-85 | 3.00E-81 PQ |
| METRN | 0.34978411 | 6.43E-44 | 9.10E-40 PQ |
| CSTB | 0.3493912 | 4.38E-108 | 6.20E-104 PQ |
| SERPINF1 | 0.34915646 | 3.89E-31 | 5.50E-27 PQ |
| HLA-B | 0.34843126 | 1.34E-30 | 1.90E-26 PQ |
| EBF2 | 0.34597706 | 4.29E-98 | 6.07E-94 PQ |
| JUN | 0.34535372 | 7.11E-18 | 1.01E-13 PQ |
| SAT1 | 0.34522352 | 7.14E-29 | 1.01E-24 PQ |
| FAM89B | 0.34447279 | 3.44E-76 | 4.87E-72 PQ |
| LGALS3 | 0.34346184 | 4.32E-142 | 6.12E-138 PQ |
| EEF1A1 | 0.34130739 | 1.02E-132 | 1.44E-128 PQ |
| RPS27 | 0.33859702 | 4.41E-176 | 6.25E-172 PQ |
| VAT1 | 0.33783277 | 6.74E-81 | 9.54E-77 PQ |
| PBXIP1 | 0.33759834 | 3.17E-35 | 4.49E-31 PQ |
| RNASEK | 0.33660424 | 6.98E-123 | 9.88E-119 PQ |
| HMGA2 | 0.33552603 | 9.12E-24 | 1.29E-19 PQ |
| LAPTM4A | 0.33294804 | 2.69E-29 | 3.80E-25 PQ |
| DYNLRB1 | 0.32932465 | 3.66E-89 | 5.18E-85 PQ |
| SNED1 | 0.32889729 | 6.49E-51 | 9.19E-47 PQ |
| FOXN3 | 0.32810656 | 8.64E-81 | 1.22E-76 PQ |
| RPL31 | 0.3277351 | 1.15E-130 | 1.63E-126 PQ |
| DAAM2 | 0.32688233 | 4.95E-32 | 7.00E-28 PQ |
| RPS25 | 0.3267693 | 7.83E-180 | 1.11E-175 PQ |
| VPS51 | 0.32607589 | 1.25E-79 | 1.77E-75 PQ |
| KCNN4 | 0.32553836 | 1.72E-41 | 2.44E-37 PQ |
| FTL | 0.32376997 | 1.48E-109 | 2.09E-105 PQ |
| CBX6 | 0.32218676 | 3.72E-76 | 5.26E-72 PQ |
| SPON2 | 0.32011702 | 4.49E-19 | 6.36E-15 PQ |
| RPL38 | 0.31890637 | 5.83E-165 | 8.25E-161 PQ |
| HCFC1R1 | 0.31843509 | 2.29E-48 | 3.24E-44 PQ |
| MXI1 | 0.31804997 | 1.98E-92 | 2.80E-88 PQ |
| CDKN1A | 0.3165233 | 3.86E-08 | 0.00054593 PQ |
| EPB41L4A-A5 | 0.31647746 | 7.19E-77 | 1.02E-72 PQ |
| LAMA5 | 0.31509791 | 8.37E-56 | 1.18E-51 PQ |
| CNIH3 | 0.31487563 | 1.28E-62 | 1.82E-58 PQ |
| PTPA | 0.31461017 | 6.90E-54 | 9.77E-50 PQ |
| SVIL | 0.31448852 | 7.84E-56 | 1.11E-51 PQ |
| NPDC1 | 0.31409746 | 8.46E-73 | 1.20E-68 PQ |
| MT1X | 0.31215519 | 1.69E-37 | 2.40E-33 PQ |
| CMTM7 | 0.31148786 | 9.49E-47 | 1.34E-42 PQ |

|  |  |  |  |
| --- | --- | --- | --- |
| IGF2R | 0.31075736 | 1.08E-60 | 1.52E-56 PQ |
| TMEM163 | 0.30978695 | 2.55E-56 | 3.61E-52 PQ |
| IFITM1 | 0.30905858 | 4.08E-55 | 5.78E-51 PQ |
| TSC22D4 | 0.30859817 | 5.23E-69 | 7.39E-65 PQ |
| HES1 | 0.30813346 | 1.65E-06 | 0.0233244 PQ |
| KLHDC8B | 0.30644448 | 2.82E-24 | 3.98E-20 PQ |
| LUM | 0.30600687 | 2.77E-05 | 0.39227465 PQ |
| TSC22D3 | 0.30533882 | 1.75E-27 | 2.47E-23 PQ |
| RENBP | 0.30385573 | 9.17E-98 | 1.30E-93 PQ |
| RPS28 | 0.30237486 | 8.41E-199 | 1.19E-194 PQ |
| TOMM7 | 0.30111168 | 2.32E-148 | 3.28E-144 PQ |
| RPS12 | 0.30047163 | 1.70E-128 | 2.40E-124 PQ |
| PAQR8 | 0.29967457 | 5.64E-66 | 7.98E-62 PQ |
| KRT19 | 0.29933492 | 4.24E-64 | 6.00E-60 PQ |
| CITED2 | 0.29863307 | 4.91E-30 | 6.94E-26 PQ |
| COMMD6 | 0.29698281 | 1.90E-117 | 2.69E-113 PQ |
| CDKN1C | 0.29532929 | 3.94E-61 | 5.57E-57 PQ |
| RPL36A | 0.29415878 | 1.20E-157 | 1.69E-153 PQ |
| PLAAT4 | 0.29411597 | 8.02E-61 | 1.13E-56 PQ |
| CHP1 | 0.29345734 | 1.11E-61 | 1.57E-57 PQ |
| IFI27L2 | 0.2933934 | 1.37E-85 | 1.94E-81 PQ |
| GAS6 | 0.29336437 | 1.22E-43 | 1.72E-39 PQ |
| CST3 | 0.29306418 | 2.34E-40 | 3.32E-36 PQ |
| RPL23 | 0.29177172 | 5.37E-105 | 7.59E-101 PQ |
| PDK4 | 0.29126114 | 1.47E-30 | 2.08E-26 PQ |
| BCL2L1 | 0.29094462 | 1.24E-47 | 1.75E-43 PQ |
| SERINC2 | 0.29000361 | 9.89E-18 | 1.40E-13 PQ |
| HLA-DQA1 | 0.28858994 | 1.23E-78 | 1.74E-74 PQ |
| TTN | 0.28742452 | 7.45E-15 | 1.05E-10 PQ |
| CTSA | 0.28694555 | 1.70E-48 | 2.40E-44 PQ |
| PDXK | 0.28599288 | 6.34E-29 | 8.97E-25 PQ |
| RRAS | 0.28576262 | 5.39E-52 | 7.63E-48 PQ |
| ITPR3 | 0.28544362 | 7.56E-33 | 1.07E-28 PQ |
| RHBDF1 | 0.28534454 | 5.17E-41 | 7.32E-37 PQ |
| FABP3 | 0.28421514 | 1.63E-60 | 2.30E-56 PQ |
| F2R | 0.28414196 | 6.09E-43 | 8.61E-39 PQ |
| EIF4A2 | 0.2834994 | 2.93E-89 | 4.14E-85 PQ |
| RPL37A | 0.28234276 | 1.24E-194 | 1.76E-190 PQ |
| COX7C | 0.28204851 | 7.39E-108 | 1.05E-103 PQ |
| STAT2 | 0.28089064 | 6.30E-59 | 8.91E-55 PQ |
| GLMP | 0.28069322 | 7.12E-58 | 1.01E-53 PQ |
| SLC6A8 | 0.28055377 | 7.91E-60 | 1.12E-55 PQ |
| BTG1 | 0.28028411 | 0.0019953 | 1 PQ |
| SLC27A1 | 0.2796071 | 7.84E-53 | 1.11E-48 PQ |
| CDKN2A | 0.27941138 | 9.85E-64 | 1.39E-59 PQ |
| SIGIRR | 0.27935645 | 5.37E-61 | 7.60E-57 PQ |

|  |  |  |  |
| --- | --- | --- | --- |
| TPBG | 0.27900346 | 1.30E-23 | 1.84E-19 PQ |
| ANXA11 | 0.27825026 | 1.17E-61 | 1.66E-57 PQ |
| EEF2 | 0.27723297 | 8.47E-126 | 1.20E-121 PQ |
| FZD8 | 0.27703911 | 1.07E-36 | 1.52E-32 PQ |
| SCD | 0.27576787 | 7.61E-77 | 1.08E-72 PQ |
| RPL27 | 0.27422166 | 2.07E-144 | 2.93E-140 PQ |
| SPRY4 | 0.27411216 | 1.18E-27 | 1.67E-23 PQ |
| COX6B1 | 0.2734318 | 1.83E-109 | 2.60E-105 PQ |
| LETMD1 | 0.27314894 | 4.22E-43 | 5.97E-39 PQ |
| SIRT2 | 0.27312737 | 8.96E-21 | 1.27E-16 PQ |
| KIFAP3 | 0.27308845 | 2.78E-47 | 3.93E-43 PQ |
| S100A4 | 0.27284757 | 4.18E-08 | 0.00059185 PQ |
| IER5L | 0.27277815 | 1.67E-53 | 2.37E-49 PQ |
| AC100810.1 | 0.27212037 | 4.35E-57 | 6.16E-53 PQ |
| OST4 | 0.27084631 | 1.23E-104 | 1.74E-100 PQ |
| ZSCAN18 | 0.27076525 | 4.02E-157 | 5.69E-153 PQ |
| FKBP1A | 0.27055284 | 1.45E-70 | 2.05E-66 PQ |
| NINJ1 | 0.26989186 | 1.32E-32 | 1.87E-28 PQ |
| EMC10 | 0.26942766 | 1.29E-52 | 1.83E-48 PQ |
| CTNNB1 | 0.26759096 | 4.40E-27 | 6.23E-23 PQ |
| IFI16 | 0.26752431 | 1.11E-57 | 1.56E-53 PQ |
| MFAP2 | 0.26463327 | 1.93E-38 | 2.74E-34 PQ |
| FHL3 | 0.26378788 | 1.44E-52 | 2.04E-48 PQ |
| ARL6IP5 | 0.26219636 | 1.28E-102 | 1.81E-98 PQ |
| LGMN | 0.26171936 | 1.06E-65 | 1.51E-61 PQ |
| SMIM29 | 0.26018267 | 2.01E-35 | 2.84E-31 PQ |
| SQOR | 0.25985187 | 1.15E-56 | 1.63E-52 PQ |
| COMT | 0.25921314 | 8.71E-69 | 1.23E-64 PQ |
| MXD4 | 0.2587203 | 9.54E-53 | 1.35E-48 PQ |
| FNBP1L | 0.25763075 | 4.22E-43 | 5.97E-39 PQ |
| QPCT | 0.25657405 | 6.38E-43 | 9.03E-39 PQ |
| TGFBI | 0.25650996 | 1.42E-36 | 2.01E-32 PQ |
| CA12 | 0.2564076 | 1.14E-114 | 1.61E-110 PQ |
| LINC00520 | 0.25585142 | 1.38E-132 | 1.96E-128 PQ |
| MAP1LC3B | 0.25574925 | 2.29E-42 | 3.24E-38 PQ |
| ADM | 0.2538008 | 5.44E-24 | 7.70E-20 PQ |
| HSD17B14 | 0.25313198 | 4.15E-78 | 5.88E-74 PQ |
| SLC25A23 | 0.25298992 | 1.76E-50 | 2.49E-46 PQ |
| TSPYL1 | 0.25289411 | 8.41E-42 | 1.19E-37 PQ |
| CTXN1 | 0.25241615 | 1.56E-75 | 2.21E-71 PQ |
| COL1A1 | 0.25222436 | 6.23E-21 | 8.82E-17 PQ |
| CPQ | 0.25110391 | 1.29E-61 | 1.83E-57 PQ |
| RPL36 | 0.25076698 | 1.70E-154 | 2.41E-150 PQ |
| ATP1A1 | 0.25064425 | 1.51E-41 | 2.14E-37 PQ |
| DDIT3 | 0.25016525 | 8.12E-40 | 1.15E-35 PQ |
| MKI67 | 1.01810102 | 0 | 0 PP |

|  |  |  |  |  |
| --- | --- | --- | --- | --- |
| CENPF | 0.96930366 | 4.34E-286 | 6.14E-282 | PP |
| HIST1H4C | 0.9228054 | 5.64E-140 | 7.98E-136 | PP |
| ASPM | 0.83194455 | 1.02E-238 | 1.44E-234 | PP |
| CDKN3 | 0.80495573 | 2.28E-250 | 3.22E-246 | PP |
| SMC4 | 0.79941637 | 1.21E-239 | 1.71E-235 | PP |
| HIST1H1B | 0.78861359 | 1.60E-104 | 2.27E-100 | PP |
| HIST1H3D | 0.77686218 | 2.32E-64 | 3.29E-60 | PP |
| CCNB1 | 0.7762336 | 1.78E-161 | 2.52E-157 | PP |
| TPX2 | 0.77215884 | 4.91E-227 | 6.95E-223 | PP |
| PRC1 | 0.76323606 | 3.49E-245 | 4.93E-241 | PP |
| TOP2A | 0.73497436 | 8.73E-225 | 1.24E-220 | PP |
| CEP55 | 0.73458154 | 7.69E-240 | 1.09E-235 | PP |
| RRM2 | 0.72443397 | 5.20E-220 | 7.36E-216 | PP |
| PBK | 0.70429886 | 4.52E-238 | 6.40E-234 | PP |
| KIF20B | 0.70289104 | 7.69E-171 | 1.09E-166 | PP |
| PTTG1 | 0.69013791 | 9.66E-219 | 1.37E-214 | PP |
| PCLAF | 0.68292704 | 2.21E-182 | 3.13E-178 | PP |
| DLGAP5 | 0.67152781 | 3.52E-177 | 4.98E-173 | PP |
| NUSAP1 | 0.66744554 | 8.16E-215 | 1.15E-210 | PP |
| BIRC5 | 0.66571921 | 1.75E-216 | 2.47E-212 | PP |
| STC1 | 0.66446617 | 1.21E-115 | 1.72E-111 | PP |
| CCNB2 | 0.66172831 | 2.10E-183 | 2.96E-179 | PP |
| TUBA1B | 0.65935557 | 8.32E-215 | 1.18E-210 | PP |
| NUF2 | 0.65781028 | 5.75E-181 | 8.13E-177 | PP |
| ANLN | 0.65666317 | 9.30E-200 | 1.32E-195 | PP |
| CDK1 | 0.65128038 | 5.23E-166 | 7.40E-162 | PP |
| UBE2S | 0.64088673 | 4.57E-186 | 6.46E-182 | PP |
| TYMS | 0.63948079 | 3.48E-181 | 4.93E-177 | PP |
| MOK | 0.6329669 | 7.94E-187 | 1.12E-182 | PP |
| TK1 | 0.62590683 | 7.62E-213 | 1.08E-208 | PP |
| NCAPG | 0.62050117 | 1.74E-190 | 2.47E-186 | PP |
| DIAPH3 | 0.60784913 | 7.28E-206 | 1.03E-201 | PP |
| CDC20 | 0.59890798 | 3.74E-160 | 5.29E-156 | PP |
| CDCA3 | 0.59634515 | 1.20E-170 | 1.69E-166 | PP |
| SMC2 | 0.59291049 | 1.14E-179 | 1.61E-175 | PP |
| GTSE1 | 0.59230961 | 1.44E-184 | 2.04E-180 | PP |
| CENPE | 0.59177611 | 3.14E-139 | 4.44E-135 | PP |
| HIST1H1A | 0.58765403 | 2.13E-67 | 3.02E-63 | PP |
| H2AFZ | 0.58327608 | 2.15E-221 | 3.04E-217 | PP |
| HIST1H1E | 0.58318637 | 1.20E-46 | 1.70E-42 | PP |
| TMPO | 0.58208037 | 9.43E-177 | 1.33E-172 | PP |
| UBE2T | 0.58046247 | 7.15E-142 | 1.01E-137 | PP |
| UBE2C | 0.5742058 | 4.15E-158 | 5.88E-154 | PP |
| CLSPN | 0.57038045 | 1.37E-118 | 1.93E-114 | PP |
| DHFR | 0.5662097 | 4.51E-126 | 6.38E-122 | PP |
| ZWINT | 0.56341381 | 1.86E-158 | 2.63E-154 | PP |

|  |  |  |  |  |
| --- | --- | --- | --- | --- |
| ATAD2 | 0.56336132 | 2.53E-117 | 3.58E-113 | PP |
| RRM1 | 0.55930161 | 6.12E-138 | 8.66E-134 | PP |
| KIF11 | 0.55885818 | 1.38E-170 | 1.96E-166 | PP |
| DNMT1 | 0.55648883 | 5.16E-175 | 7.31E-171 | PP |
| CKS1B | 0.55494717 | 3.95E-153 | 5.59E-149 | PP |
| CENPM | 0.55427809 | 1.83E-167 | 2.59E-163 | PP |
| TACC3 | 0.5495279 | 3.46E-169 | 4.89E-165 | PP |
| HIST1H3B | 0.54228323 | 6.78E-79 | 9.60E-75 | PP |
| DTYMK | 0.5391168 | 3.35E-209 | 4.74E-205 | PP |
| FOXMI | 0.53907417 | 1.35E-190 | 1.91E-186 | PP |
| LMNB2 | 0.53834946 | 3.05E-179 | 4.32E-175 | PP |
| MAD2L1 | 0.5378096 | 1.16E-158 | 1.65E-154 | PP |
| AURKB | 0.53665138 | 1.34E-136 | 1.89E-132 | PP |
| CENPK | 0.53204738 | 3.13E-147 | 4.42E-143 | PP |
| NCAPD2 | 0.53197416 | 7.51E-164 | 1.06E-159 | PP |
| KIF23 | 0.53083413 | 1.67E-131 | 2.36E-127 | PP |
| SFRP1 | 0.52956438 | 1.54E-111 | 2.18E-107 | PP |
| CIP2A | 0.52896155 | 3.63E-168 | 5.13E-164 | PP |
| TUBA1C | 0.52627766 | 2.28E-134 | 3.22E-130 | PP |
| CIT | 0.5211461 | 4.03E-156 | 5.71E-152 | PP |
| SPC25 | 0.51889441 | 3.00E-151 | 4.24E-147 | PP |
| MIS18BP1 | 0.51625824 | 2.29E-122 | 3.24E-118 | PP |
| KIFC1 | 0.50922057 | 3.12E-134 | 4.41E-130 | PP |
| RAD21 | 0.5088499 | 2.59E-133 | 3.66E-129 | PP |
| GGH | 0.50639498 | 2.08E-175 | 2.94E-171 | PP |
| HMGB3 | 0.50136542 | 6.47E-127 | 9.15E-123 | PP |
| DEK | 0.50094351 | 4.02E-194 | 5.68E-190 | PP |
| SGO2 | 0.50084578 | 4.97E-107 | 7.04E-103 | PP |
| PLK1 | 0.50048203 | 1.07E-99 | 1.52E-95 | PP |
| HMMR | 0.50022575 | 1.09E-116 | 1.54E-112 | PP |
| FEN1 | 0.49829748 | 3.41E-104 | 4.83E-100 | PP |
| HMGB2 | 0.49801631 | 1.44E-114 | 2.03E-110 | PP |
| ECT2 | 0.49580553 | 4.84E-114 | 6.84E-110 | PP |
| TUBB4B | 0.4942957 | 3.28E-124 | 4.65E-120 | PP |
| DDX39A | 0.49363065 | 2.18E-177 | 3.09E-173 | PP |
| SYNE2 | 0.49175753 | 5.56E-87 | 7.87E-83 | PP |
| DEPDC1 | 0.49120172 | 1.34E-136 | 1.89E-132 | PP |
| FOSL1 | 0.49052835 | 1.91E-122 | 2.70E-118 | PP |
| TUBB | 0.49011013 | 1.06E-150 | 1.51E-146 | PP |
| USP1 | 0.49007156 | 1.59E-122 | 2.25E-118 | PP |
| RAD51AP1 | 0.48881502 | 2.98E-134 | 4.22E-130 | PP |
| ANP32E | 0.48716619 | 1.20E-148 | 1.69E-144 | PP |
| C12orf75 | 0.48604489 | 4.21E-178 | 5.95E-174 | PP |
| KIF4A | 0.48390864 | 3.75E-154 | 5.31E-150 | PP |
| NSD2 | 0.47978632 | 9.55E-147 | 1.35E-142 | PP |
| TUBB6 | 0.47507193 | 2.86E-124 | 4.04E-120 | PP |

|  |  |  |  |  |
| --- | --- | --- | --- | --- |
| MAP1B | 0.47431457 | 1.18E-141 | 1.67E-137 | PP |
| PHF19 | 0.47307793 | 5.77E-154 | 8.16E-150 | PP |
| NCEH1 | 0.4723219 | 1.63E-107 | 2.31E-103 | PP |
| NEK2 | 0.46855685 | 2.81E-127 | 3.97E-123 | PP |
| SGO1 | 0.46808622 | 5.17E-137 | 7.31E-133 | PP |
| TMEM106C | 0.46801464 | 2.17E-106 | 3.06E-102 | PP |
| TROAP | 0.4667762 | 4.56E-137 | 6.45E-133 | PP |
| CKAP2L | 0.46605915 | 1.64E-121 | 2.33E-117 | PP |
| HELLS | 0.46601815 | 4.38E-67 | 6.20E-63 | PP |
| KNL1 | 0.46260519 | 3.71E-151 | 5.25E-147 | PP |
| HNRNPAB | 0.46065027 | 6.18E-155 | 8.75E-151 | PP |
| PARBP | 0.45915649 | 1.56E-146 | 2.21E-142 | PP |
| CCDC88A | 0.45891303 | 1.36E-102 | 1.93E-98 | PP |
| CCDC34 | 0.45883376 | 1.52E-126 | 2.16E-122 | PP |
| FANCI | 0.45790486 | 1.66E-120 | 2.35E-116 | PP |
| CDKN2D | 0.45745813 | 5.57E-67 | 7.89E-63 | PP |
| DUT | 0.45741964 | 6.10E-73 | 8.64E-69 | PP |
| ARL6IP1 | 0.45596879 | 2.15E-69 | 3.05E-65 | PP |
| KIF2C | 0.45524727 | 5.09E-121 | 7.20E-117 | PP |
| HIST1H1C | 0.45397194 | 1.96E-11 | 2.78E-07 | PP |
| LMNB1 | 0.45017033 | 2.22E-121 | 3.14E-117 | PP |
| ORC6 | 0.44878674 | 5.41E-103 | 7.65E-99 | PP |
| CENPA | 0.44743625 | 5.71E-109 | 8.07E-105 | PP |
| SHCBP1 | 0.44583334 | 3.21E-148 | 4.54E-144 | PP |
| CKS2 | 0.44428741 | 1.86E-80 | 2.63E-76 | PP |
| NUDT1 | 0.44225664 | 4.14E-157 | 5.86E-153 | PP |
| KPNB1 | 0.44218115 | 2.68E-155 | 3.79E-151 | PP |
| CCNA2 | 0.44039524 | 2.07E-107 | 2.93E-103 | PP |
| NUCKS1 | 0.43715445 | 4.07E-219 | 5.75E-215 | PP |
| CENPN | 0.43700514 | 2.02E-147 | 2.86E-143 | PP |
| DNAJC9 | 0.43660548 | 1.85E-115 | 2.61E-111 | PP |
| CENPW | 0.43656706 | 7.54E-142 | 1.07E-137 | PP |
| FAM111A | 0.43618008 | 8.03E-87 | 1.14E-82 | PP |
| FAM111B | 0.43588712 | 6.06E-50 | 8.57E-46 | PP |
| MNS1 | 0.43536873 | 1.09E-97 | 1.54E-93 | PP |
| ESCO2 | 0.43510545 | 5.81E-113 | 8.22E-109 | PP |
| RACGAP1 | 0.43246201 | 6.14E-123 | 8.68E-119 | PP |
| HIST1H2AG | 0.43196269 | 2.74E-78 | 3.87E-74 | PP |
| PSIP1 | 0.43112425 | 2.27E-141 | 3.21E-137 | PP |
| KIF14 | 0.43053557 | 2.16E-103 | 3.06E-99 | PP |
| RPL39L | 0.42999604 | 1.63E-127 | 2.30E-123 | PP |
| TCF19 | 0.42850035 | 1.07E-93 | 1.51E-89 | PP |
| ASF1B | 0.42566651 | 2.37E-106 | 3.36E-102 | PP |
| NUCB2 | 0.42240072 | 2.23E-133 | 3.15E-129 | PP |
| BUB1 | 0.42109417 | 1.71E-119 | 2.42E-115 | PP |
| SPC24 | 0.42015685 | 1.58E-135 | 2.24E-131 | PP |

|  |  |  |  |
| --- | --- | --- | --- |
| HJURP | 0.41936229 | 2.60E-78 | 3.68E-74 PP |
| MND1 | 0.41920401 | 6.75E-136 | 9.55E-132 PP |
| HMGB1 | 0.41843059 | 8.10E-190 | 1.15E-185 PP |
| MCM7 | 0.41744679 | 1.01E-92 | 1.43E-88 PP |
| CDCA2 | 0.41638913 | 4.54E-113 | 6.43E-109 PP |
| JPT1 | 0.41507661 | 3.73E-141 | 5.27E-137 PP |
| NASP | 0.41432054 | 3.09E-109 | 4.37E-105 PP |
| PRKDC | 0.41187051 | 1.21E-96 | 1.71E-92 PP |
| LYAR | 0.41133307 | 7.45E-123 | 1.05E-118 PP |
| FBXO5 | 0.410289 | 4.83E-74 | 6.83E-70 PP |
| KPNA2 | 0.41006538 | 1.09E-83 | 1.54E-79 PP |
| CENPU | 0.4094098 | 2.94E-101 | 4.17E-97 PP |
| CAVIN3 | 0.40621235 | 4.65E-113 | 6.59E-109 PP |
| CKAP5 | 0.40531506 | 1.30E-91 | 1.83E-87 PP |
| PSMD2 | 0.40516101 | 1.34E-147 | 1.90E-143 PP |
| GMNN | 0.40504731 | 5.13E-96 | 7.25E-92 PP |
| CALM3 | 0.40380678 | 3.86E-121 | 5.45E-117 PP |
| ANPEP | 0.40144852 | 1.34E-58 | 1.89E-54 PP |
| ITGB3BP | 0.39822762 | 1.13E-99 | 1.60E-95 PP |
| TPM4 | 0.39754262 | 8.60E-136 | 1.22E-131 PP |
| H2AFX | 0.39729554 | 1.93E-89 | 2.73E-85 PP |
| AXL | 0.39723038 | 1.14E-109 | 1.62E-105 PP |
| NDC80 | 0.39204709 | 7.72E-102 | 1.09E-97 PP |
| SERPINE1 | 0.39169642 | 2.81E-35 | 3.97E-31 PP |
| POC1A | 0.39145683 | 1.15E-123 | 1.62E-119 PP |
| PPM1G | 0.38972662 | 4.52E-146 | 6.40E-142 PP |
| HNRNPR | 0.38891845 | 8.48E-137 | 1.20E-132 PP |
| RANBP1 | 0.38881561 | 1.01E-148 | 1.43E-144 PP |
| PRR11 | 0.38803661 | 1.01E-110 | 1.43E-106 PP |
| HSP90AA1 | 0.3876284 | 2.32E-140 | 3.29E-136 PP |
| SIVA1 | 0.38537248 | 2.96E-132 | 4.19E-128 PP |
| MCM4 | 0.38472527 | 3.72E-61 | 5.26E-57 PP |
| PBX3 | 0.38424295 | 4.10E-93 | 5.80E-89 PP |
| MELK | 0.38133824 | 6.94E-92 | 9.81E-88 PP |
| SMC3 | 0.38118336 | 6.43E-88 | 9.10E-84 PP |
| AHNAK | 0.38090612 | 3.29E-112 | 4.65E-108 PP |
| CCNA1 | 0.38079 | 4.13E-56 | 5.85E-52 PP |
| TPM3 | 0.38010918 | 1.81E-140 | 2.56E-136 PP |
| RFC4 | 0.37897967 | 1.23E-80 | 1.74E-76 PP |
| SMTN | 0.37813024 | 1.15E-79 | 1.63E-75 PP |
| YWHAH | 0.37732887 | 6.78E-128 | 9.60E-124 PP |
| HMGN2 | 0.37580055 | 4.80E-116 | 6.79E-112 PP |
| DCBLD2 | 0.37217443 | 7.01E-83 | 9.91E-79 PP |
| BCL2A1 | 0.37182168 | 4.88E-64 | 6.90E-60 PP |
| CRIP1 | 0.37001743 | 1.94E-44 | 2.75E-40 PP |
| CEP170 | 0.37000255 | 2.91E-50 | 4.12E-46 PP |

|  |  |  |  |  |
| --- | --- | --- | --- | --- |
| CTNNAL1 | 0.36874967 | 1.14E-93 | 1.62E-89 | PP |
| SUPT16H | 0.3682534 | 5.11E-69 | 7.24E-65 | PP |
| SNRPD1 | 0.36686022 | 9.86E-135 | 1.40E-130 | PP |
| AURKA | 0.3666365 | 1.02E-46 | 1.44E-42 | PP |
| CCT5 | 0.36539276 | 1.61E-149 | 2.28E-145 | PP |
| ARHGAP11A | 0.36518259 | 8.94E-110 | 1.27E-105 | PP |
| CDCA4 | 0.36494488 | 3.76E-58 | 5.32E-54 | PP |
| HLA-DRB5 | 0.36476728 | 4.62E-18 | 6.53E-14 | PP |
| TMSB15A | 0.36437325 | 1.74E-88 | 2.46E-84 | PP |
| BRCA1 | 0.36352109 | 6.09E-82 | 8.61E-78 | PP |
| SSRP1 | 0.36279768 | 2.03E-119 | 2.88E-115 | PP |
| CKAP2 | 0.36278923 | 2.59E-40 | 3.66E-36 | PP |
| PCNA | 0.36040356 | 6.63E-42 | 9.38E-38 | PP |
| PALM2-AKAF | 0.36027227 | 1.77E-88 | 2.50E-84 | PP |
| COTL1 | 0.3594299 | 1.44E-97 | 2.03E-93 | PP |
| ARHGAP29 | 0.35827017 | 4.90E-62 | 6.93E-58 | PP |
| TTK | 0.35819354 | 5.70E-89 | 8.06E-85 | PP |
| MYBL2 | 0.3559846 | 9.70E-92 | 1.37E-87 | PP |
| RPA3 | 0.35559276 | 8.14E-94 | 1.15E-89 | PP |
| SMC1A | 0.35502806 | 1.46E-63 | 2.06E-59 | PP |
| SPDL1 | 0.35404502 | 9.91E-63 | 1.40E-58 | PP |
| SAC3D1 | 0.35132861 | 5.60E-86 | 7.93E-82 | PP |
| EZR | 0.35047343 | 4.29E-89 | 6.07E-85 | PP |
| TEX30 | 0.34985236 | 2.86E-91 | 4.04E-87 | PP |
| BUB3 | 0.34771182 | 3.02E-106 | 4.27E-102 | PP |
| MASTL | 0.34768959 | 2.59E-71 | 3.67E-67 | PP |
| KIF20A | 0.34707272 | 3.62E-80 | 5.12E-76 | PP |
| CDC45 | 0.34704561 | 3.02E-69 | 4.28E-65 | PP |
| NRIP3 | 0.34703002 | 5.05E-72 | 7.14E-68 | PP |
| TAGLN2 | 0.3465395 | 4.13E-106 | 5.84E-102 | PP |
| SLBP | 0.34647977 | 1.48E-61 | 2.10E-57 | PP |
| SFPQ | 0.34626319 | 1.90E-95 | 2.69E-91 | PP |
| MZT1 | 0.3462448 | 6.83E-83 | 9.67E-79 | PP |
| SSR3 | 0.34613847 | 3.67E-113 | 5.19E-109 | PP |
| DKC1 | 0.34543113 | 8.67E-95 | 1.23E-90 | PP |
| CDCA5 | 0.34514647 | 1.05E-96 | 1.48E-92 | PP |
| GINS2 | 0.34512953 | 9.01E-66 | 1.27E-61 | PP |
| LRR1 | 0.34440043 | 1.02E-88 | 1.44E-84 | PP |
| CSE1L | 0.3443186 | 1.04E-91 | 1.46E-87 | PP |
| KNSTRN | 0.34343582 | 8.25E-61 | 1.17E-56 | PP |
| CHAF1A | 0.3432243 | 9.67E-62 | 1.37E-57 | PP |
| NT5E | 0.34226377 | 1.10E-89 | 1.56E-85 | PP |
| HSPD1 | 0.34157546 | 1.06E-115 | 1.51E-111 | PP |
| SMS | 0.34137645 | 7.07E-100 | 1.00E-95 | PP |
| PKMYT1 | 0.34118408 | 7.53E-80 | 1.07E-75 | PP |
| NANS | 0.34118322 | 4.07E-111 | 5.76E-107 | PP |

|  |  |  |  |  |
| --- | --- | --- | --- | --- |
| KIF15 | 0.34093524 | 1.55E-109 | 2.20E-105 | PP |
| CENPX | 0.34076559 | 5.04E-74 | 7.13E-70 | PP |
| FLNA | 0.33760655 | 2.99E-92 | 4.23E-88 | PP |
| PTGES3 | 0.33691099 | 6.57E-180 | 9.30E-176 | PP |
| GAS2L3 | 0.33687916 | 6.02E-74 | 8.52E-70 | PP |
| CACYBP | 0.33658358 | 2.14E-113 | 3.03E-109 | PP |
| BAZ1B | 0.33567008 | 3.10E-64 | 4.39E-60 | PP |
| NCAPH | 0.3354431 | 6.41E-98 | 9.07E-94 | PP |
| WDR34 | 0.3354276 | 2.07E-77 | 2.93E-73 | PP |
| PA2G4 | 0.3347911 | 3.78E-133 | 5.35E-129 | PP |
| AC007952.4 | 0.33468853 | 1.32E-50 | 1.87E-46 | PP |
| VRK1 | 0.33456214 | 2.25E-72 | 3.19E-68 | PP |
| MCM3 | 0.33349602 | 3.71E-49 | 5.25E-45 | PP |
| SKA2 | 0.33285574 | 4.67E-93 | 6.60E-89 | PP |
| SRSF2 | 0.33276831 | 4.20E-117 | 5.95E-113 | PP |
| STIP1 | 0.33254154 | 9.60E-102 | 1.36E-97 | PP |
| HNRNPD | 0.33084591 | 2.67E-99 | 3.78E-95 | PP |
| FUS | 0.33081415 | 5.78E-110 | 8.18E-106 | PP |
| MCM10 | 0.33063579 | 4.72E-65 | 6.68E-61 | PP |
| HNRNPU | 0.32990954 | 5.94E-65 | 8.40E-61 | PP |
| RIF1 | 0.32973996 | 7.53E-52 | 1.07E-47 | PP |
| PRIM1 | 0.32871091 | 2.48E-60 | 3.51E-56 | PP |
| ACTG2 | 0.32834066 | 2.38E-56 | 3.37E-52 | PP |
| CBR3 | 0.32798592 | 7.48E-81 | 1.06E-76 | PP |
| STMN1 | 0.32718163 | 2.93E-114 | 4.15E-110 | PP |
| RUVBL2 | 0.32716446 | 1.30E-90 | 1.84E-86 | PP |
| AC253572.2 | 0.32605225 | 1.65E-39 | 2.33E-35 | PP |
| C1QL1 | 0.32599779 | 1.34E-81 | 1.90E-77 | PP |
| CAVIN1 | 0.32561714 | 7.72E-107 | 1.09E-102 | PP |
| MCL1 | 0.32490649 | 7.17E-55 | 1.01E-50 | PP |
| MXD3 | 0.3244242 | 7.07E-82 | 1.00E-77 | PP |
| CDC6 | 0.3241559 | 2.50E-49 | 3.54E-45 | PP |
| ATAD5 | 0.32394461 | 3.52E-66 | 4.99E-62 | PP |
| ALCAM | 0.32363402 | 4.33E-78 | 6.12E-74 | PP |
| LRRFIP2 | 0.32309056 | 4.05E-77 | 5.72E-73 | PP |
| TUBG1 | 0.32263213 | 4.43E-62 | 6.26E-58 | PP |
| POLD3 | 0.32038836 | 7.60E-56 | 1.08E-51 | PP |
| NTNG1 | 0.32031074 | 4.95E-50 | 7.01E-46 | PP |
| LRRC59 | 0.32025595 | 2.44E-90 | 3.45E-86 | PP |
| MANCR | 0.31865352 | 3.20E-66 | 4.53E-62 | PP |
| CCDC18 | 0.31850761 | 8.41E-71 | 1.19E-66 | PP |
| TFPI2 | 0.31829806 | 5.30E-64 | 7.50E-60 | PP |
| YES1 | 0.31825363 | 8.72E-48 | 1.23E-43 | PP |
| PTBP1 | 0.31773551 | 3.62E-86 | 5.13E-82 | PP |
| CENPH | 0.3175589 | 2.34E-81 | 3.31E-77 | PP |
| PARP1 | 0.31734057 | 7.50E-51 | 1.06E-46 | PP |

|  |  |  |  |
| --- | --- | --- | --- |
| PSMC3 | 0.31694199 | 4.62E-100 | 6.54E-96 PP |
| C19orf33 | 0.3165668 | 6.75E-50 | 9.54E-46 PP |
| NT5DC2 | 0.31604512 | 1.13E-79 | 1.59E-75 PP |
| FBL | 0.31601041 | 1.52E-105 | 2.15E-101 PP |
| NOP56 | 0.31578543 | 1.13E-85 | 1.60E-81 PP |
| CDCP1 | 0.315514 | 6.91E-61 | 9.78E-57 PP |
| UQCC3 | 0.31428487 | 7.54E-92 | 1.07E-87 PP |
| LBR | 0.31405809 | 6.36E-64 | 9.01E-60 PP |
| PGP | 0.31383839 | 1.34E-92 | 1.90E-88 PP |
| HLA-DQB1 | 0.31373907 | 2.60E-44 | 3.68E-40 PP |
| RGMB | 0.31355027 | 1.95E-52 | 2.76E-48 PP |
| G0S2 | 0.31348192 | 1.33E-23 | 1.88E-19 PP |
| HNRNPA2B1 | 0.31342484 | 2.81E-135 | 3.97E-131 PP |
| GALNT6 | 0.31255472 | 5.87E-62 | 8.30E-58 PP |
| H2AFV | 0.31247717 | 4.75E-110 | 6.71E-106 PP |
| ANP32B | 0.31173887 | 3.15E-135 | 4.45E-131 PP |
| BUB1B | 0.3116405 | 2.92E-83 | 4.13E-79 PP |
| ARPC5L | 0.3113735 | 1.02E-98 | 1.44E-94 PP |
| MYH9 | 0.3109226 | 2.73E-70 | 3.86E-66 PP |
| PNP | 0.31073851 | 4.25E-67 | 6.02E-63 PP |
| C9orf40 | 0.31064965 | 2.76E-68 | 3.91E-64 PP |
| SRPK1 | 0.31020931 | 2.92E-73 | 4.13E-69 PP |
| SACS | 0.31010748 | 1.51E-54 | 2.13E-50 PP |
| LGALS1 | 0.30999977 | 1.34E-104 | 1.90E-100 PP |
| SKA3 | 0.3096681 | 2.93E-111 | 4.14E-107 PP |
| IKBIP | 0.30895865 | 1.39E-74 | 1.96E-70 PP |
| EZH2 | 0.3089079 | 3.08E-64 | 4.35E-60 PP |
| UHRF1 | 0.30835751 | 2.34E-67 | 3.31E-63 PP |
| CDT1 | 0.30818282 | 1.90E-65 | 2.69E-61 PP |
| BARD1 | 0.30763525 | 6.26E-55 | 8.86E-51 PP |
| HNRNPA3 | 0.30749623 | 1.10E-114 | 1.55E-110 PP |
| SRRT | 0.30720596 | 9.68E-68 | 1.37E-63 PP |
| ILF2 | 0.30661117 | 3.39E-111 | 4.80E-107 PP |
| ADTRP | 0.30523366 | 2.72E-46 | 3.85E-42 PP |
| RBM25 | 0.30522673 | 3.50E-66 | 4.95E-62 PP |
| CAV1 | 0.30514586 | 2.02E-93 | 2.86E-89 PP |
| EXOSC8 | 0.30432379 | 1.29E-69 | 1.82E-65 PP |
| NCL | 0.30432082 | 1.76E-44 | 2.48E-40 PP |
| QSER1 | 0.30406434 | 1.66E-63 | 2.35E-59 PP |
| TEDC1 | 0.30393675 | 1.96E-79 | 2.77E-75 PP |
| SRSF7 | 0.30240173 | 1.40E-87 | 1.99E-83 PP |
| BCL2L12 | 0.30146787 | 2.50E-64 | 3.54E-60 PP |
| RHEB | 0.30122669 | 2.76E-101 | 3.90E-97 PP |
| PEX5L | 0.30088185 | 1.10E-39 | 1.56E-35 PP |
| PPP1CA | 0.30078 | 5.56E-106 | 7.86E-102 PP |
| C16orf95 | 0.29982616 | 1.61E-53 | 2.28E-49 PP |

|  |  |  |  |
| --- | --- | --- | --- |
| ECI2 | 0.29961704 | 1.78E-77 | 2.52E-73 PP |
| TOMM40 | 0.29928698 | 1.57E-86 | 2.22E-82 PP |
| EIF4EBP1 | 0.29890386 | 2.54E-66 | 3.60E-62 PP |
| MAGOHB | 0.29860619 | 6.65E-79 | 9.41E-75 PP |
| C2orf88 | 0.29846827 | 9.82E-69 | 1.39E-64 PP |
| DRAP1 | 0.29830633 | 4.72E-102 | 6.69E-98 PP |
| HNRNPM | 0.29669945 | 5.96E-103 | 8.44E-99 PP |
| MSN | 0.29668969 | 3.81E-62 | 5.40E-58 PP |
| SNRPB | 0.2966604 | 3.49E-112 | 4.94E-108 PP |
| SPAG5 | 0.2965941 | 1.07E-98 | 1.51E-94 PP |
| TRIP13 | 0.29649432 | 2.01E-87 | 2.85E-83 PP |
| RTKN2 | 0.29638045 | 1.46E-66 | 2.07E-62 PP |
| RAB3B | 0.29507708 | 1.86E-58 | 2.63E-54 PP |
| PAFAH1B3 | 0.29493194 | 1.04E-48 | 1.47E-44 PP |
| DMBT1 | 0.29429503 | 6.85E-51 | 9.69E-47 PP |
| FGF5 | 0.29428422 | 1.14E-59 | 1.62E-55 PP |
| ACTL6A | 0.29405685 | 2.53E-85 | 3.59E-81 PP |
| GNPNAT1 | 0.2939467 | 8.56E-56 | 1.21E-51 PP |
| FARP1 | 0.29353961 | 8.16E-68 | 1.15E-63 PP |
| BZW1 | 0.2928473 | 3.02E-85 | 4.27E-81 PP |
| MPP6 | 0.29252599 | 4.96E-69 | 7.01E-65 PP |
| KMT5A | 0.29151558 | 9.46E-50 | 1.34E-45 PP |
| TJP1 | 0.29128559 | 5.80E-48 | 8.21E-44 PP |
| SVIP | 0.29001414 | 1.21E-56 | 1.71E-52 PP |
| C3orf14 | 0.2892108 | 1.17E-65 | 1.66E-61 PP |
| E2F1 | 0.28906272 | 1.10E-40 | 1.56E-36 PP |
| RFC2 | 0.28872191 | 1.49E-43 | 2.11E-39 PP |
| CALU | 0.28782522 | 2.73E-88 | 3.86E-84 PP |
| RECQL | 0.28772359 | 3.44E-57 | 4.87E-53 PP |
| WDHD1 | 0.28707559 | 3.42E-63 | 4.84E-59 PP |
| CLEC11A | 0.2859337 | 6.75E-70 | 9.55E-66 PP |
| NCAPG2 | 0.28389662 | 4.27E-87 | 6.04E-83 PP |
| CBX5 | 0.2826149 | 1.41E-66 | 1.99E-62 PP |
| HP1BP3 | 0.28256002 | 2.60E-46 | 3.68E-42 PP |
| PIMREG | 0.28204742 | 2.27E-88 | 3.22E-84 PP |
| SYNCRIP | 0.28134241 | 2.29E-73 | 3.24E-69 PP |
| VCAN | 0.28129645 | 7.91E-69 | 1.12E-64 PP |
| UAP1 | 0.2807903 | 8.23E-64 | 1.16E-59 PP |
| DBF4 | 0.28016059 | 3.78E-44 | 5.36E-40 PP |
| BRCA2 | 0.27996975 | 6.86E-65 | 9.70E-61 PP |
| HIRIP3 | 0.27979402 | 6.06E-57 | 8.57E-53 PP |
| KIF18A | 0.2796718 | 6.76E-54 | 9.57E-50 PP |
| GMPS | 0.27952743 | 3.51E-53 | 4.96E-49 PP |
| SAE1 | 0.27950109 | 6.78E-68 | 9.60E-64 PP |
| CCT2 | 0.27820059 | 4.92E-101 | 6.97E-97 PP |
| NUDC | 0.27810012 | 1.11E-97 | 1.57E-93 PP |

|  |  |  |  |
| --- | --- | --- | --- |
| H3F3B | 0.27754758 | 2.55E-127 | 3.61E-123 PP |
| NUDT5 | 0.27673846 | 5.06E-51 | 7.17E-47 PP |
| EIF5 | 0.2761412 | 6.36E-101 | 9.00E-97 PP |
| CEP78 | 0.27600015 | 5.85E-50 | 8.28E-46 PP |
| DHCR24 | 0.27561026 | 3.79E-50 | 5.36E-46 PP |
| PPP1R14A | 0.27545085 | 4.30E-72 | 6.09E-68 PP |
| TPRKB | 0.27418698 | 1.17E-81 | 1.66E-77 PP |
| ROCK2 | 0.27354031 | 4.89E-49 | 6.92E-45 PP |
| RAN | 0.27321138 | 2.11E-126 | 2.99E-122 PP |
| STEAP1 | 0.27295606 | 5.83E-56 | 8.24E-52 PP |
| DEPDC1B | 0.27292029 | 2.34E-80 | 3.31E-76 PP |
| LPXN | 0.27289047 | 5.37E-58 | 7.60E-54 PP |
| CMC2 | 0.27246741 | 4.91E-52 | 6.95E-48 PP |
| PNN | 0.27236519 | 8.10E-52 | 1.15E-47 PP |
| EBNA1BP2 | 0.27205084 | 2.45E-69 | 3.47E-65 PP |
| ATP2B4 | 0.27194618 | 1.23E-51 | 1.74E-47 PP |
| MARCKS | 0.27193857 | 1.01E-41 | 1.43E-37 PP |
| DLC1 | 0.27166706 | 1.80E-48 | 2.55E-44 PP |
| FAM83D | 0.27150418 | 8.30E-62 | 1.17E-57 PP |
| CHEK1 | 0.27130136 | 4.89E-59 | 6.92E-55 PP |
| MAPKAP1 | 0.27117311 | 2.58E-58 | 3.65E-54 PP |
| SDC1 | 0.27111876 | 7.94E-41 | 1.12E-36 PP |
| SNRPG | 0.26993364 | 1.27E-94 | 1.80E-90 PP |
| NOP58 | 0.26989189 | 3.48E-64 | 4.92E-60 PP |
| BLM | 0.26986119 | 7.23E-66 | 1.02E-61 PP |
| ACTN4 | 0.26953862 | 3.16E-79 | 4.46E-75 PP |
| MLEC | 0.26888376 | 1.33E-73 | 1.88E-69 PP |
| ILF3 | 0.26872298 | 5.62E-63 | 7.95E-59 PP |
| EXO1 | 0.26832397 | 1.48E-64 | 2.09E-60 PP |
| ZEB1 | 0.26809605 | 9.03E-45 | 1.28E-40 PP |
| HAT1 | 0.26705885 | 5.51E-50 | 7.80E-46 PP |
| CNIH4 | 0.26705054 | 3.26E-64 | 4.61E-60 PP |
| CRNDE | 0.26697998 | 8.68E-56 | 1.23E-51 PP |
| MIS18A | 0.26696959 | 1.51E-56 | 2.13E-52 PP |
| C11orf24 | 0.26670859 | 1.46E-59 | 2.07E-55 PP |
| ARPC5 | 0.26649882 | 2.87E-95 | 4.05E-91 PP |
| HSP90B1 | 0.26508543 | 1.03E-71 | 1.46E-67 PP |
| CDCA8 | 0.26505734 | 1.01E-58 | 1.43E-54 PP |
| ARHGDIB | 0.26459425 | 2.04E-56 | 2.89E-52 PP |
| COMMD4 | 0.26455212 | 2.30E-67 | 3.26E-63 PP |
| SMURF2 | 0.26424586 | 1.48E-37 | 2.09E-33 PP |
| TFDP1 | 0.2642124 | 5.06E-56 | 7.16E-52 PP |
| PRSS23 | 0.26338271 | 7.73E-48 | 1.09E-43 PP |
| HAS2 | 0.26337679 | 1.81E-27 | 2.56E-23 PP |
| TOPBP1 | 0.26328552 | 2.26E-55 | 3.19E-51 PP |
| POP7 | 0.26308785 | 3.55E-72 | 5.03E-68 PP |

|  |  |  |  |
| --- | --- | --- | --- |
| NUP50 | 0.26278429 | 1.09E-53 | 1.54E-49 PP |
| NMU | 0.26274724 | 1.73E-60 | 2.45E-56 PP |
| GSPT1 | 0.26264207 | 4.45E-53 | 6.30E-49 PP |
| SF3B2 | 0.26219841 | 1.02E-65 | 1.45E-61 PP |
| TPR | 0.26204847 | 2.12E-47 | 3.00E-43 PP |
| DTL | 0.2616276 | 1.17E-44 | 1.66E-40 PP |
| EIF4G1 | 0.26135333 | 1.34E-45 | 1.89E-41 PP |
| S100A10 | 0.26130767 | 2.20E-82 | 3.11E-78 PP |
| CTSC | 0.25992874 | 1.71E-65 | 2.43E-61 PP |
| EIF1AX | 0.25992453 | 8.55E-90 | 1.21E-85 PP |
| PRKAG2 | 0.25924599 | 3.45E-51 | 4.88E-47 PP |
| HACD3 | 0.25895153 | 1.21E-65 | 1.71E-61 PP |
| ALYREF | 0.25887733 | 1.60E-52 | 2.27E-48 PP |
| ZWILCH | 0.25860985 | 6.94E-68 | 9.82E-64 PP |
| CLIC1 | 0.25848774 | 4.76E-88 | 6.73E-84 PP |
| LOXL1-AS1 | 0.25720872 | 1.13E-37 | 1.60E-33 PP |
| FANCD2 | 0.25708192 | 1.07E-81 | 1.51E-77 PP |
| G3BP1 | 0.25674916 | 1.38E-60 | 1.95E-56 PP |
| PITPNC1 | 0.2566117 | 3.50E-49 | 4.96E-45 PP |
| MT-ND6 | 0.25651707 | 5.57E-30 | 7.88E-26 PP |
| MTHFD1 | 0.25644747 | 3.61E-35 | 5.10E-31 PP |
| NOLC1 | 0.25609223 | 1.58E-41 | 2.24E-37 PP |
| DLEU2 | 0.25579641 | 2.79E-67 | 3.95E-63 PP |
| HSPB11 | 0.25555752 | 1.08E-43 | 1.53E-39 PP |
| NCAPH2 | 0.25465056 | 1.40E-50 | 1.98E-46 PP |
| XRCC6 | 0.25463376 | 2.83E-83 | 4.00E-79 PP |
| TCERG1 | 0.25401414 | 2.28E-48 | 3.23E-44 PP |
| KHDRBS1 | 0.25388419 | 2.95E-71 | 4.17E-67 PP |
| GLIPR1 | 0.25295942 | 3.03E-55 | 4.29E-51 PP |
| TCOF1 | 0.25269662 | 3.40E-50 | 4.81E-46 PP |
| BNC1 | 0.25254133 | 7.65E-43 | 1.08E-38 PP |
| CKB | 0.25251707 | 9.45E-38 | 1.34E-33 PP |
| XRCC5 | 0.25162855 | 6.06E-62 | 8.57E-58 PP |
| ILK | 0.25153742 | 1.42E-50 | 2.00E-46 PP |
| PXMP2 | 0.25125569 | 3.68E-35 | 5.21E-31 PP |
| ANTXR2 | 0.25122901 | 3.23E-45 | 4.56E-41 PP |
| EIF2S1 | 0.2510326 | 3.03E-63 | 4.28E-59 PP |
| RASA1 | 0.25101066 | 5.08E-48 | 7.19E-44 PP |
| TNFRSF12A | 0.25068875 | 3.15E-63 | 4.45E-59 PP |
| HIST2H2AC | 0.25059735 | 4.84E-09 | 6.84E-05 PP |
